## Supplementary figures and images for "The *Plasmodium* NOT1-G Paralogue Acts as an Essential Nexus for Sexual Stage Maturation and Parasite Transmission"

### Supp Figures 1-4

Hart *et al.* Figure S1

A.

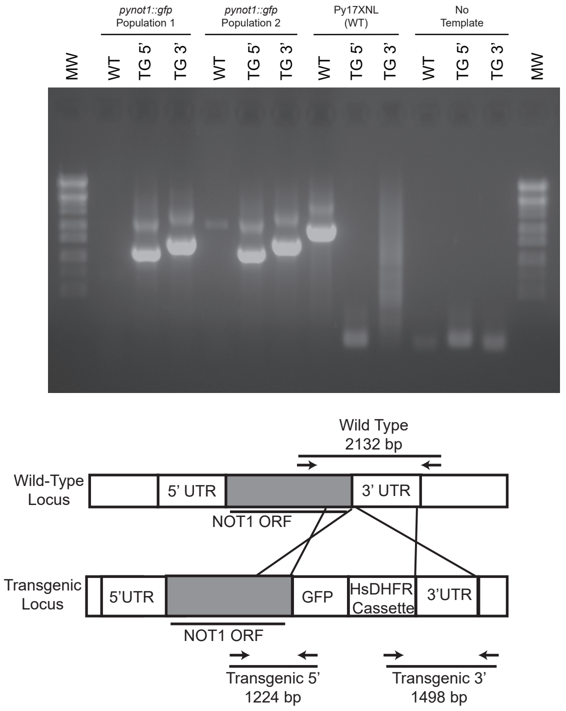


B.

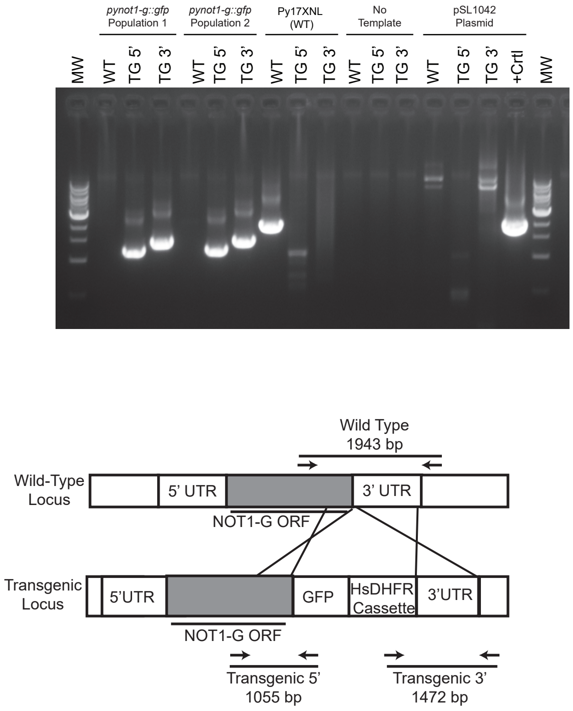

#### Hart *et al.* Figure S2

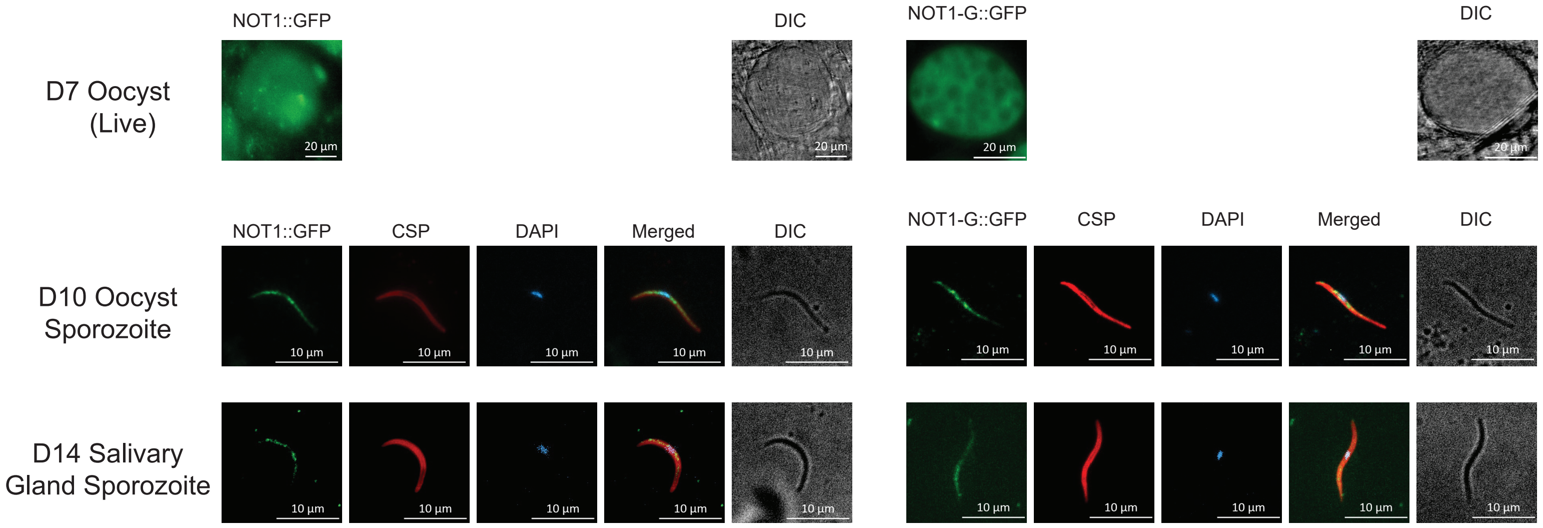

#### Hart *et al.* Figure S3

A.

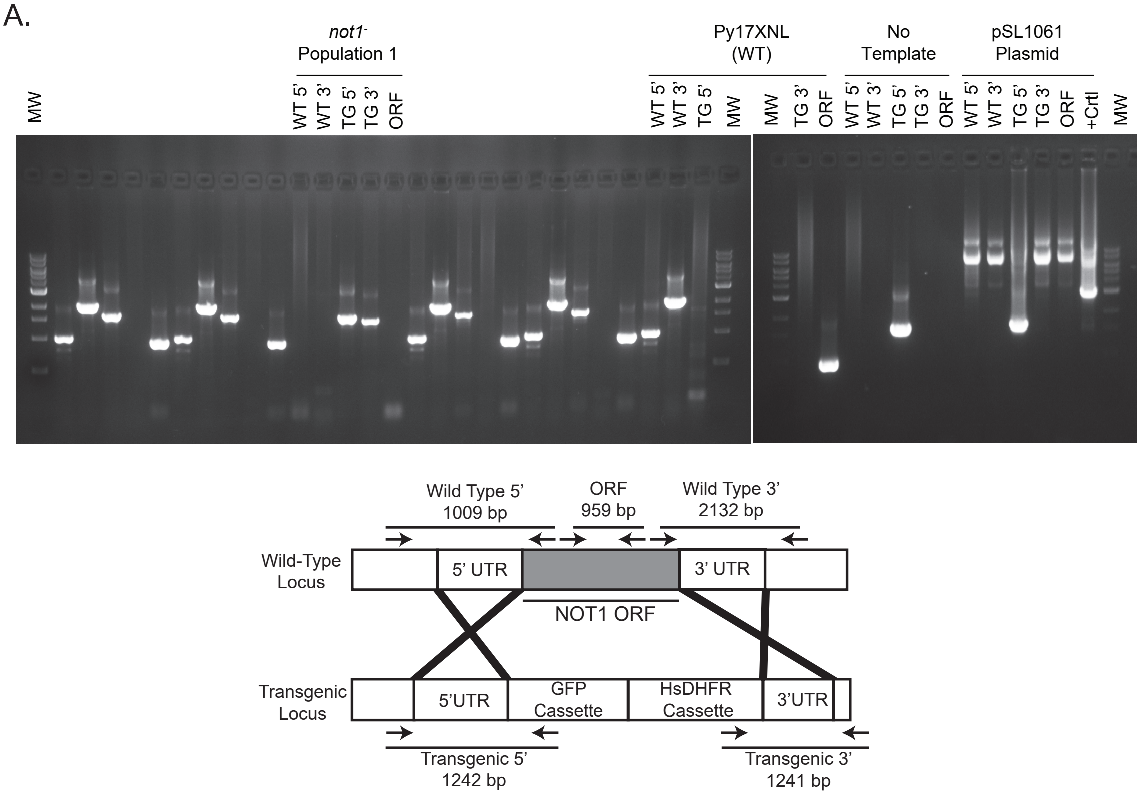


B.

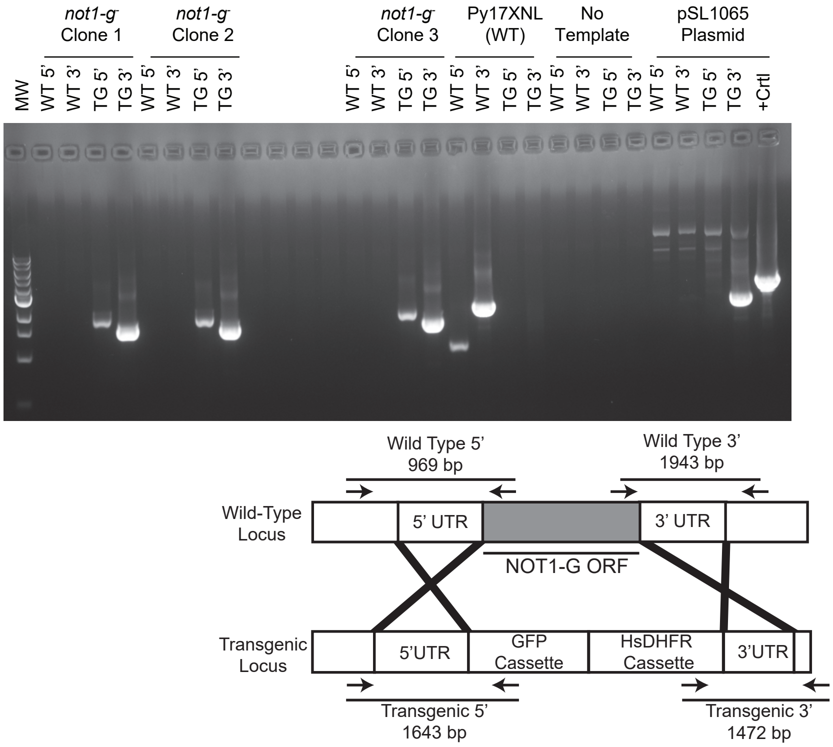

#### Hart *et al.* Figure S4

A.

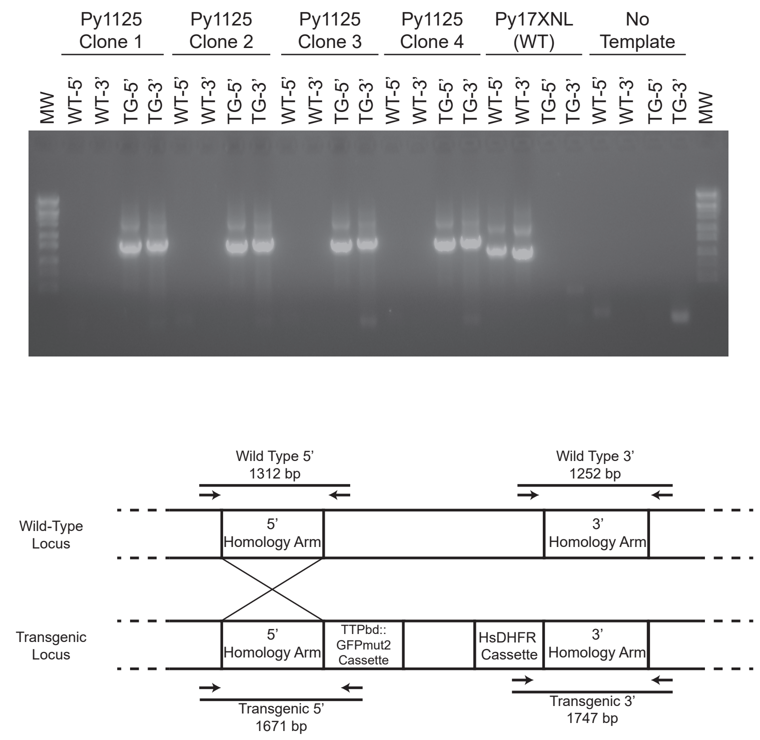


B.

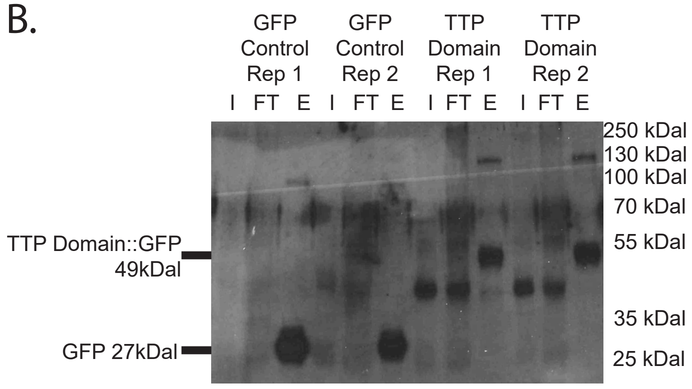


C.

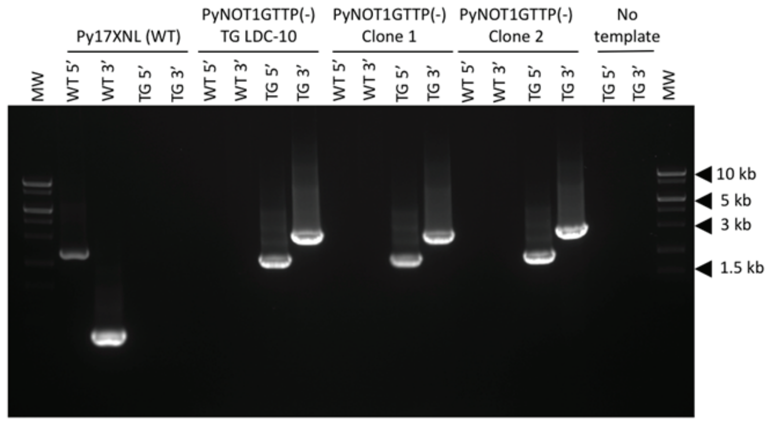


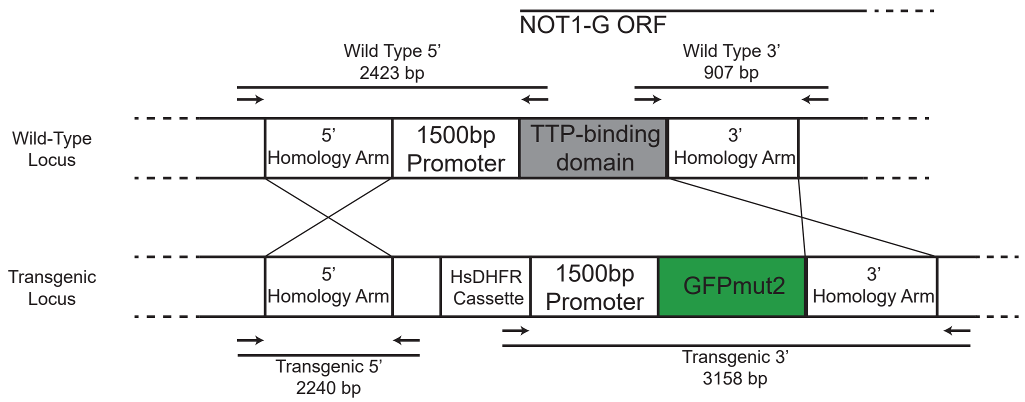
