## Supplementary material for "The *Plasmodium* NOT1-G Paralogue Acts as an Essential Nexus for Sexual Stage Maturation and Parasite Transmission": Supp File 1 - Complete Plasmid Files

**Hart et al. Supplemental File 1 - Complete Plasmid Sequences**

```
>Py0489_PyWT-GFPmut2
LOCUS       pSL0489__final_         7758 bp    DNA        circular     21-APR-2021
SOURCE
  ORGANISM
COMMENT     Lindner Lab - used to create Py0489 = Py17XNL WT-GFP
COMMENT     ApEinfo:methylated:1
FEATURES             Location/Qualifiers
     exon            6735..7298
                     /vntifkey="61"
                     /locus_tag="hDHFR"
                     /label="hDHFR"
                     /ApEinfo_label="hDHFR"
                     /ApEinfo_fwdcolor="pink"
                     /ApEinfo_revcolor="pink"
                     /ApEinfo_graphicformat="arrow_data {{0 0.5 0 1 2 0 0 -1 0
                     -0.5} {} 0} width 5 offset 0"
     misc_feature    1898..2471
                     /locus_tag="PbEF1a promoter"
                     /label="PbEF1a promoter"
                     /ApEinfo_label="PbEF1a promoter"
                     /ApEinfo_fwdcolor="cyan"
                     /ApEinfo_revcolor="green"
                     /ApEinfo_graphicformat="arrow_data {{0 0.5 0 1 2 0 0 -1 0
                     -0.5} {} 0} width 5 offset 0"
     misc_feature    2516..3194
                     /locus_tag="GFPmut2"
                     /label="GFPmut2"
                     /ApEinfo_label="GFPmut2"
                     /ApEinfo_fwdcolor="cyan"
                     /ApEinfo_revcolor="green"
                     /ApEinfo_graphicformat="arrow_data {{0 0.5 0 1 2 0 0 -1 0
                     -0.5} {} 0} width 5 offset 0"
     misc_feature    3201..3648
                     /locus_tag="PbDHFR/TS\3'UTR"
                     /label="PbDHFR/TS\3'UTR"
                     /ApEinfo_label="PbDHFR/TS\3'UTR"
                     /ApEinfo_fwdcolor="#ff0000"
                     /ApEinfo_revcolor="green"
                     /ApEinfo_graphicformat="arrow_data {{0 0.5 0 1 2 0 0 -1 0
                     -0.5} {} 0} width 5 offset 0"
     rep_origin      5045..5727
                     /locus_tag="ColE1 origin"
                     /label="ColE1 origin"
                     /ApEinfo_label="ColE1 origin"
                     /ApEinfo_fwdcolor="gray50"
                     /ApEinfo_revcolor="gray50"
                     /ApEinfo_graphicformat="arrow_data {{0 0.5 0 1 2 0 0 -1 0
                     -0.5} {} 0} width 5 offset 0"
     CDS             4288..4947
                     /locus_tag="AmpR"
                     /label="AmpR"
                     /ApEinfo_label="AmpR"
                     /ApEinfo_fwdcolor="yellow"
                     /ApEinfo_revcolor="yellow"
                     /ApEinfo_graphicformat="arrow_data {{0 0.5 0 1 2 0 0 -1 0
                     -0.5} {} 0} width 5 offset 0"
     misc_feature    7305..7755
                     /locus_tag="PbDHFR-TS 3'UTR"
                     /label="PbDHFR-TS 3'UTR"
                     /ApEinfo_label="PbDHFR-TS 3'UTR"
                     /ApEinfo_fwdcolor="cyan"
                     /ApEinfo_revcolor="green"
                     /ApEinfo_graphicformat="arrow_data {{0 0.5 0 1 2 0 0 -1 0
                     -0.5} {} 0} width 5 offset 0"
     misc_feature    join(6131..6440,6445..6729)
                     /locus_tag="PbEF1a 5'UTR"
                     /label="PbEF1a 5'UTR"
                     /ApEinfo_label="PbEF1a 5'UTR"
                     /ApEinfo_fwdcolor="#804040"
                     /ApEinfo_revcolor="#804040"
                     /ApEinfo_graphicformat="arrow_data {{0 0.5 0 1 2 0 0 -1 0
                     -0.5} {} 0} width 5 offset 0"
     misc_feature    7..918
                     /locus_tag="py p230p Homology Arm 1"
                     /label="py p230p Homology Arm 1"
                     /ApEinfo_label="py p230p Homology Arm 1"
                     /ApEinfo_fwdcolor="cyan"
                     /ApEinfo_revcolor="green"
                     /ApEinfo_graphicformat="arrow_data {{0 0.5 0 1 2 0 0 -1 0
                     -0.5} {} 0} width 5 offset 0"
     misc_feature    925..1873
                     /locus_tag="py p230p Homology Arm 2"
                     /label="py p230p Homology Arm 2"
                     /ApEinfo_label="py p230p Homology Arm 2"
                     /ApEinfo_fwdcolor="cyan"
                     /ApEinfo_revcolor="green"
                     /ApEinfo_graphicformat="arrow_data {{0 0.5 0 1 2 0 0 -1 0
                     -0.5} {} 0} width 5 offset 0"
ORIGIN
        1 ggtaccAGCT TTGTGTTTTA TTTGGATGTG CAACAATAGA TTTATGATTA ACCAATGGTA
       61 AATCATTATA ATCATATATT TCATCTTCAT TTACATATTT TACTAAAACA GTTTTTACAA
      121 TTTGTTCTGT TTCTTTATCT ACACAATGAC ATTCAAATTG ATATGATTTT GTTAATGTAT
      181 AATTAAAACC AGCTAATGCA TATTCTGGAT TTTCAAATGA ATAGTTAATA ATATTTTTAG
      241 TATTAATTTT ATAGGAATCA TGTAGATAGC CTTTTTCATA TACTTGAGTA AAACATTTTT
      301 CTGGCATTTT TTTTGTATTT TCGGGGCATC TAAAACCAAA ACTTGTATTT GATTTGGAAA
      361 TTATTATACA TAGATCTCCT TCACTATAAG CATTTAAATA ATTATATAAT TCACCTTTTC
      421 TAAAATCACA AAATGAAACT GGTCTATCTT TTAAATCTGT TCGAACATGT AATAAAATAA
      481 TTGCTCTTCT ATAATCAGTC CCACCATTTG TTACACCACA TTCTATGGTT AGTGGATATG
      541 ATCCATGATT TATAAATGAG GTCATAAAAA ATTTAAAACC TTTAAATTGT TCATTTATTT
      601 CTGGTATATC TATATCTTTT GTATTAAAAA ATAAATCTGT AAATTTATTT ATAGGAAATA
      661 CATTACTTAA ATATTTTATG GTATTAATAG GTTCTTTACC ATATACGATT CTATTATCTG
      721 TTATTTGCTC ACAATTATAC ATGAATAAAC TAAATTCATT TATTGTTTTT TCACATATTT
      781 GTATTTCACT TAATTTGCTT AAATTATAAT CAAAATTACA CATCTGATAA GGTATATCGT
      841 TAAAATATTT AATCGTTTTA CTATCATCAT CAACTATTTT TAAAATGTGA TCAGATGTTG
      901 CCATTCCATC ACCAAATGgg gcccTATGGA ACTACATCTA TATAAGAGAT TTTTTTATTT
      961 TTATATATTG GTTTTGAATC TAGATATTCA TAATTTTTAA TATATATTGT TCCTATATTT
     1021 TTGGTTTTAT AATCTTCACA TGTACAATAA AAATCAAATT CTTTTTTTGT AATTTTTGAA
     1081 ATCATTAAAA ATGTTGATGA ATTATTTATA AAATAATCTG CATCTATAAC TTTTATTTCA
     1141 TTAAATATAT TTTCGATATT TTCAGTTATT TGTTCATTCA TGTTACTTTT TATATATACC
     1201 TTTCTAAAAC AATTTTTTGG ATTAATTTTT CCAGTTGGAC AATTTAAGCT AACTACTATA
     1261 TTTTCTTTTA ATTCGATTTC ACAAAATTCA TTTTGTTTTT CTAAATCATA AACATTGGAA
     1321 TAGTAAAATT TTTTTTTAGA ATTACCTGAA AAATCACATC CATATAATTT ATTTTTTGTT
     1381 GTATTTTGAG AAACCCTTAT CACTGTAATA CCTTTTTGAC CTTTCATATT ATTAATTATT
     1441 GTTTTAGTAT TATCACAATA TATTTTCAAT ATTAATACTT TTTCTACAAT TAATGGAGTA
     1501 ACAAAAGTGA AATTTCCTTT CTCTTTGAAC ATTTTATTTT TATCTAATAT GATAACACCT
     1561 GGTAATATAT CATCTAATTT TTCTTTTTTG TCTAGATACA TTACTTCATC ATTTTCTGGT
     1621 ATGTATTCTA TATTTTTATA TTTTTCTAAT TCTTCTGGTT CAACATTTGC ATTTTCTAAA
     1681 TAAAAATTAT TTTTTTGACT ACTACGTTGT GATGTTTTCT TATTATTTAT TATGATATTA
     1741 CTATCAAAAA TGTCATCATC ACTTTTTACA ATTTTGGGAC ATTTAACTTG AACCATATTC
     1801 AGATTCTGTT CTAGATGTAT ATAACATGAA TGAACTGTAT CTACTTCATT AACGGGTTCA
     1861 AGAGAATTCG GTAgccgcgg tggcggccgc TctcgagAGC TTAATTCTTT TCGAGCTCTT
     1921 TATGCTTAAG TTTACAATTT AATATTCATA CTTTAAGTAT TTTTTGTAGT ATCCTAGATA
     1981 TTGTGCTTTA AATGCTCACC CCTCAAAGCA CCAGTAATAT TTTCATCCAC TGAAATACCA
     2041 TTAAATTTTC AAAAAAATAC TATGCATATA ATGTTATACA TATAAACATA AAACGCCATG
     2101 TAAATCAAAA AATATATAAA AATATGTATA AAAATAAATA TGCACTAAAT ATAAGCTAAT
     2161 TATGCATAAA AATTAAAGTG CCCTTTATTA ACTAGctagT CGTAATTATT TATATTTCTA
     2221 TGTTATAAAA AAATCCTCAT ATAATAATAT AATTAATATA TGTAATGTTT TTTTTATTTT
     2281 ATAATTTTAA TATAAAATAA TATGTAAATT AATTCAAAAA ATAAATATAA TTGTTGTGAA
     2341 ACAAAAAACG TAATTTTTTC ATTTGCCTTC AAAATTTAAA TTTATTTTAA TATTTCCTAA
     2401 AATATATATA CTTTGTGTAT AAATATATAA AAATATATAT TTGCTTATAA ATAAATAAAA
     2461 ATTTTATAAA Aatgactagt agtaaaggag aagaactttt cactggagtt gtcccAATTC
     2521 TTGTTGAATT AGATGGTGAT GTTAATGGGC ACAAATTTTC TGTCAGTGGA GAGGGTGAAG
     2581 GTGATGCAAC ATACGGAAAA CTTACCCTTA AATTTATTTG CACTACTGGA AAACTACCTG
     2641 TTCCATGGCC AACACTTGTC ACTACTTTCG CGTATGGTCT TCAATGCTTT GCGAGATACC
     2701 CAGATCATAT GAAACAGCAT GACTTTTTCA AGAGTGCCAT GCCCGAAGGT TATGTACAGG
     2761 AAAGAACTAT ATTTTTCAAA GATGACGGGA ACTACAAGAC ACGTGCTGAA GTCAAGTTTG
     2821 AAGGTGATAC CCTTGTTAAT AGAATCGAGT TAAAAGGTAT TGATTTTAAA GAAGATGGAA
     2881 ACATTCTTGG ACACAAATTG GAATACAACT ATAACTCACA CAATGTATAC ATCATGGCAG
     2941 ACAAACAAAA GAATGGAATC AAAGTTAACT TCAAAATTAG ACACAACATT GAAGATGGAA
     3001 GCGTTCAACT AGCAGACCAT TATCAACAAA ATACTCCAAT TGGCGATGGC CCTGTCCTTT
     3061 TACCAGACAA CCATTACCTG TCCACACAAT CTGCCCTTTC GAAAGATCCC AACGAAAAGA
     3121 GAGACCACAT GGTCCTTCTT GAGTTTGTAA CAGCTGCTGG GATTACACAT GGCATGGATG
     3181 AACTATACAA ATAAggatcc GTTTTTCTTA CTTATATATT TATACCAATT GATTGTATTT
     3241 ATAACTGTAA AAATGTGTAT GTTGTGTGCA TATTTTTTTT TGTGCATGCA CATGCATGTA
     3301 AATAGCTAAA ATTATGAACA TTTTATTTTT TGTTCAGAAA AAAAAAACTT TACACACATA
     3361 AAATGGCTAG TATGAATAGC CATATTTTAT ATAAATTAAA TCCTATGAAT TTATGACCAT
     3421 ATTAAAAATT TAGATATTTA TGGAACATAA TATGTTTGAA ACAATAAGAC AAAATTATTA
     3481 TTATTATTAT TATTTTTACT GTTATAATTA TGTTGTCTCT TCAATGATTC ATAAATAGTT
     3541 GGACTTGATT TTTAAAATGT TTATAATATG ATTAGCATAG TTAAATAAAA AAAGTTGAAA
     3601 AATTAAAAAA AAACATATAA ACACAAATGA TGTTTTTTCC TTCAATTTcg gcgcctgatg
     3661 cggtattttc tccttacgca tctgtgcggt atttcacacc gcatatggtg cactctcagt
     3721 acaatctgct ctgatgccgc atagttaagc cagccccgac acccgccaac acccgctgac
     3781 gcgccctgac gggcttgtct gctcccggca tccgcttaca gacaagctgt gaccgtctcc
     3841 gggagctgca tgtgtcagag gttttcaccg tcatcaccga aacgcgcgag acgaaagggc
     3901 ctcgtgatac gcctattttt ataggttaat gtcatgataa taatggtttc ttagacgtca
     3961 ggtggcactt ttcggggaaa tgtgcgcgga acccctattt gtttattttt ctaaatacat
     4021 tcaaatatgt atccgctcat gagacaataa ccctgataaa tgcttcaata atattgaaaa
     4081 aggaagagta tgagtattca acatttccgt gtcgccctta ttcccttttt tgcggcattt
     4141 tgccttcctg tttttgctca cccagaaacg ctggtgaaag taaaagatgc tgaagatcag
     4201 ttgggtgcac gagtgggtta catcgaactg gatctcaaca gcggtaagat ccttgagagt
     4261 tttcgccccg aagaacgttt tccaatgatg agcactttta aagttctgct atgtggcgcg
     4321 gtattatccc gtattgacgc cgggcaagag caactcggtc gccgcataca ctattctcag
     4381 aatgacttgg ttgagtactc accagtcaca gaaaagcatc ttacggatgg catgacagta
     4441 agagaattat gcagtgctgc cataaccatg agtgataaca ctgcggccaa cttacttctg
     4501 acaacgatcg gaggaccgaa ggagctaacc gcttttttgc acaacatggg ggatcatgta
     4561 actcgccttg atcgttggga accggagctg aatgaagcca taccaaacga cgagcgtgac
     4621 accacgatgc ctgtagcaat ggcaacaacg ttgcgcaaac tattaactgg cgaactactt
     4681 actctagctt cccggcaaca attaatagac tggatggagg cggataaagt tgcaggacca
     4741 cttctgcgct cggcccttcc ggctggctgg tttattgctg ataaatctgg agccggtgag
     4801 cgtgggtctc gcggtatcat tgcagcactg gggccagatg gtaagccctc ccgtatcgta
     4861 gttatctaca cgacggggag tcaggcaact atggatgaac gaaatagaca gatcgctgag
     4921 ataggtgcct cactgattaa gcattggtaa ctgtcagacc aagtttactc atatatactt
     4981 tagattgatt taaaacttca tttttaattt aaaaggatct aggtgaagat cctttttgat
     5041 aatctcatga ccaaaatccc ttaacgtgag ttttcgttcc actgagcgtc agaccccgta
     5101 gaaaagatca aaggatcttc ttgagatcct ttttttctgc gcgtaatctg ctgcttgcaa
     5161 acaaaaaaac caccgctacc agcggtggtt tgtttgccgg atcaagagct accaactctt
     5221 tttccgaagg taactggctt cagcagagcg cagataccaa atactgtcct tctagtgtag
     5281 ccgtagttag gccaccactt caagaactct gtagcaccgc ctacatacct cgctctgcta
     5341 atcctgttac cagtggctgc tgccagtggc gataagtcgt gtcttaccgg gttggactca
     5401 agacgatagt taccggataa ggcgcagcgg tcgggctgaa cggggggttc gtgcacacag
     5461 cccagcttgg agcgaacgac ctacaccgaa ctgagatacc tacagcgtga gcattgagaa
     5521 agcgccacgc ttcccgaagg gagaaaggcg gacaggtatc cggtaagcgg cagggtcgga
     5581 acaggagagc gcacgaggga gcttccaggg ggaaacgcct ggtatcttta tagtcctgtc
     5641 gggtttcgcc acctctgact tgagcgtcga tttttgtgat gctcgtcagg ggggcggagc
     5701 ctatggaaaa acgccagcaa cgcggccttt ttacggttcc tggccttttg ctggcctttt
     5761 gctcacatgt tctttcctgc gttatcccct gattctgtgg ataaccgtat taccgccttt
     5821 gagtgagctg ataccgctcg ccgcagccga acgaccgagc gcagcgagtc agtgagcgag
     5881 gaagcggaag agcgcccaat acgcaaaccg cctctccccg cgcgttggcc gattcattaa
     5941 tgcagctggc acgacaggtt tcccgactgg aaagcgggca gtgagcgcaa cgcaattaat
     6001 gtgagttagc tcactcatta ggcaccccag gctttacact ttatgcttcc ggctcgtatg
     6061 ttgtgtggaa ttgtgagcgg ataacaattt cacacaggaa acagctatga ccatgattac
     6121 gccaagcttg ataattcctg cagcccagct taattctttt cgagctcttt atgcttaagt
     6181 ttacaattta atattcatac tttaagtatt ttttgtagta tcctagatat tgtgctttaa
     6241 atgctcaccc ctcaaagcac cagtaatatt ttcatccact gaaataccat taaattttca
     6301 aaaaaatact atgcatataa tgttatacat ataaacataa aacgccatgt aaatcaaaaa
     6361 atatataaaa atatgtataa aaataaatat gcactaaata taagctaatt atgcataaaa
     6421 attaaagtgc cctttattaa ctagctagtc gtaattattt atatttctat gttataaaaa
     6481 aatcctcata taataatata attaatatat gtaatgtttt ttttatttta taattttaat
     6541 ataaaataat atgtaaatta attcaaaaaa taaatataat tgttgtgaaa caaaaaacgt
     6601 aattttttca tttgccttca aaatttaaat ttattttaat atttcctaaa atatatatac
     6661 tttgtgtata aatatataaa aatatatatt tgcttataaa taaataaaaa attttataaa
     6721 acataggggg atccatggtt ggttcgctaa actgcatcgt cgctgtgtcc cagaacatgg
     6781 gcatcggcaa gaacggggac ctgccctggc caccgctcag gaacgaattt agatatttcc
     6841 agagaatgac cacaacctct tcagtagaag gtaaacagaa tctggtgatt atgggtaaga
     6901 agacctggtt ctccattcct gagaagaatc gacctttaaa gggtagaatt aatttagttc
     6961 tcagcagaga actcaaggaa cctccacaag gagctcattt tctttccaga agtctagatg
     7021 atgccttaaa acttactgaa caaccagaat tagcaaataa agtagacatg gtctggatag
     7081 ttggtggcag ttctgtttat aaggaagcca tgaatcaccc aggccatctt aaactatttg
     7141 tgacaaggat catgcaagac tttgaaagtg acacgttttt tccagaaatt gatttggaga
     7201 aatataaact tctgccagaa tacccaggtg ttctctctga tgtccaggag gagaaaggca
     7261 ttaagtacaa atttgaagta tatgagaaga atgattaagg atcccgtttt tcttacttat
     7321 atatttatac caattgattg tatttataac tgtaaaaatg tgtatgttgt gtgcatattt
     7381 ttttttgtgc atgcacatgc atgtaaatag ctaaaattat gaacatttta ttttttgttc
     7441 agaaaaaaaa aactttacac acataaaatg gctagtatga atagccatat tttatataaa
     7501 ttaaatccta tgaatttatg accatattaa aaatttagat atttatggaa cataatatgt
     7561 ttgaaacaat aagacaaaat tattattatt attattattt ttactgttat aattatgttg
     7621 tctcttcaat gattcataaa tagttggact tgatttttaa aatgtttata atatgattag
     7681 catagttaaa taaaaaaagt tgaaaaatta aaaaaaaaca tataaacaca aatgatgttt
     7741 tttccttcaa tttcgatg
//
```

```
>Py1115_PyWT-mScarlet
LOCUS       pSL1115__final_         7837 bp    DNA        circular     21-APR-2021
SOURCE
  ORGANISM
COMMENT     Lindner Lab - used to create Py1115 = Py17XNL WT-mScarlet
COMMENT     ApEinfo:methylated:1
FEATURES             Location/Qualifiers
     exon            6814..7377
                     /vntifkey="61"
                     /locus_tag="hDHFR"
                     /label="hDHFR"
                     /ApEinfo_label="hDHFR"
                     /ApEinfo_fwdcolor="pink"
                     /ApEinfo_revcolor="pink"
                     /ApEinfo_graphicformat="arrow_data {{0 0.5 0 1 2 0 0 -1 0
                     -0.5} {} 0} width 5 offset 0"
     misc_feature    1898..2471
                     /locus_tag="PbEF1a promoter"
                     /label="PbEF1a promoter"
                     /ApEinfo_label="PbEF1a promoter"
                     /ApEinfo_fwdcolor="cyan"
                     /ApEinfo_revcolor="green"
                     /ApEinfo_graphicformat="arrow_data {{0 0.5 0 1 2 0 0 -1 0
                     -0.5} {} 0} width 5 offset 0"
     misc_feature    3280..3727
                     /locus_tag="PbDHFR/TS\3'UTR"
                     /label="PbDHFR/TS\3'UTR"
                     /ApEinfo_label="PbDHFR/TS\3'UTR"
                     /ApEinfo_fwdcolor="#ff0000"
                     /ApEinfo_revcolor="green"
                     /ApEinfo_graphicformat="arrow_data {{0 0.5 0 1 2 0 0 -1 0
                     -0.5} {} 0} width 5 offset 0"
     rep_origin      5124..5806
                     /locus_tag="ColE1 origin"
                     /label="ColE1 origin"
                     /ApEinfo_label="ColE1 origin"
                     /ApEinfo_fwdcolor="gray50"
                     /ApEinfo_revcolor="gray50"
                     /ApEinfo_graphicformat="arrow_data {{0 0.5 0 1 2 0 0 -1 0
                     -0.5} {} 0} width 5 offset 0"
     CDS             4367..5026
                     /locus_tag="AmpR"
                     /label="AmpR"
                     /ApEinfo_label="AmpR"
                     /ApEinfo_fwdcolor="yellow"
                     /ApEinfo_revcolor="yellow"
                     /ApEinfo_graphicformat="arrow_data {{0 0.5 0 1 2 0 0 -1 0
                     -0.5} {} 0} width 5 offset 0"
     misc_feature    7384..7834
                     /locus_tag="PbDHFR-TS 3'UTR"
                     /label="PbDHFR-TS 3'UTR"
                     /ApEinfo_label="PbDHFR-TS 3'UTR"
                     /ApEinfo_fwdcolor="cyan"
                     /ApEinfo_revcolor="green"
                     /ApEinfo_graphicformat="arrow_data {{0 0.5 0 1 2 0 0 -1 0
                     -0.5} {} 0} width 5 offset 0"
     misc_feature    join(6210..6519,6524..6808)
                     /locus_tag="PbEF1a 5'UTR"
                     /label="PbEF1a 5'UTR"
                     /ApEinfo_label="PbEF1a 5'UTR"
                     /ApEinfo_fwdcolor="#804040"
                     /ApEinfo_revcolor="#804040"
                     /ApEinfo_graphicformat="arrow_data {{0 0.5 0 1 2 0 0 -1 0
                     -0.5} {} 0} width 5 offset 0"
     misc_feature    7..918
                     /locus_tag="py p230p Homology Arm 1"
                     /label="py p230p Homology Arm 1"
                     /ApEinfo_label="py p230p Homology Arm 1"
                     /ApEinfo_fwdcolor="cyan"
                     /ApEinfo_revcolor="green"
                     /ApEinfo_graphicformat="arrow_data {{0 0.5 0 1 2 0 0 -1 0
                     -0.5} {} 0} width 5 offset 0"
     misc_feature    925..1873
                     /locus_tag="py p230p Homology Arm 2"
                     /label="py p230p Homology Arm 2"
                     /ApEinfo_label="py p230p Homology Arm 2"
                     /ApEinfo_fwdcolor="cyan"
                     /ApEinfo_revcolor="green"
                     /ApEinfo_graphicformat="arrow_data {{0 0.5 0 1 2 0 0 -1 0
                     -0.5} {} 0} width 5 offset 0"
     misc_feature    2481..3176
                     /locus_tag="mScarlet-N1"
                     /label="mScarlet-N1"
                     /ApEinfo_label="mScarlet-N1"
                     /ApEinfo_fwdcolor="cyan"
                     /ApEinfo_revcolor="green"
                     /ApEinfo_graphicformat="arrow_data {{0 1 2 0 0 -1} {} 0}
                     width 5 offset 0"
ORIGIN
        1 ggtaccAGCT TTGTGTTTTA TTTGGATGTG CAACAATAGA TTTATGATTA ACCAATGGTA
       61 AATCATTATA ATCATATATT TCATCTTCAT TTACATATTT TACTAAAACA GTTTTTACAA
      121 TTTGTTCTGT TTCTTTATCT ACACAATGAC ATTCAAATTG ATATGATTTT GTTAATGTAT
      181 AATTAAAACC AGCTAATGCA TATTCTGGAT TTTCAAATGA ATAGTTAATA ATATTTTTAG
      241 TATTAATTTT ATAGGAATCA TGTAGATAGC CTTTTTCATA TACTTGAGTA AAACATTTTT
      301 CTGGCATTTT TTTTGTATTT TCGGGGCATC TAAAACCAAA ACTTGTATTT GATTTGGAAA
      361 TTATTATACA TAGATCTCCT TCACTATAAG CATTTAAATA ATTATATAAT TCACCTTTTC
      421 TAAAATCACA AAATGAAACT GGTCTATCTT TTAAATCTGT TCGAACATGT AATAAAATAA
      481 TTGCTCTTCT ATAATCAGTC CCACCATTTG TTACACCACA TTCTATGGTT AGTGGATATG
      541 ATCCATGATT TATAAATGAG GTCATAAAAA ATTTAAAACC TTTAAATTGT TCATTTATTT
      601 CTGGTATATC TATATCTTTT GTATTAAAAA ATAAATCTGT AAATTTATTT ATAGGAAATA
      661 CATTACTTAA ATATTTTATG GTATTAATAG GTTCTTTACC ATATACGATT CTATTATCTG
      721 TTATTTGCTC ACAATTATAC ATGAATAAAC TAAATTCATT TATTGTTTTT TCACATATTT
      781 GTATTTCACT TAATTTGCTT AAATTATAAT CAAAATTACA CATCTGATAA GGTATATCGT
      841 TAAAATATTT AATCGTTTTA CTATCATCAT CAACTATTTT TAAAATGTGA TCAGATGTTG
      901 CCATTCCATC ACCAAATGgg gcccTATGGA ACTACATCTA TATAAGAGAT TTTTTTATTT
      961 TTATATATTG GTTTTGAATC TAGATATTCA TAATTTTTAA TATATATTGT TCCTATATTT
     1021 TTGGTTTTAT AATCTTCACA TGTACAATAA AAATCAAATT CTTTTTTTGT AATTTTTGAA
     1081 ATCATTAAAA ATGTTGATGA ATTATTTATA AAATAATCTG CATCTATAAC TTTTATTTCA
     1141 TTAAATATAT TTTCGATATT TTCAGTTATT TGTTCATTCA TGTTACTTTT TATATATACC
     1201 TTTCTAAAAC AATTTTTTGG ATTAATTTTT CCAGTTGGAC AATTTAAGCT AACTACTATA
     1261 TTTTCTTTTA ATTCGATTTC ACAAAATTCA TTTTGTTTTT CTAAATCATA AACATTGGAA
     1321 TAGTAAAATT TTTTTTTAGA ATTACCTGAA AAATCACATC CATATAATTT ATTTTTTGTT
     1381 GTATTTTGAG AAACCCTTAT CACTGTAATA CCTTTTTGAC CTTTCATATT ATTAATTATT
     1441 GTTTTAGTAT TATCACAATA TATTTTCAAT ATTAATACTT TTTCTACAAT TAATGGAGTA
     1501 ACAAAAGTGA AATTTCCTTT CTCTTTGAAC ATTTTATTTT TATCTAATAT GATAACACCT
     1561 GGTAATATAT CATCTAATTT TTCTTTTTTG TCTAGATACA TTACTTCATC ATTTTCTGGT
     1621 ATGTATTCTA TATTTTTATA TTTTTCTAAT TCTTCTGGTT CAACATTTGC ATTTTCTAAA
     1681 TAAAAATTAT TTTTTTGACT ACTACGTTGT GATGTTTTCT TATTATTTAT TATGATATTA
     1741 CTATCAAAAA TGTCATCATC ACTTTTTACA ATTTTGGGAC ATTTAACTTG AACCATATTC
     1801 AGATTCTGTT CTAGATGTAT ATAACATGAA TGAACTGTAT CTACTTCATT AACGGGTTCA
     1861 AGAGAATTCG GTAgccgcgg tggcggccgc TctcgagAGC TTAATTCTTT TCGAGCTCTT
     1921 TATGCTTAAG TTTACAATTT AATATTCATA CTTTAAGTAT TTTTTGTAGT ATCCTAGATA
     1981 TTGTGCTTTA AATGCTCACC CCTCAAAGCA CCAGTAATAT TTTCATCCAC TGAAATACCA
     2041 TTAAATTTTC AAAAAAATAC TATGCATATA ATGTTATACA TATAAACATA AAACGCCATG
     2101 TAAATCAAAA AATATATAAA AATATGTATA AAAATAAATA TGCACTAAAT ATAAGCTAAT
     2161 TATGCATAAA AATTAAAGTG CCCTTTATTA ACTAGctagT CGTAATTATT TATATTTCTA
     2221 TGTTATAAAA AAATCCTCAT ATAATAATAT AATTAATATA TGTAATGTTT TTTTTATTTT
     2281 ATAATTTTAA TATAAAATAA TATGTAAATT AATTCAAAAA ATAAATATAA TTGTTGTGAA
     2341 ACAAAAAACG TAATTTTTTC ATTTGCCTTC AAAATTTAAA TTTATTTTAA TATTTCCTAA
     2401 AATATATATA CTTTGTGTAT AAATATATAA AAATATATAT TTGCTTATAA ATAAATAAAA
     2461 ATTTTATAAA Aatgactagt GTGAGCAAGG GCGAGGCAGT GATCAAGGAG TTCATGCGGT
     2521 TCAAGGTGCA CATGGAGGGC TCCATGAACG GCCACGAGTT CGAGATCGAG GGCGAGGGCG
     2581 AGGGCCGCCC CTACGAGGGC ACCCAGACCG CCAAGCTGAA GGTGACCAAG GGTGGCCCCC
     2641 TGCCCTTCTC CTGGGACATC CTGTCCCCTC AGTTCATGTA CGGCTCCAGG GCCTTCACCA
     2701 AGCACCCCGC CGACATCCCC GACTACTATA AGCAGTCCTT CCCCGAGGGC TTCAAGTGGG
     2761 AGCGCGTGAT GAACTTCGAG GACGGCGGCG CCGTGACCGT GACCCAGGAC ACCTCCCTGG
     2821 AGGACGGCAC CCTGATCTAC AAGGTGAAGC TtCGCGGCAC CAACTTCCCT CCTGACGGCC
     2881 CCGTAATGCA GAAGAAGACA ATGGGCTGGG AAGCATCCAC CGAGCGGTTG TACCCCGAGG
     2941 ACGGCGTGCT GAAGGGCGAC ATTAAGATGG CCCTGCGCCT GAAGGACGGC GGtCGCTACC
     3001 TGGCGGACTT CAAGACCACC TACAAGGCCA AGAAGCCCGT GCAGATGCCC GGCGCCTACA
     3061 ACGTCGAtCG CAAGTTGGAC ATCACCTCCC ACAACGAGGA CTACACCGTG GTGGAACAGT
     3121 ACGAACGCTC CGAGGGCCGC CACTCCACCG GCGGCATGGA CGAGCTGTAC AAGTAATTCG
     3181 AAAGATCCCA ACGAAAAGAG AGACCACATG GTCCTTCTTG AGTTTGTAAC AGCTGCTGGG
     3241 ATTACACATG GCATGGATGA ACTATACAAA TAAggatccG TTTTTCTTAC TTATATATTT
     3301 ATACCAATTG ATTGTATTTA TAACTGTAAA AATGTGTATG TTGTGTGCAT ATTTTTTTTT
     3361 GTGCATGCAC ATGCATGTAA ATAGCTAAAA TTATGAACAT TTTATTTTTT GTTCAGAAAA
     3421 AAAAAACTTT ACACACATAA AATGGCTAGT ATGAATAGCC ATATTTTATA TAAATTAAAT
     3481 CCTATGAATT TATGACCATA TTAAAAATTT AGATATTTAT GGAACATAAT ATGTTTGAAA
     3541 CAATAAGACA AAATTATTAT TATTATTATT ATTTTTACTG TTATAATTAT GTTGTCTCTT
     3601 CAATGATTCA TAAATAGTTG GACTTGATTT TTAAAATGTT TATAATATGA TTAGCATAGT
     3661 TAAATAAAAA AAGTTGAAAA ATTAAAAAAA AACATATAAA CACAAATGAT GTTTTTTCCT
     3721 TCAATTTcgg cgcctgatgc ggtattttct ccttacgcat ctgtgcggta tttcacaccg
     3781 catatggtgc actctcagta caatctgctc tgatgccgca tagttaagcc agccccgaca
     3841 cccgccaaca cccgctgacg cgccctgacg ggcttgtctg ctcccggcat ccgcttacag
     3901 acaagctgtg accgtctccg ggagctgcat gtgtcagagg ttttcaccgt catcaccgaa
     3961 acgcgcgaga cgaaagggcc tcgtgatacg cctattttta taggttaatg tcatgataat
     4021 aatggtttct tagacgtcag gtggcacttt tcggggaaat gtgcgcggaa cccctatttg
     4081 tttatttttc taaatacatt caaatatgta tccgctcatg agacaataac cctgataaat
     4141 gcttcaataa tattgaaaaa ggaagagtat gagtattcaa catttccgtg tcgcccttat
     4201 tccctttttt gcggcatttt gccttcctgt ttttgctcac ccagaaacgc tggtgaaagt
     4261 aaaagatgct gaagatcagt tgggtgcacg agtgggttac atcgaactgg atctcaacag
     4321 cggtaagatc cttgagagtt ttcgccccga agaacgtttt ccaatgatga gcacttttaa
     4381 agttctgcta tgtggcgcgg tattatcccg tattgacgcc gggcaagagc aactcggtcg
     4441 ccgcatacac tattctcaga atgacttggt tgagtactca ccagtcacag aaaagcatct
     4501 tacggatggc atgacagtaa gagaattatg cagtgctgcc ataaccatga gtgataacac
     4561 tgcggccaac ttacttctga caacgatcgg aggaccgaag gagctaaccg cttttttgca
     4621 caacatgggg gatcatgtaa ctcgccttga tcgttgggaa ccggagctga atgaagccat
     4681 accaaacgac gagcgtgaca ccacgatgcc tgtagcaatg gcaacaacgt tgcgcaaact
     4741 attaactggc gaactactta ctctagcttc ccggcaacaa ttaatagact ggatggaggc
     4801 ggataaagtt gcaggaccac ttctgcgctc ggcccttccg gctggctggt ttattgctga
     4861 taaatctgga gccggtgagc gtgggtctcg cggtatcatt gcagcactgg ggccagatgg
     4921 taagccctcc cgtatcgtag ttatctacac gacggggagt caggcaacta tggatgaacg
     4981 aaatagacag atcgctgaga taggtgcctc actgattaag cattggtaac tgtcagacca
     5041 agtttactca tatatacttt agattgattt aaaacttcat ttttaattta aaaggatcta
     5101 ggtgaagatc ctttttgata atctcatgac caaaatccct taacgtgagt tttcgttcca
     5161 ctgagcgtca gaccccgtag aaaagatcaa aggatcttct tgagatcctt tttttctgcg
     5221 cgtaatctgc tgcttgcaaa caaaaaaacc accgctacca gcggtggttt gtttgccgga
     5281 tcaagagcta ccaactcttt ttccgaaggt aactggcttc agcagagcgc agataccaaa
     5341 tactgtcctt ctagtgtagc cgtagttagg ccaccacttc aagaactctg tagcaccgcc
     5401 tacatacctc gctctgctaa tcctgttacc agtggctgct gccagtggcg ataagtcgtg
     5461 tcttaccggg ttggactcaa gacgatagtt accggataag gcgcagcggt cgggctgaac
     5521 ggggggttcg tgcacacagc ccagcttgga gcgaacgacc tacaccgaac tgagatacct
     5581 acagcgtgag cattgagaaa gcgccacgct tcccgaaggg agaaaggcgg acaggtatcc
     5641 ggtaagcggc agggtcggaa caggagagcg cacgagggag cttccagggg gaaacgcctg
     5701 gtatctttat agtcctgtcg ggtttcgcca cctctgactt gagcgtcgat ttttgtgatg
     5761 ctcgtcaggg gggcggagcc tatggaaaaa cgccagcaac gcggcctttt tacggttcct
     5821 ggccttttgc tggccttttg ctcacatgtt ctttcctgcg ttatcccctg attctgtgga
     5881 taaccgtatt accgcctttg agtgagctga taccgctcgc cgcagccgaa cgaccgagcg
     5941 cagcgagtca gtgagcgagg aagcggaaga gcgcccaata cgcaaaccgc ctctccccgc
     6001 gcgttggccg attcattaat gcagctggca cgacaggttt cccgactgga aagcgggcag
     6061 tgagcgcaac gcaattaatg tgagttagct cactcattag gcaccccagg ctttacactt
     6121 tatgcttccg gctcgtatgt tgtgtggaat tgtgagcgga taacaatttc acacaggaaa
     6181 cagctatgac catgattacg ccaagcttga taattcctgc agcccagctt aattcttttc
     6241 gagctcttta tgcttaagtt tacaatttaa tattcatact ttaagtattt tttgtagtat
     6301 cctagatatt gtgctttaaa tgctcacccc tcaaagcacc agtaatattt tcatccactg
     6361 aaataccatt aaattttcaa aaaaatacta tgcatataat gttatacata taaacataaa
     6421 acgccatgta aatcaaaaaa tatataaaaa tatgtataaa aataaatatg cactaaatat
     6481 aagctaatta tgcataaaaa ttaaagtgcc ctttattaac tagctagtcg taattattta
     6541 tatttctatg ttataaaaaa atcctcatat aataatataa ttaatatatg taatgttttt
     6601 tttattttat aattttaata taaaataata tgtaaattaa ttcaaaaaat aaatataatt
     6661 gttgtgaaac aaaaaacgta attttttcat ttgccttcaa aatttaaatt tattttaata
     6721 tttcctaaaa tatatatact ttgtgtataa atatataaaa atatatattt gcttataaat
     6781 aaataaaaaa ttttataaaa cataggggga tccatggttg gttcgctaaa ctgcatcgtc
     6841 gctgtgtccc agaacatggg catcggcaag aacggggacc tgccctggcc accgctcagg
     6901 aacgaattta gatatttcca gagaatgacc acaacctctt cagtagaagg taaacagaat
     6961 ctggtgatta tgggtaagaa gacctggttc tccattcctg agaagaatcg acctttaaag
     7021 ggtagaatta atttagttct cagcagagaa ctcaaggaac ctccacaagg agctcatttt
     7081 ctttccagaa gtctagatga tgccttaaaa cttactgaac aaccagaatt agcaaataaa
     7141 gtagacatgg tctggatagt tggtggcagt tctgtttata aggaagccat gaatcaccca
     7201 ggccatctta aactatttgt gacaaggatc atgcaagact ttgaaagtga cacgtttttt
     7261 ccagaaattg atttggagaa atataaactt ctgccagaat acccaggtgt tctctctgat
     7321 gtccaggagg agaaaggcat taagtacaaa tttgaagtat atgagaagaa tgattaagga
     7381 tcccgttttt cttacttata tatttatacc aattgattgt atttataact gtaaaaatgt
     7441 gtatgttgtg tgcatatttt tttttgtgca tgcacatgca tgtaaatagc taaaattatg
     7501 aacattttat tttttgttca gaaaaaaaaa actttacaca cataaaatgg ctagtatgaa
     7561 tagccatatt ttatataaat taaatcctat gaatttatga ccatattaaa aatttagata
     7621 tttatggaac ataatatgtt tgaaacaata agacaaaatt attattatta ttattatttt
     7681 tactgttata attatgttgt ctcttcaatg attcataaat agttggactt gatttttaaa
     7741 atgtttataa tatgattagc atagttaaat aaaaaaagtt gaaaaattaa aaaaaaacat
     7801 ataaacacaa atgatgtttt ttccttcaat ttcgatg
//
```

>Py1065_*pynot1-g^-^*

```
LOCUS       pSL1065                 6892 bp    DNA        circular     21-APR-2021
SOURCE
  ORGANISM
COMMENT     Lindner Lab - used to create Py1065 = Py17XNL deletion of pynot1-g
COMMENT     ApEinfo:methylated:1
FEATURES             Location/Qualifiers
     exon            5869..6432
                     /vntifkey="61"
                     /locus_tag="hDHFR"
                     /label="hDHFR"
                     /ApEinfo_label="hDHFR"
                     /ApEinfo_fwdcolor="pink"
                     /ApEinfo_revcolor="pink"
                     /ApEinfo_graphicformat="arrow_data {{0 1 2 0 0 -1} {} 0}
                     width 5 offset 0"
     misc_feature    1650..2328
                     /locus_tag="GFPmut2"
                     /label="GFPmut2"
                     /ApEinfo_label="GFPmut2"
                     /ApEinfo_fwdcolor="cyan"
                     /ApEinfo_revcolor="green"
                     /ApEinfo_graphicformat="arrow_data {{0 1 2 0 0 -1} {} 0}
                     width 5 offset 0"
     misc_feature    2335..2782
                     /locus_tag="PbDHFR/TS\3'UTR"
                     /label="PbDHFR/TS\3'UTR"
                     /ApEinfo_label="PbDHFR/TS\3'UTR"
                     /ApEinfo_fwdcolor="#ff0000"
                     /ApEinfo_revcolor="green"
                     /ApEinfo_graphicformat="arrow_data {{0 1 2 0 0 -1} {} 0}
                     width 5 offset 0"
     rep_origin      4179..4861
                     /locus_tag="ColE1 origin"
                     /label="ColE1 origin"
                     /ApEinfo_label="ColE1 origin"
                     /ApEinfo_fwdcolor="gray50"
                     /ApEinfo_revcolor="gray50"
                     /ApEinfo_graphicformat="arrow_data {{0 1 2 0 0 -1} {} 0}
                     width 5 offset 0"
     CDS             3422..4081
                     /locus_tag="AmpR"
                     /label="AmpR"
                     /ApEinfo_label="AmpR"
                     /ApEinfo_fwdcolor="yellow"
                     /ApEinfo_revcolor="yellow"
                     /ApEinfo_graphicformat="arrow_data {{0 1 2 0 0 -1} {} 0}
                     width 5 offset 0"
     misc_feature    join(5265..5574,5579..5863)
                     /locus_tag="PbEF1a 5'UTR"
                     /label="PbEF1a 5'UTR"
                     /ApEinfo_label="PbEF1a 5'UTR"
                     /ApEinfo_fwdcolor="#804040"
                     /ApEinfo_revcolor="#804040"
                     /ApEinfo_graphicformat="arrow_data {{0 1 2 0 0 -1} {} 0}
                     width 5 offset 0"
     misc_feature    6440..6887
                     /locus_tag="PbDHFR-TS 3'UTR"
                     /label="PbDHFR-TS 3'UTR"
                     /ApEinfo_label="PbDHFR-TS 3'UTR"
                     /ApEinfo_fwdcolor="cyan"
                     /ApEinfo_revcolor="green"
                     /ApEinfo_graphicformat="arrow_data {{0 1 2 0 0 -1} {} 0}
                     width 5 offset 0"
     misc_feature    748..1608
                     /locus_tag="PyNOT1-G 5' Homology Arm"
                     /label="PyNOT1-G 5' Homology Arm"
                     /ApEinfo_label="PyNOT1-G 5' Homology Arm"
                     /ApEinfo_fwdcolor="cyan"
                     /ApEinfo_revcolor="green"
                     /ApEinfo_graphicformat="arrow_data {{0 0.5 0 1 2 0 0 -1 0
                     -0.5} {} 0} width 5 offset 0"
     misc_feature    7..730
                     /locus_tag="PyNOT1-G 3' Homology Arm"
                     /label="PyNOT1-G 3' Homology Arm"
                     /ApEinfo_label="PyNOT1-G 3' Homology Arm"
                     /ApEinfo_fwdcolor="cyan"
                     /ApEinfo_revcolor="green"
                     /ApEinfo_graphicformat="arrow_data {{0 0.5 0 1 2 0 0 -1 0
                     -0.5} {} 0} width 5 offset 0"
ORIGIN
        1 ggtaccCAAC TCATACAAAC TCTTCTTAGA ATGTAATATA TATGTACATG TATATATATT
       61 ATTATTATTA TTATTTTATT TTTTTTTTTT TTTTTTTTTA CTTACACATG TATGTATGCA
      121 CATACATATA TTACAATTTT ATACCATTTT AAAATTAGTC TTATTCATCA TCGAAGAAAA
      181 ATTTATTATC CTTTGCATAC ATACATACAT ACATATATAT ATATATATAT ACATATATAT
      241 ATACATGCAT TTATTTTACT ATTTTTTTAC AATTTATAAT TGTATTTTTA AAATAGCAGT
      301 TTTGTAATTT TTTTCAAATT CATATTTTTT TTTTCCCATT TTTTTGTTAA ACTTTTAAAA
      361 CACAACAATA GTATCGATAG TACTAGTAGT ATATAATTAT AAAATGCGTT TATTTATTTT
      421 TTCTACATTT CTACAATTTT TCTACAATTT TCCTACAATT TTAAAATACC ACATATAAAC
      481 CTCAATACAC ACATCCATTC CGTTTTAATA AAAAAAAAAA TATATATTTC CGAAATAACG
      541 CTAGGAGGAA AATAAAATTC CCCTGATTTT TACTATCATT TTAATGTTTT CATTATTCTA
      601 TATCTTTAAT AATATACATG AAATGTAATT TTTATGTGTG CAAAAATATT TCAAGTTTAT
      661 TATTAAGCAA AAAATTTACA AAAATATATA CACATATGAG AAATATGAAT CCCGCTTTGT
      721 CTATCCATAC ggcggccgcg cgggcccCCT AGTTTATTTA TCTACAAACG TATGCCCACA
      781 ATTTCAACTA TACAATGTGT TATATGTATA ATAAATATTT GTATACAATA TATACTTATT
      841 AATATAATAC AATATATATA TATATATACA TAATTAATAT ATTTTTAGTG CATATAAATG
      901 TACTTATTTA ATTCGTAAAA ATATTGCTAA TTAAACCCCA ATATTGTATA CATATACTAA
      961 GGACATAAAT TCCTCTACGG TTTTATTCAC ATGCATTAAA AAAATATAAC TTAAGTATAT
     1021 TTTTTCTGTA TATCCCCAAA AAAAAAAATG TGGTCAAAAA AAATATTATA TATATATAAT
     1081 TAATTTTATA GGAAAAAAAC ATTCCCAATA AAATTAAAAA AAATTCTCAA TTTTTTTAAT
     1141 ATTATTTTCC TTTTTGGTGA TATTCATATA TAGTTAATGA ATATAACATA ACACCAATTT
     1201 ATAATCCTGC GTTTACTTTA TATGTAAAGT TTTCATAAAA TCATAATAAC TACGCAGTTA
     1261 TAATAAGTTA AAAAGGGAAA ATGAAAAGAA AAAAAATATA TTTTTCTATA TGTGTGTGCA
     1321 TATATATTGT ATATATATAT TTATAATCAT AAACAAAATA AGGCAAAAAA AAATACCAAA
     1381 AAAAAATAAT AAAAAATAAA ATAAATAAAT AACCTCCAAA AAAATAGAAA ACAATCCAAA
     1441 CAAGCCAAAC ATACCAAACA TACCAAACTT AGCAATCACA GCTGAAATTA AAAAAATAAC
     1501 AAACACACCA AATCGTGTAA ACGCAAAAAA TACCACATAA CAAAACACAG TCAAATACCC
     1561 ATATATATTG TTCATACCAT ATTGAGGTAA TTACAATAAA ACATGTGCcc tagtagtaaa
     1621 ggagaagaac ttttcactgg agttgtcccA ATTCTTGTTG AATTAGATGG TGATGTTAAT
     1681 GGGCACAAAT TTTCTGTCAG TGGAGAGGGT GAAGGTGATG CAACATACGG AAAACTTACC
     1741 CTTAAATTTA TTTGCACTAC TGGAAAACTA CCTGTTCCAT GGCCAACACT TGTCACTACT
     1801 TTCGCGTATG GTCTTCAATG CTTTGCGAGA TACCCAGATC ATATGAAACA GCATGACTTT
     1861 TTCAAGAGTG CCATGCCCGA AGGTTATGTA CAGGAAAGAA CTATATTTTT CAAAGATGAC
     1921 GGGAACTACA AGACACGTGC TGAAGTCAAG TTTGAAGGTG ATACCCTTGT TAATAGAATC
     1981 GAGTTAAAAG GTATTGATTT TAAAGAAGAT GGAAACATTC TTGGACACAA ATTGGAATAC
     2041 AACTATAACT CACACAATGT ATACATCATG GCAGACAAAC AAAAGAATGG AATCAAAGTT
     2101 AACTTCAAAA TTAGACACAA CATTGAAGAT GGAAGCGTTC AACTAGCAGA CCATTATCAA
     2161 CAAAATACTC CAATTGGCGA TGGCCCTGTC CTTTTACCAG ACAACCATTA CCTGTCCACA
     2221 CAATCTGCCC TTTCGAAAGA TCCCAACGAA AAGAGAGACC ACATGGTCCT TCTTGAGTTT
     2281 GTAACAGCTG CTGGGATTAC ACATGGCATG GATGAACTAT ACAAATAAgg atccGTTTTT
     2341 CTTACTTATA TATTTATACC AATTGATTGT ATTTATAACT GTAAAAATGT GTATGTTGTG
     2401 TGCATATTTT TTTTTGTGCA TGCACATGCA TGTAAATAGC TAAAATTATG AACATTTTAT
     2461 TTTTTGTTCA GAAAAAAAAA ACTTTACACA CATAAAATGG CTAGTATGAA TAGCCATATT
     2521 TTATATAAAT TAAATCCTAT GAATTTATGA CCATATTAAA AATTTAGATA TTTATGGAAC
     2581 ATAATATGTT TGAAACAATA AGACAAAATT ATTATTATTA TTATTATTTT TACTGTTATA
     2641 ATTATGTTGT CTCTTCAATG ATTCATAAAT AGTTGGACTT GATTTTTAAA ATGTTTATAA
     2701 TATGATTAGC ATAGTTAAAT AAAAAAAGTT GAAAAATTAA AAAAAAACAT ATAAACACAA
     2761 ATGATGTTTT TTCCTTCAAT TTcggcgcct gatgcggtat tttctcctta cgcatctgtg
     2821 cggtatttca caccgcatat ggtgcactct cagtacaatc tgctctgatg ccgcatagtt
     2881 aagccagccc cgacacccgc caacacccgc tgacgcgccc tgacgggctt gtctgctccc
     2941 ggcatccgct tacagacaag ctgtgaccgt ctccgggagc tgcatgtgtc agaggttttc
     3001 accgtcatca ccgaaacgcg cgagacgaaa gggcctcgtg atacgcctat ttttataggt
     3061 taatgtcatg ataataatgg tttcttagac gtcaggtggc acttttcggg gaaatgtgcg
     3121 cggaacccct atttgtttat ttttctaaat acattcaaat atgtatccgc tcatgagaca
     3181 ataaccctga taaatgcttc aataatattg aaaaaggaag agtatgagta ttcaacattt
     3241 ccgtgtcgcc cttattccct tttttgcggc attttgcctt cctgtttttg ctcacccaga
     3301 aacgctggtg aaagtaaaag atgctgaaga tcagttgggt gcacgagtgg gttacatcga
     3361 actggatctc aacagcggta agatccttga gagttttcgc cccgaagaac gttttccaat
     3421 gatgagcact tttaaagttc tgctatgtgg cgcggtatta tcccgtattg acgccgggca
     3481 agagcaactc ggtcgccgca tacactattc tcagaatgac ttggttgagt actcaccagt
     3541 cacagaaaag catcttacgg atggcatgac agtaagagaa ttatgcagtg ctgccataac
     3601 catgagtgat aacactgcgg ccaacttact tctgacaacg atcggaggac cgaaggagct
     3661 aaccgctttt ttgcacaaca tgggggatca tgtaactcgc cttgatcgtt gggaaccgga
     3721 gctgaatgaa gccataccaa acgacgagcg tgacaccacg atgcctgtag caatggcaac
     3781 aacgttgcgc aaactattaa ctggcgaact acttactcta gcttcccggc aacaattaat
     3841 agactggatg gaggcggata aagttgcagg accacttctg cgctcggccc ttccggctgg
     3901 ctggtttatt gctgataaat ctggagccgg tgagcgtggg tctcgcggta tcattgcagc
     3961 actggggcca gatggtaagc cctcccgtat cgtagttatc tacacgacgg ggagtcaggc
     4021 aactatggat gaacgaaata gacagatcgc tgagataggt gcctcactga ttaagcattg
     4081 gtaactgtca gaccaagttt actcatatat actttagatt gatttaaaac ttcattttta
     4141 atttaaaagg atctaggtga agatcctttt tgataatctc atgaccaaaa tcccttaacg
     4201 tgagttttcg ttccactgag cgtcagaccc cgtagaaaag atcaaaggat cttcttgaga
     4261 tccttttttt ctgcgcgtaa tctgctgctt gcaaacaaaa aaaccaccgc taccagcggt
     4321 ggtttgtttg ccggatcaag agctaccaac tctttttccg aaggtaactg gcttcagcag
     4381 agcgcagata ccaaatactg tccttctagt gtagccgtag ttaggccacc acttcaagaa
     4441 ctctgtagca ccgcctacat acctcgctct gctaatcctg ttaccagtgg ctgctgccag
     4501 tggcgataag tcgtgtctta ccgggttgga ctcaagacga tagttaccgg ataaggcgca
     4561 gcggtcgggc tgaacggggg gttcgtgcac acagcccagc ttggagcgaa cgacctacac
     4621 cgaactgaga tacctacagc gtgagcattg agaaagcgcc acgcttcccg aagggagaaa
     4681 ggcggacagg tatccggtaa gcggcagggt cggaacagga gagcgcacga gggagcttcc
     4741 agggggaaac gcctggtatc tttatagtcc tgtcgggttt cgccacctct gacttgagcg
     4801 tcgatttttg tgatgctcgt caggggggcg gagcctatgg aaaaacgcca gcaacgcggc
     4861 ctttttacgg ttcctggcct tttgctggcc ttttgctcac atgttctttc ctgcgttatc
     4921 ccctgattct gtggataacc gtattaccgc ctttgagtga gctgataccg ctcgccgcag
     4981 ccgaacgacc gagcgcagcg agtcagtgag cgaggaagcg gaagagcgcc caatacgcaa
     5041 accgcctctc cccgcgcgtt ggccgattca ttaatgcagc tggcacgaca ggtttcccga
     5101 ctggaaagcg ggcagtgagc gcaacgcaat taatgtgagt tagctcactc attaggcacc
     5161 ccaggcttta cactttatgc ttccggctcg tatgttgtgt ggaattgtga gcggataaca
     5221 atttcacaca ggaaacagct atgaccatga ttacgccaag cttgataatt cctgcagccc
     5281 agcttaattc ttttcgagct ctttatgctt aagtttacaa tttaatattc atactttaag
     5341 tattttttgt agtatcctag atattgtgct ttaaatgctc acccctcaaa gcaccagtaa
     5401 tattttcatc cactgaaata ccattaaatt ttcaaaaaaa tactatgcat ataatgttat
     5461 acatataaac ataaaacgcc atgtaaatca aaaaatatat aaaaatatgt ataaaaataa
     5521 atatgcacta aatataagct aattatgcat aaaaattaaa gtgcccttta ttaactagct
     5581 agtcgtaatt atttatattt ctatgttata aaaaaatcct catataataa tataattaat
     5641 atatgtaatg ttttttttat tttataattt taatataaaa taatatgtaa attaattcaa
     5701 aaaataaata taattgttgt gaaacaaaaa acgtaatttt ttcatttgcc ttcaaaattt
     5761 aaatttattt taatatttcc taaaatatat atactttgtg tataaatata taaaaatata
     5821 tatttgctta taaataaata aaaaatttta taaaacatag ggggatccat ggttggttcg
     5881 ctaaactgca tcgtcgctgt gtcccagaac atgggcatcg gcaagaacgg ggacctgccc
     5941 tggccaccgc tcaggaacga atttagatat ttccagagaa tgaccacaac ctcttcagta
     6001 gaaggtaaac agaatctggt gattatgggt aagaagacct ggttctccat tcctgagaag
     6061 aatcgacctt taaagggtag aattaattta gttctcagca gagaactcaa ggaacctcca
     6121 caaggagctc attttctttc cagaagtcta gatgatgcct taaaacttac tgaacaacca
     6181 gaattagcaa ataaagtaga catggtctgg atagttggtg gcagttctgt ttataaggaa
     6241 gccatgaatc acccaggcca tcttaaacta tttgtgacaa ggatcatgca agactttgaa
     6301 agtgacacgt tttttccaga aattgatttg gagaaatata aacttctgcc agaataccca
     6361 ggtgttctct ctgatgtcca ggaggagaaa ggcattaagt acaaatttga agtatatgag
     6421 aagaatgatt aaggatcccg tttttcttac ttatatattt ataccaattg attgtattta
     6481 taactgtaaa aatgtgtatg ttgtgtgcat attttttttt gtgcatgcac atgcatgtaa
     6541 atagctaaaa ttatgaacat tttatttttt gttcagaaaa aaaaaacttt acacacataa
     6601 aatggctagt atgaatagcc atattttata taaattaaat cctatgaatt tatgaccata
     6661 ttaaaaattt agatatttat ggaacataat atgtttgaaa caataagaca aaattattat
     6721 tattattatt atttttactg ttataattat gttgtctctt caatgattca taaatagttg
     6781 gacttgattt ttaaaatgtt tataatatga ttagcatagt taaataaaaa aagttgaaaa
     6841 attaaaaaaa aacatataaa cacaaatgat gttttttcct tcaatttcga tg
//
```

>Py1061_*pynot1^-^*

```
LOCUS       pSL1061__final_         6929 bp    DNA        circular     21-APR-2021
SOURCE
  ORGANISM
COMMENT     Lindner Lab - used to create Py1061 = Py17XNL deletion of pynot1
COMMENT     ApEinfo:methylated:1
FEATURES             Location/Qualifiers
     exon            5163..5726
                     /vntifkey="61"
                     /locus_tag="hDHFR"
                     /label="hDHFR"
                     /ApEinfo_label="hDHFR"
                     /ApEinfo_fwdcolor="pink"
                     /ApEinfo_revcolor="pink"
                     /ApEinfo_graphicformat="arrow_data {{0 1 2 0 0 -1} {} 0}
                     width 5 offset 0"
     misc_feature    944..1622
                     /locus_tag="GFPmut2"
                     /label="GFPmut2"
                     /ApEinfo_label="GFPmut2"
                     /ApEinfo_fwdcolor="cyan"
                     /ApEinfo_revcolor="green"
                     /ApEinfo_graphicformat="arrow_data {{0 1 2 0 0 -1} {} 0}
                     width 5 offset 0"
     misc_feature    5733..6183
                     /vntifkey="21"
                     /locus_tag="PbDHFR/TS\3'UTR"
                     /label="PbDHFR/TS\3'UTR"
                     /ApEinfo_label="PbDHFR/TS\3'UTR"
                     /ApEinfo_fwdcolor="#eb0214"
                     /ApEinfo_revcolor="#eb0214"
                     /ApEinfo_graphicformat="arrow_data {{0 1 2 0 0 -1} {} 0}
                     width 5 offset 0"
     misc_feature    1629..2076
                     /locus_tag="PbDHFR/TS\3'UTR(1)"
                     /label="PbDHFR/TS\3'UTR(1)"
                     /ApEinfo_label="PbDHFR/TS\3'UTR"
                     /ApEinfo_fwdcolor="#ff0000"
                     /ApEinfo_revcolor="green"
                     /ApEinfo_graphicformat="arrow_data {{0 1 2 0 0 -1} {} 0}
                     width 5 offset 0"
     misc_feature    45..878
                     /locus_tag="pynot1 - 5' Homology Arm (KO)"
                     /label="pynot1 - 5' Homology Arm (KO)"
                     /ApEinfo_label="pynot1 - 5' Homology Arm (KO)"
                     /ApEinfo_fwdcolor="cyan"
                     /ApEinfo_revcolor="green"
                     /ApEinfo_graphicformat="arrow_data {{0 1 2 0 0 -1} {} 0}
                     width 5 offset 0"
     rep_origin      3473..4155
                     /locus_tag="ColE1 origin"
                     /label="ColE1 origin"
                     /ApEinfo_label="ColE1 origin"
                     /ApEinfo_fwdcolor="gray50"
                     /ApEinfo_revcolor="gray50"
                     /ApEinfo_graphicformat="arrow_data {{0 1 2 0 0 -1} {} 0}
                     width 5 offset 0"
     CDS             2716..3375
                     /locus_tag="AmpR"
                     /label="AmpR"
                     /ApEinfo_label="AmpR"
                     /ApEinfo_fwdcolor="yellow"
                     /ApEinfo_revcolor="yellow"
                     /ApEinfo_graphicformat="arrow_data {{0 1 2 0 0 -1} {} 0}
                     width 5 offset 0"
     misc_feature    join(4559..4868,4873..5157)
                     /locus_tag="PbEF1a 5'UTR"
                     /label="PbEF1a 5'UTR"
                     /ApEinfo_label="PbEF1a 5'UTR"
                     /ApEinfo_fwdcolor="#804040"
                     /ApEinfo_revcolor="#804040"
                     /ApEinfo_graphicformat="arrow_data {{0 1 2 0 0 -1} {} 0}
                     width 5 offset 0"
     misc_feature    6193..6912
                     /locus_tag="pynot1 - 3' homology arm (KO)"
                     /label="pynot1 - 3' homology arm (KO)"
                     /ApEinfo_label="pynot1 - 3' homology arm (KO)"
                     /ApEinfo_fwdcolor="cyan"
                     /ApEinfo_revcolor="green"
                     /ApEinfo_graphicformat="arrow_data {{0 0.5 0 1 2 0 0 -1 0
                     -0.5} {} 0} width 5 offset 0"
ORIGIN
        1 CCAACTTGCC CCCCCAATAA CAGTTATATA TATGTACCAA AAAACAACCT AGAATTTACT
       61 TAAATTTCCA TACCAATTTA ATATATTGTA TAAATGAATA CAAAGAAAAA ATGGAAAAAA
      121 ATACGACTTA AAAGGTATAG AAACGCCGAA TTCTCAAACT GTATATATGG ATATTATGCA
      181 CATACAATTA TTATACTATG AACCAATATA TTATATATAT ATATATATAT AATATATATA
      241 TATAATGAGT GCATATGATA AATATACTTA ATAAACATAA CCAGAAAACA TATTCTCTTA
      301 CCATTTTAAA GAAATTCACA CATTAAATTT GTATATATAT TAAATATATT ATATAATATA
      361 TTATATTATA TATATTAATA TTGTAAATAT TTATTTTGCC ATATATTATA TTAGCTTATA
      421 TGTAATTACG TACAGTATAG CGCAACGAGT ATTTGAATAA ATTTTTCTTT TAATCAACCT
      481 TATTATATTT TTTCATAGAT AATTTTTTTA TTTTTTATTT CCTTTTCATT GATATATTTT
      541 ATTATTTCAT TTTTATGTAT GTATATATAT ATATAATAAA TAGTTTAAAC AAAAATTGTA
      601 TTCAATATAT ATGCATATAT AATAATTATT TATAATTTTT TTTTAACAAG TTTTTAAACA
      661 GTTAAATCCG TATTTAGTAT ATATGTTTTT TTAATATATA TAAGTATATG TATGTGTTAT
      721 GTATATATAT ATTTTAAATA TGTCACAATA TTTTTAAAAC ACACAATTAT AGGATTTATA
      781 TAAAAATAAA ATAAGTAACA AAAGAGAAGA TATAACGCAT TTATGTACAT ATAATATATT
      841 TATCGTTGTA CAATTGTAAA TATGAATTTT TACATATCAA AATATGAGAA TGGATTACCA
      901 TGcctagtag taaaggagaa gaacttttca ctggagttgt cccAATTCTT GTTGAATTAG
      961 ATGGTGATGT TAATGGGCAC AAATTTTCTG TCAGTGGAGA GGGTGAAGGT GATGCAACAT
     1021 ACGGAAAACT TACCCTTAAA TTTATTTGCA CTACTGGAAA ACTACCTGTT CCATGGCCAA
     1081 CACTTGTCAC TACTTTCGCG TATGGTCTTC AATGCTTTGC GAGATACCCA GATCATATGA
     1141 AACAGCATGA CTTTTTCAAG AGTGCCATGC CCGAAGGTTA TGTACAGGAA AGAACTATAT
     1201 TTTTCAAAGA TGACGGGAAC TACAAGACAC GTGCTGAAGT CAAGTTTGAA GGTGATACCC
     1261 TTGTTAATAG AATCGAGTTA AAAGGTATTG ATTTTAAAGA AGATGGAAAC ATTCTTGGAC
     1321 ACAAATTGGA ATACAACTAT AACTCACACA ATGTATACAT CATGGCAGAC AAACAAAAGA
     1381 ATGGAATCAA AGTTAACTTC AAAATTAGAC ACAACATTGA AGATGGAAGC GTTCAACTAG
     1441 CAGACCATTA TCAACAAAAT ACTCCAATTG GCGATGGCCC TGTCCTTTTA CCAGACAACC
     1501 ATTACCTGTC CACACAATCT GCCCTTTCGA AAGATCCCAA CGAAAAGAGA GACCACATGG
     1561 TCCTTCTTGA GTTTGTAACA GCTGCTGGGA TTACACATGG CATGGATGAA CTATACAAAT
     1621 AAggatccGT TTTTCTTACT TATATATTTA TACCAATTGA TTGTATTTAT AACTGTAAAA
     1681 ATGTGTATGT TGTGTGCATA TTTTTTTTTG TGCATGCACA TGCATGTAAA TAGCTAAAAT
     1741 TATGAACATT TTATTTTTTG TTCAGAAAAA AAAAACTTTA CACACATAAA ATGGCTAGTA
     1801 TGAATAGCCA TATTTTATAT AAATTAAATC CTATGAATTT ATGACCATAT TAAAAATTTA
     1861 GATATTTATG GAACATAATA TGTTTGAAAC AATAAGACAA AATTATTATT ATTATTATTA
     1921 TTTTTACTGT TATAATTATG TTGTCTCTTC AATGATTCAT AAATAGTTGG ACTTGATTTT
     1981 TAAAATGTTT ATAATATGAT TAGCATAGTT AAATAAAAAA AGTTGAAAAA TTAAAAAAAA
     2041 ACATATAAAC ACAAATGATG TTTTTTCCTT CAATTTcggc gcctgatgcg gtattttctc
     2101 cttacgcatc tgtgcggtat ttcacaccgc atatggtgca ctctcagtac aatctgctct
     2161 gatgccgcat agttaagcca gccccgacac ccgccaacac ccgctgacgc gccctgacgg
     2221 gcttgtctgc tcccggcatc cgcttacaga caagctgtga ccgtctccgg gagctgcatg
     2281 tgtcagaggt tttcaccgtc atcaccgaaa cgcgcgagac gaaagggcct cgtgatacgc
     2341 ctatttttat aggttaatgt catgataata atggtttctt agacgtcagg tggcactttt
     2401 cggggaaatg tgcgcggaac ccctatttgt ttatttttct aaatacattc aaatatgtat
     2461 ccgctcatga gacaataacc ctgataaatg cttcaataat attgaaaaag gaagagtatg
     2521 agtattcaac atttccgtgt cgcccttatt cccttttttg cggcattttg ccttcctgtt
     2581 tttgctcacc cagaaacgct ggtgaaagta aaagatgctg aagatcagtt gggtgcacga
     2641 gtgggttaca tcgaactgga tctcaacagc ggtaagatcc ttgagagttt tcgccccgaa
     2701 gaacgttttc caatgatgag cacttttaaa gttctgctat gtggcgcggt attatcccgt
     2761 attgacgccg ggcaagagca actcggtcgc cgcatacact attctcagaa tgacttggtt
     2821 gagtactcac cagtcacaga aaagcatctt acggatggca tgacagtaag agaattatgc
     2881 agtgctgcca taaccatgag tgataacact gcggccaact tacttctgac aacgatcgga
     2941 ggaccgaagg agctaaccgc ttttttgcac aacatggggg atcatgtaac tcgccttgat
     3001 cgttgggaac cggagctgaa tgaagccata ccaaacgacg agcgtgacac cacgatgcct
     3061 gtagcaatgg caacaacgtt gcgcaaacta ttaactggcg aactacttac tctagcttcc
     3121 cggcaacaat taatagactg gatggaggcg gataaagttg caggaccact tctgcgctcg
     3181 gcccttccgg ctggctggtt tattgctgat aaatctggag ccggtgagcg tgggtctcgc
     3241 ggtatcattg cagcactggg gccagatggt aagccctccc gtatcgtagt tatctacacg
     3301 acggggagtc aggcaactat ggatgaacga aatagacaga tcgctgagat aggtgcctca
     3361 ctgattaagc attggtaact gtcagaccaa gtttactcat atatacttta gattgattta
     3421 aaacttcatt tttaatttaa aaggatctag gtgaagatcc tttttgataa tctcatgacc
     3481 aaaatccctt aacgtgagtt ttcgttccac tgagcgtcag accccgtaga aaagatcaaa
     3541 ggatcttctt gagatccttt ttttctgcgc gtaatctgct gcttgcaaac aaaaaaacca
     3601 ccgctaccag cggtggtttg tttgccggat caagagctac caactctttt tccgaaggta
     3661 actggcttca gcagagcgca gataccaaat actgtccttc tagtgtagcc gtagttaggc
     3721 caccacttca agaactctgt agcaccgcct acatacctcg ctctgctaat cctgttacca
     3781 gtggctgctg ccagtggcga taagtcgtgt cttaccgggt tggactcaag acgatagtta
     3841 ccggataagg cgcagcggtc gggctgaacg gggggttcgt gcacacagcc cagcttggag
     3901 cgaacgacct acaccgaact gagataccta cagcgtgagc attgagaaag cgccacgctt
     3961 cccgaaggga gaaaggcgga caggtatccg gtaagcggca gggtcggaac aggagagcgc
     4021 acgagggagc ttccaggggg aaacgcctgg tatctttata gtcctgtcgg gtttcgccac
     4081 ctctgacttg agcgtcgatt tttgtgatgc tcgtcagggg ggcggagcct atggaaaaac
     4141 gccagcaacg cggccttttt acggttcctg gccttttgct ggccttttgc tcacatgttc
     4201 tttcctgcgt tatcccctga ttctgtggat aaccgtatta ccgcctttga gtgagctgat
     4261 accgctcgcc gcagccgaac gaccgagcgc agcgagtcag tgagcgagga agcggaagag
     4321 cgcccaatac gcaaaccgcc tctccccgcg cgttggccga ttcattaatg cagctggcac
     4381 gacaggtttc ccgactggaa agcgggcagt gagcgcaacg caattaatgt gagttagctc
     4441 actcattagg caccccaggc tttacacttt atgcttccgg ctcgtatgtt gtgtggaatt
     4501 gtgagcggat aacaatttca cacaggaaac agctatgacc atgattacgc caagcttgat
     4561 aattcctgca gcccagctta attcttttcg agctctttat gcttaagttt acaatttaat
     4621 attcatactt taagtatttt ttgtagtatc ctagatattg tgctttaaat gctcacccct
     4681 caaagcacca gtaatatttt catccactga aataccatta aattttcaaa aaaatactat
     4741 gcatataatg ttatacatat aaacataaaa cgccatgtaa atcaaaaaat atataaaaat
     4801 atgtataaaa ataaatatgc actaaatata agctaattat gcataaaaat taaagtgccc
     4861 tttattaact agctagtcgt aattatttat atttctatgt tataaaaaaa tcctcatata
     4921 ataatataat taatatatgt aatgtttttt ttattttata attttaatat aaaataatat
     4981 gtaaattaat tcaaaaaata aatataattg ttgtgaaaca aaaaacgtaa ttttttcatt
     5041 tgccttcaaa atttaaattt attttaatat ttcctaaaat atatatactt tgtgtataaa
     5101 tatataaaaa tatatatttg cttataaata aataaaaaat tttataaaac atagggggat
     5161 ccatggttgg ttcgctaaac tgcatcgtcg ctgtgtccca gaacatgggc atcggcaaga
     5221 acggggacct gccctggcca ccgctcagga acgaatttag atatttccag agaatgacca
     5281 caacctcttc agtagaaggt aaacagaatc tggtgattat gggtaagaag acctggttct
     5341 ccattcctga gaagaatcga cctttaaagg gtagaattaa tttagttctc agcagagaac
     5401 tcaaggaacc tccacaagga gctcattttc tttccagaag tctagatgat gccttaaaac
     5461 ttactgaaca accagaatta gcaaataaag tagacatggt ctggatagtt ggtggcagtt
     5521 ctgtttataa ggaagccatg aatcacccag gccatcttaa actatttgtg acaaggatca
     5581 tgcaagactt tgaaagtgac acgttttttc cagaaattga tttggagaaa tataaacttc
     5641 tgccagaata cccaggtgtt ctctctgatg tccaggagga gaaaggcatt aagtacaaat
     5701 ttgaagtata tgagaagaat gattaaggat cccgtttttc ttacttatat atttatacca
     5761 attgattgta tttataactg taaaaatgtg tatgttgtgt gcatattttt ttttgtgcat
     5821 gcacatgcat gtaaatagct aaaattatga acattttatt ttttgttcag aaaaaaaaaa
     5881 ctttacacac ataaaatggc tagtatgaat agccatattt tatataaatt aaatcctatg
     5941 aatttatgac catattaaaa atttagatat ttatggaaca taatatgttt gaaacaataa
     6001 gacaaaatta ttattattat tattattttt actgttataa ttatgttgtc tcttcaatga
     6061 ttcataaata gttggacttg atttttaaaa tgtttataat atgattagca tagttaaata
     6121 aaaaaagttg aaaaattaaa aaaaaacata taaacacaaa tgatgttttt tccttcaatt
     6181 tcgatgggta ccCATATTGT TATTTTTAGA ATATTTATAA GTTCGTTTTT ATTATTTTTG
     6241 TTGTAAATAA AAGGTATATA TATTATGTGT ATATAAGCGC GTCTATATGT ACTTACATTA
     6301 TATACATATG CGCGCATACC ATATTTGTGG GATTTTGTTT TTTTCCTCAT AATTAATTTT
     6361 TTAAAGAATA AGCTACACTT TTTCAGTTAA AATTTTATGT ACTACCATCA TTTTTTTTTA
     6421 AATACTTCAA AAGAGAAATA AAAGCAAAAT AATATAAATC CTATAATTGT GTTGTTAGTT
     6481 TGCCGTTTAC ATTTTCATAA ATATGTTTTT GAAATTTCAA TAAATATTGT GTACACATAT
     6541 ATGTAGTATG TATGTATGCA TATGAGAATG AAGTATATTT GTATCATATG CTATGTTTTT
     6601 TTATTTTTTT TTAAGTACAA TTTAATTTTT TTTTTCATTA TATTGTTACA ATAATTTTGG
     6661 ATTTTTATGA AACTTTTAAA CGAAAAGTTA TGCATTTTTC TTTTATTTTT TTAACATCAA
     6721 TTTTTATTAT ATATATATAA GATAAAATTC TTTACTAACA CAATAACATT ATTAAGCATA
     6781 TAAAAATACC AAATTAAATT GAATTAAATT AAATTAAATT AAATTACTAG TTTATAATTT
     6841 CCCAAAATGA AAATACTAAG GGTTTTATTT TTTTAATTAT ACGATTGGCA TATGGATACT
     6901 TAGCATAGAG TGggcggccg cgcgggccc
//

> Py1042_PyNOT1-G::GFP
LOCUS       pSL1042__final_         6754 bp    DNA        circular     21-APR-2021
SOURCE
  ORGANISM
COMMENT     Lindner Lab - used to create Py1042 = Py17XNL PyNOT1-G::GFP
COMMENT
COMMENT     ApEinfo:methylated:1
FEATURES             Location/Qualifiers
     exon            5731..6294
                     /vntifkey="61"
                     /locus_tag="hDHFR"
                     /label="hDHFR"
                     /ApEinfo_label="hDHFR"
                     /ApEinfo_fwdcolor="pink"
                     /ApEinfo_revcolor="pink"
                     /ApEinfo_graphicformat="arrow_data {{0 1 2 0 0 -1} {} 0}
                     width 5 offset 0"
     misc_feature    1512..2190
                     /locus_tag="GFPmut2"
                     /label="GFPmut2"
                     /ApEinfo_label="GFPmut2"
                     /ApEinfo_fwdcolor="#8080c0"
                     /ApEinfo_revcolor="#ff8000"
                     /ApEinfo_graphicformat="arrow_data {{0 1 2 0 0 -1} {} 0}
                     width 5 offset 0"
     misc_feature    2197..2644
                     /locus_tag="PbDHFR/TS\3'UTR(1)"
                     /label="PbDHFR/TS\3'UTR(1)"
                     /ApEinfo_label="PbDHFR/TS\3'UTR(1)"
                     /ApEinfo_fwdcolor="#8080c0"
                     /ApEinfo_revcolor="#ff8000"
                     /ApEinfo_graphicformat="arrow_data {{0 1 2 0 0 -1} {} 0}
                     width 5 offset 0"
     misc_feature    748..1470
                     /locus_tag="PyNOT1-G ORF Homology Arm"
                     /label="PyNOT1-G ORF Homology Arm"
                     /ApEinfo_label="PyNOT1-G ORF Homology Arm"
                     /ApEinfo_fwdcolor="cyan"
                     /ApEinfo_revcolor="green"
                     /ApEinfo_graphicformat="arrow_data {{0 1 2 0 0 -1} {} 0}
                     width 5 offset 0"
     rep_origin      4041..4723
                     /locus_tag="ColE1 origin"
                     /label="ColE1 origin"
                     /ApEinfo_label="ColE1 origin"
                     /ApEinfo_fwdcolor="pink"
                     /ApEinfo_revcolor="pink"
                     /ApEinfo_graphicformat="arrow_data {{0 1 2 0 0 -1} {} 0}
                     width 5 offset 0"
     CDS             3284..3943
                     /locus_tag="AmpR"
                     /label="AmpR"
                     /ApEinfo_label="AmpR"
                     /ApEinfo_fwdcolor="pink"
                     /ApEinfo_revcolor="pink"
                     /ApEinfo_graphicformat="arrow_data {{0 1 2 0 0 -1} {} 0}
                     width 5 offset 0"
     misc_feature    join(5127..5436,5441..5725)
                     /locus_tag="PbEF1a 5'UTR"
                     /label="PbEF1a 5'UTR"
                     /ApEinfo_label="PbEF1a 5'UTR"
                     /ApEinfo_fwdcolor="#8080c0"
                     /ApEinfo_revcolor="#ff8000"
                     /ApEinfo_graphicformat="arrow_data {{0 1 2 0 0 -1} {} 0}
                     width 5 offset 0"
     misc_feature    6302..6749
                     /locus_tag="PbDHFR-TS 3'UTR(1)"
                     /label="PbDHFR-TS 3'UTR(1)"
                     /ApEinfo_label="PbDHFR-TS 3'UTR(1)"
                     /ApEinfo_fwdcolor="#8080c0"
                     /ApEinfo_revcolor="#ff8000"
                     /ApEinfo_graphicformat="arrow_data {{0 1 2 0 0 -1} {} 0}
                     width 5 offset 0"
     misc_feature    7..730
                     /locus_tag="PyNOT1-G 3' Homology Arm"
                     /label="PyNOT1-G 3' Homology Arm"
                     /ApEinfo_label="PyNOT1-G 3' Homology Arm"
                     /ApEinfo_fwdcolor="cyan"
                     /ApEinfo_revcolor="green"
                     /ApEinfo_graphicformat="arrow_data {{0 0.5 0 1 2 0 0 -1 0
                     -0.5} {} 0} width 5 offset 0"
ORIGIN
        1 ggtaccCAAC TCATACAAAC TCTTCTTAGA ATGTAATATA TATGTACATG TATATATATT
       61 ATTATTATTA TTATTTTATT TTTTTTTTTT TTTTTTTTTA CTTACACATG TATGTATGCA
      121 CATACATATA TTACAATTTT ATACCATTTT AAAATTAGTC TTATTCATCA TCGAAGAAAA
      181 ATTTATTATC CTTTGCATAC ATACATACAT ACATATATAT ATATATATAT ACATATATAT
      241 ATACATGCAT TTATTTTACT ATTTTTTTAC AATTTATAAT TGTATTTTTA AAATAGCAGT
      301 TTTGTAATTT TTTTCAAATT CATATTTTTT TTTTCCCATT TTTTTGTTAA ACTTTTAAAA
      361 CACAACAATA GTATCGATAG TACTAGTAGT ATATAATTAT AAAATGCGTT TATTTATTTT
      421 TTCTACATTT CTACAATTTT TCTACAATTT TCCTACAATT TTAAAATACC ACATATAAAC
      481 CTCAATACAC ACATCCATTC CGTTTTAATA AAAAAAAAAA TATATATTTC CGAAATAACG
      541 CTAGGAGGAA AATAAAATTC CCCTGATTTT TACTATCATT TTAATGTTTT CATTATTCTA
      601 TATCTTTAAT AATATACATG AAATGTAATT TTTATGTGTG CAAAAATATT TCAAGTTTAT
      661 TATTAAGCAA AAAATTTACA AAAATATATA CACATATGAG AAATATGAAT CCCGCTTTGT
      721 CTATCCATAC ggcggccgcg cgggcccCTC TCAAATATTT CGAATTCCAT ACAAAATAAC
      781 AAACAACCAA TTTTAGATGA AAATAAAGAT AAAATAAAAG AATTTTATAA TAATATAAAT
      841 ATGAATCAAT TATTGCAAAA TTCTAACGAG AAAAATAATA TTAATAATAC TGATTTAATT
      901 AGGCAAATTC TAAATTCAGA AAATGTTAAT TTAAAAAATT ATACTACAAC TAATGGGCCT
      961 ACTATTACTC CTCATGTTAA ATTCACTAAC CAATATGATA CCAATTCACA CCCAAATGAC
     1021 ATAACAAATA ATATAAAACA AAAACAAAAT GTTATTAATA ATTTACACAC ATATAATATA
     1081 AATTCGAATA ATAACAATAG TAGCAATAAC AATAACAATC ACATTTTGAA TGCTTCGTTT
     1141 TATAGAGATA TTGGAAATTT GAGTGGAATA AAAACAAATA ATAATTCACA AAACAATATC
     1201 GAATTATCAC AAAATAATCT ATTCATTAAT AACCCCTACT ATTTTCAGGA TAAAAATATT
     1261 ATGTTAGATT CACATTTACA AAATAATATT AATCAAATTA AAAGAAATCC AAGATTAAAT
     1321 AACTCACAAA TACATGAAAA TAATGAAAAT GATTCAAATA ATATTCAAAA ACATTTTAAT
     1381 TTCGAAATAA ATAATAAATT AACAGATCAA AATAATAGAG ATTTTTTCAA TTCAAATCAT
     1441 TTTTATGACT CAAAACCTGA TTACTATAAC cctagtagta aaggagaaga acttttcact
     1501 ggagttgtcc cAATTCTTGT TGAATTAGAT GGTGATGTTA ATGGGCACAA ATTTTCTGTC
     1561 AGTGGAGAGG GTGAAGGTGA TGCAACATAC GGAAAACTTA CCCTTAAATT TATTTGCACT
     1621 ACTGGAAAAC TACCTGTTCC ATGGCCAACA CTTGTCACTA CTTTCGCGTA TGGTCTTCAA
     1681 TGCTTTGCGA GATACCCAGA TCATATGAAA CAGCATGACT TTTTCAAGAG TGCCATGCCC
     1741 GAAGGTTATG TACAGGAAAG AACTATATTT TTCAAAGATG ACGGGAACTA CAAGACACGT
     1801 GCTGAAGTCA AGTTTGAAGG TGATACCCTT GTTAATAGAA TCGAGTTAAA AGGTATTGAT
     1861 TTTAAAGAAG ATGGAAACAT TCTTGGACAC AAATTGGAAT ACAACTATAA CTCACACAAT
     1921 GTATACATCA TGGCAGACAA ACAAAAGAAT GGAATCAAAG TTAACTTCAA AATTAGACAC
     1981 AACATTGAAG ATGGAAGCGT TCAACTAGCA GACCATTATC AACAAAATAC TCCAATTGGC
     2041 GATGGCCCTG TCCTTTTACC AGACAACCAT TACCTGTCCA CACAATCTGC CCTTTCGAAA
     2101 GATCCCAACG AAAAGAGAGA CCACATGGTC CTTCTTGAGT TTGTAACAGC TGCTGGGATT
     2161 ACACATGGCA TGGATGAACT ATACAAATAA ggatccGTTT TTCTTACTTA TATATTTATA
     2221 CCAATTGATT GTATTTATAA CTGTAAAAAT GTGTATGTTG TGTGCATATT TTTTTTTGTG
     2281 CATGCACATG CATGTAAATA GCTAAAATTA TGAACATTTT ATTTTTTGTT CAGAAAAAAA
     2341 AAACTTTACA CACATAAAAT GGCTAGTATG AATAGCCATA TTTTATATAA ATTAAATCCT
     2401 ATGAATTTAT GACCATATTA AAAATTTAGA TATTTATGGA ACATAATATG TTTGAAACAA
     2461 TAAGACAAAA TTATTATTAT TATTATTATT TTTACTGTTA TAATTATGTT GTCTCTTCAA
     2521 TGATTCATAA ATAGTTGGAC TTGATTTTTA AAATGTTTAT AATATGATTA GCATAGTTAA
     2581 ATAAAAAAAG TTGAAAAATT AAAAAAAAAC ATATAAACAC AAATGATGTT TTTTCCTTCA
     2641 ATTTcggcgc ctgatgcggt attttctcct tacgcatctg tgcggtattt cacaccgcat
     2701 atggtgcact ctcagtacaa tctgctctga tgccgcatag ttaagccagc cccgacaccc
     2761 gccaacaccc gctgacgcgc cctgacgggc ttgtctgctc ccggcatccg cttacagaca
     2821 agctgtgacc gtctccggga gctgcatgtg tcagaggttt tcaccgtcat caccgaaacg
     2881 cgcgagacga aagggcctcg tgatacgcct atttttatag gttaatgtca tgataataat
     2941 ggtttcttag acgtcaggtg gcacttttcg gggaaatgtg cgcggaaccc ctatttgttt
     3001 atttttctaa atacattcaa atatgtatcc gctcatgaga caataaccct gataaatgct
     3061 tcaataatat tgaaaaagga agagtatgag tattcaacat ttccgtgtcg cccttattcc
     3121 cttttttgcg gcattttgcc ttcctgtttt tgctcaccca gaaacgctgg tgaaagtaaa
     3181 agatgctgaa gatcagttgg gtgcacgagt gggttacatc gaactggatc tcaacagcgg
     3241 taagatcctt gagagttttc gccccgaaga acgttttcca atgatgagca cttttaaagt
     3301 tctgctatgt ggcgcggtat tatcccgtat tgacgccggg caagagcaac tcggtcgccg
     3361 catacactat tctcagaatg acttggttga gtactcacca gtcacagaaa agcatcttac
     3421 ggatggcatg acagtaagag aattatgcag tgctgccata accatgagtg ataacactgc
     3481 ggccaactta cttctgacaa cgatcggagg accgaaggag ctaaccgctt ttttgcacaa
     3541 catgggggat catgtaactc gccttgatcg ttgggaaccg gagctgaatg aagccatacc
     3601 aaacgacgag cgtgacacca cgatgcctgt agcaatggca acaacgttgc gcaaactatt
     3661 aactggcgaa ctacttactc tagcttcccg gcaacaatta atagactgga tggaggcgga
     3721 taaagttgca ggaccacttc tgcgctcggc ccttccggct ggctggttta ttgctgataa
     3781 atctggagcc ggtgagcgtg ggtctcgcgg tatcattgca gcactggggc cagatggtaa
     3841 gccctcccgt atcgtagtta tctacacgac ggggagtcag gcaactatgg atgaacgaaa
     3901 tagacagatc gctgagatag gtgcctcact gattaagcat tggtaactgt cagaccaagt
     3961 ttactcatat atactttaga ttgatttaaa acttcatttt taatttaaaa ggatctaggt
     4021 gaagatcctt tttgataatc tcatgaccaa aatcccttaa cgtgagtttt cgttccactg
     4081 agcgtcagac cccgtagaaa agatcaaagg atcttcttga gatccttttt ttctgcgcgt
     4141 aatctgctgc ttgcaaacaa aaaaaccacc gctaccagcg gtggtttgtt tgccggatca
     4201 agagctacca actctttttc cgaaggtaac tggcttcagc agagcgcaga taccaaatac
     4261 tgtccttcta gtgtagccgt agttaggcca ccacttcaag aactctgtag caccgcctac
     4321 atacctcgct ctgctaatcc tgttaccagt ggctgctgcc agtggcgata agtcgtgtct
     4381 taccgggttg gactcaagac gatagttacc ggataaggcg cagcggtcgg gctgaacggg
     4441 gggttcgtgc acacagccca gcttggagcg aacgacctac accgaactga gatacctaca
     4501 gcgtgagcat tgagaaagcg ccacgcttcc cgaagggaga aaggcggaca ggtatccggt
     4561 aagcggcagg gtcggaacag gagagcgcac gagggagctt ccagggggaa acgcctggta
     4621 tctttatagt cctgtcgggt ttcgccacct ctgacttgag cgtcgatttt tgtgatgctc
     4681 gtcagggggg cggagcctat ggaaaaacgc cagcaacgcg gcctttttac ggttcctggc
     4741 cttttgctgg ccttttgctc acatgttctt tcctgcgtta tcccctgatt ctgtggataa
     4801 ccgtattacc gcctttgagt gagctgatac cgctcgccgc agccgaacga ccgagcgcag
     4861 cgagtcagtg agcgaggaag cggaagagcg cccaatacgc aaaccgcctc tccccgcgcg
     4921 ttggccgatt cattaatgca gctggcacga caggtttccc gactggaaag cgggcagtga
     4981 gcgcaacgca attaatgtga gttagctcac tcattaggca ccccaggctt tacactttat
     5041 gcttccggct cgtatgttgt gtggaattgt gagcggataa caatttcaca caggaaacag
     5101 ctatgaccat gattacgcca agcttgataa ttcctgcagc ccagcttaat tcttttcgag
     5161 ctctttatgc ttaagtttac aatttaatat tcatacttta agtatttttt gtagtatcct
     5221 agatattgtg ctttaaatgc tcacccctca aagcaccagt aatattttca tccactgaaa
     5281 taccattaaa ttttcaaaaa aatactatgc atataatgtt atacatataa acataaaacg
     5341 ccatgtaaat caaaaaatat ataaaaatat gtataaaaat aaatatgcac taaatataag
     5401 ctaattatgc ataaaaatta aagtgccctt tattaactag ctagtcgtaa ttatttatat
     5461 ttctatgtta taaaaaaatc ctcatataat aatataatta atatatgtaa tgtttttttt
     5521 attttataat tttaatataa aataatatgt aaattaattc aaaaaataaa tataattgtt
     5581 gtgaaacaaa aaacgtaatt ttttcatttg ccttcaaaat ttaaatttat tttaatattt
     5641 cctaaaatat atatactttg tgtataaata tataaaaata tatatttgct tataaataaa
     5701 taaaaaattt tataaaacat agggggatcc atggttggtt cgctaaactg catcgtcgct
     5761 gtgtcccaga acatgggcat cggcaagaac ggggacctgc cctggccacc gctcaggaac
     5821 gaatttagat atttccagag aatgaccaca acctcttcag tagaaggtaa acagaatctg
     5881 gtgattatgg gtaagaagac ctggttctcc attcctgaga agaatcgacc tttaaagggt
     5941 agaattaatt tagttctcag cagagaactc aaggaacctc cacaaggagc tcattttctt
     6001 tccagaagtc tagatgatgc cttaaaactt actgaacaac cagaattagc aaataaagta
     6061 gacatggtct ggatagttgg tggcagttct gtttataagg aagccatgaa tcacccaggc
     6121 catcttaaac tatttgtgac aaggatcatg caagactttg aaagtgacac gttttttcca
     6181 gaaattgatt tggagaaata taaacttctg ccagaatacc caggtgttct ctctgatgtc
     6241 caggaggaga aaggcattaa gtacaaattt gaagtatatg agaagaatga ttaaggatcc
     6301 cgtttttctt acttatatat ttataccaat tgattgtatt tataactgta aaaatgtgta
     6361 tgttgtgtgc atattttttt ttgtgcatgc acatgcatgt aaatagctaa aattatgaac
     6421 attttatttt ttgttcagaa aaaaaaaact ttacacacat aaaatggcta gtatgaatag
     6481 ccatatttta tataaattaa atcctatgaa tttatgacca tattaaaaat ttagatattt
     6541 atggaacata atatgtttga aacaataaga caaaattatt attattatta ttatttttac
     6601 tgttataatt atgttgtctc ttcaatgatt cataaatagt tggacttgat ttttaaaatg
     6661 tttataatat gattagcata gttaaataaa aaaagttgaa aaattaaaaa aaaacatata
     6721 aacacaaatg atgttttttc cttcaatttc gatg
//
```

```
> Py1023_PyNOT1::GFP
LOCUS       pSL1023__final_         6767 bp    DNA        circular     21-APR-2021
SOURCE
  ORGANISM
COMMENT     Lindner Lab - used to create Py1023 = Py17XNL PyNOT1::GFP
COMMENT
COMMENT     ApEinfo:methylated:1
FEATURES             Location/Qualifiers
     exon            5744..6307
                     /vntifkey="61"
                     /locus_tag="hDHFR"
                     /label="hDHFR"
                     /ApEinfo_label="hDHFR"
                     /ApEinfo_fwdcolor="pink"
                     /ApEinfo_revcolor="pink"
                     /ApEinfo_graphicformat="arrow_data {{0 1 2 0 0 -1} {} 0}
                     width 5 offset 0"
     misc_feature    1525..2203
                     /locus_tag="GFPmut2"
                     /label="GFPmut2"
                     /ApEinfo_label="GFPmut2"
                     /ApEinfo_fwdcolor="#8080c0"
                     /ApEinfo_revcolor="#ff8000"
                     /ApEinfo_graphicformat="arrow_data {{0 1 2 0 0 -1} {} 0}
                     width 5 offset 0"
     misc_feature    2210..2657
                     /locus_tag="PbDHFR/TS\3'UTR"
                     /label="PbDHFR/TS\3'UTR"
                     /ApEinfo_label="PbDHFR/TS\3'UTR"
                     /ApEinfo_fwdcolor="#8080c0"
                     /ApEinfo_revcolor="#ff8000"
                     /ApEinfo_graphicformat="arrow_data {{0 1 2 0 0 -1} {} 0}
                     width 5 offset 0"
     misc_feature    744..1478
                     /locus_tag="PyNOT1 ORF Homology Arm"
                     /label="PyNOT1 ORF Homology Arm"
                     /ApEinfo_label="PyNOT1 ORF Homology Arm"
                     /ApEinfo_fwdcolor="cyan"
                     /ApEinfo_revcolor="green"
                     /ApEinfo_graphicformat="arrow_data {{0 1 2 0 0 -1} {} 0}
                     width 5 offset 0"
     rep_origin      4054..4736
                     /locus_tag="ColE1 origin"
                     /label="ColE1 origin"
                     /ApEinfo_label="ColE1 origin"
                     /ApEinfo_fwdcolor="pink"
                     /ApEinfo_revcolor="pink"
                     /ApEinfo_graphicformat="arrow_data {{0 1 2 0 0 -1} {} 0}
                     width 5 offset 0"
     CDS             3297..3956
                     /locus_tag="AmpR"
                     /label="AmpR"
                     /ApEinfo_label="AmpR"
                     /ApEinfo_fwdcolor="pink"
                     /ApEinfo_revcolor="pink"
                     /ApEinfo_graphicformat="arrow_data {{0 1 2 0 0 -1} {} 0}
                     width 5 offset 0"
     misc_feature    join(5140..5449,5454..5738)
                     /locus_tag="PbEF1a 5'UTR"
                     /label="PbEF1a 5'UTR"
                     /ApEinfo_label="PbEF1a 5'UTR"
                     /ApEinfo_fwdcolor="#8080c0"
                     /ApEinfo_revcolor="#ff8000"
                     /ApEinfo_graphicformat="arrow_data {{0 1 2 0 0 -1} {} 0}
                     width 5 offset 0"
     misc_feature    6315..6762
                     /locus_tag="PbDHFR-TS 3'UTR"
                     /label="PbDHFR-TS 3'UTR"
                     /ApEinfo_label="PbDHFR-TS 3'UTR"
                     /ApEinfo_fwdcolor="#8080c0"
                     /ApEinfo_revcolor="#ff8000"
                     /ApEinfo_graphicformat="arrow_data {{0 1 2 0 0 -1} {} 0}
                     width 5 offset 0"
     misc_feature    7..726
                     /locus_tag="PyNOT1 3' Homology Arm"
                     /label="PyNOT1 3' Homology Arm"
                     /ApEinfo_label="PyNOT1 3' Homology Arm"
                     /ApEinfo_fwdcolor="cyan"
                     /ApEinfo_revcolor="green"
                     /ApEinfo_graphicformat="arrow_data {{0 0.5 0 1 2 0 0 -1 0
                     -0.5} {} 0} width 5 offset 0"
ORIGIN
        1 ggtaccCATA TTGTTATTTT TAGAATATTT ATAAGTTCGT TTTTATTATT TTTGTTGTAA
       61 ATAAAAGGTA TATATATTAT GTGTATATAA GCGCGTCTAT ATGTACTTAC ATTATATACA
      121 TATGCGCGCA TACCATATTT GTGGGATTTT GTTTTTTTCC TCATAATTAA TTTTTTAAAG
      181 AATAAGCTAC ACTTTTTCAG TTAAAATTTT ATGTACTACC ATCATTTTTT TTTAAATACT
      241 TCAAAAGAGA AATAAAAGCA AAATAATATA AATCCTATAA TTGTGTTGTT AGTTTGCCGT
      301 TTACATTTTC ATAAATATGT TTTTGAAATT TCAATAAATA TTGTGTACAC ATATATGTAG
      361 TATGTATGTA TGCATATGAG AATGAAGTAT ATTTGTATCA TATGCTATGT TTTTTTATTT
      421 TTTTTTAAGT ACAATTTAAT TTTTTTTTTC ATTATATTGT TACAATAATT TTGGATTTTT
      481 ATGAAACTTT TAAACGAAAA GTTATGCATT TTTCTTTTAT TTTTTTAACA TCAATTTTTA
      541 TTATATATAT ATAAGATAAA ATTCTTTACT AACACAATAA CATTATTAAG CATATAAAAA
      601 TACCAAATTA AATTGAATTA AATTAAATTA AATTAAATTA CTAGTTTATA ATTTCCCAAA
      661 ATGAAAATAC TAAGGGTTTT ATTTTTTTAA TTATACGATT GGCATATGGA TACTTAGCAT
      721 AGAGTGggcg gccgcgcggg cccCAAGGAG AAATTTTAAA ACGAAAAGAC ATAAAAGAAG
      781 GAGATGAAAA ACAACCAAAT CAAGAAAATC AAAGAAATGG AGAACAAAAT AATAATTTGC
      841 CTAACAAATA TGTAGATCTT GTGCAAGAAA ATAATATAGA AAATGAAATT TCATCATTTT
      901 CAAATTGTGA TAAAACAAAT ACTAAAGTGT TAAAACAAAA TAGTGTAAAT AATAAAGAAG
      961 ATAATGATGG CACAATAGAA ATTACTATTA TCAAAAAAAA TCTTGCTTAT ACATTATTTT
     1021 TGTTTTTATT AAAAGAATTA GATATGGAAG GAAGATACTT ATTATTATTG AATATAGTTA
     1081 ATCATATTAG ATATCCTAAT TCGCATACCC ATTATTTTTC ATGTCTCATT TTGTTTTTAT
     1141 TTTCATATTC AAATGATATC GTTATTAAAG AACAAATAAT TCGGGTTTTG CTGGAAAGAA
     1201 TTTTAGCTCA TAGGCCACAT CCATGGGGAT TATTAATTAC ATTTATTGAA TTAATCAAAA
     1261 ATAAAAAGTA AGAGAATATT TATTTTTTAC ACAAACAAAA AGAGGATATA CATAAATAAG
     1321 CATGTATATA ATGTATATAC TAAAATACAC ATTATTTACA TATTAATATA ATACCTTTTT
     1381 TTTATAGATT CAAAATTTGG GAATATCCAT TTGTACATGC CACATCGGAG ATTAAAAAAA
     1441 TATTCAAGTC AGTTTTTCAG ACATGCTTAG GAAATGTTcc taggctagta gtaaaggaga
     1501 agaacttttc actggagttg tcccAATTCT TGTTGAATTA GATGGTGATG TTAATGGGCA
     1561 CAAATTTTCT GTCAGTGGAG AGGGTGAAGG TGATGCAACA TACGGAAAAC TTACCCTTAA
     1621 ATTTATTTGC ACTACTGGAA AACTACCTGT TCCATGGCCA ACACTTGTCA CTACTTTCGC
     1681 GTATGGTCTT CAATGCTTTG CGAGATACCC AGATCATATG AAACAGCATG ACTTTTTCAA
     1741 GAGTGCCATG CCCGAAGGTT ATGTACAGGA AAGAACTATA TTTTTCAAAG ATGACGGGAA
     1801 CTACAAGACA CGTGCTGAAG TCAAGTTTGA AGGTGATACC CTTGTTAATA GAATCGAGTT
     1861 AAAAGGTATT GATTTTAAAG AAGATGGAAA CATTCTTGGA CACAAATTGG AATACAACTA
     1921 TAACTCACAC AATGTATACA TCATGGCAGA CAAACAAAAG AATGGAATCA AAGTTAACTT
     1981 CAAAATTAGA CACAACATTG AAGATGGAAG CGTTCAACTA GCAGACCATT ATCAACAAAA
     2041 TACTCCAATT GGCGATGGCC CTGTCCTTTT ACCAGACAAC CATTACCTGT CCACACAATC
     2101 TGCCCTTTCG AAAGATCCCA ACGAAAAGAG AGACCACATG GTCCTTCTTG AGTTTGTAAC
     2161 AGCTGCTGGG ATTACACATG GCATGGATGA ACTATACAAA TAAggatccG TTTTTCTTAC
     2221 TTATATATTT ATACCAATTG ATTGTATTTA TAACTGTAAA AATGTGTATG TTGTGTGCAT
     2281 ATTTTTTTTT GTGCATGCAC ATGCATGTAA ATAGCTAAAA TTATGAACAT TTTATTTTTT
     2341 GTTCAGAAAA AAAAAACTTT ACACACATAA AATGGCTAGT ATGAATAGCC ATATTTTATA
     2401 TAAATTAAAT CCTATGAATT TATGACCATA TTAAAAATTT AGATATTTAT GGAACATAAT
     2461 ATGTTTGAAA CAATAAGACA AAATTATTAT TATTATTATT ATTTTTACTG TTATAATTAT
     2521 GTTGTCTCTT CAATGATTCA TAAATAGTTG GACTTGATTT TTAAAATGTT TATAATATGA
     2581 TTAGCATAGT TAAATAAAAA AAGTTGAAAA ATTAAAAAAA AACATATAAA CACAAATGAT
     2641 GTTTTTTCCT TCAATTTcgg cgcctgatgc ggtattttct ccttacgcat ctgtgcggta
     2701 tttcacaccg catatggtgc actctcagta caatctgctc tgatgccgca tagttaagcc
     2761 agccccgaca cccgccaaca cccgctgacg cgccctgacg ggcttgtctg ctcccggcat
     2821 ccgcttacag acaagctgtg accgtctccg ggagctgcat gtgtcagagg ttttcaccgt
     2881 catcaccgaa acgcgcgaga cgaaagggcc tcgtgatacg cctattttta taggttaatg
     2941 tcatgataat aatggtttct tagacgtcag gtggcacttt tcggggaaat gtgcgcggaa
     3001 cccctatttg tttatttttc taaatacatt caaatatgta tccgctcatg agacaataac
     3061 cctgataaat gcttcaataa tattgaaaaa ggaagagtat gagtattcaa catttccgtg
     3121 tcgcccttat tccctttttt gcggcatttt gccttcctgt ttttgctcac ccagaaacgc
     3181 tggtgaaagt aaaagatgct gaagatcagt tgggtgcacg agtgggttac atcgaactgg
     3241 atctcaacag cggtaagatc cttgagagtt ttcgccccga agaacgtttt ccaatgatga
     3301 gcacttttaa agttctgcta tgtggcgcgg tattatcccg tattgacgcc gggcaagagc
     3361 aactcggtcg ccgcatacac tattctcaga atgacttggt tgagtactca ccagtcacag
     3421 aaaagcatct tacggatggc atgacagtaa gagaattatg cagtgctgcc ataaccatga
     3481 gtgataacac tgcggccaac ttacttctga caacgatcgg aggaccgaag gagctaaccg
     3541 cttttttgca caacatgggg gatcatgtaa ctcgccttga tcgttgggaa ccggagctga
     3601 atgaagccat accaaacgac gagcgtgaca ccacgatgcc tgtagcaatg gcaacaacgt
     3661 tgcgcaaact attaactggc gaactactta ctctagcttc ccggcaacaa ttaatagact
     3721 ggatggaggc ggataaagtt gcaggaccac ttctgcgctc ggcccttccg gctggctggt
     3781 ttattgctga taaatctgga gccggtgagc gtgggtctcg cggtatcatt gcagcactgg
     3841 ggccagatgg taagccctcc cgtatcgtag ttatctacac gacggggagt caggcaacta
     3901 tggatgaacg aaatagacag atcgctgaga taggtgcctc actgattaag cattggtaac
     3961 tgtcagacca agtttactca tatatacttt agattgattt aaaacttcat ttttaattta
     4021 aaaggatcta ggtgaagatc ctttttgata atctcatgac caaaatccct taacgtgagt
     4081 tttcgttcca ctgagcgtca gaccccgtag aaaagatcaa aggatcttct tgagatcctt
     4141 tttttctgcg cgtaatctgc tgcttgcaaa caaaaaaacc accgctacca gcggtggttt
     4201 gtttgccgga tcaagagcta ccaactcttt ttccgaaggt aactggcttc agcagagcgc
     4261 agataccaaa tactgtcctt ctagtgtagc cgtagttagg ccaccacttc aagaactctg
     4321 tagcaccgcc tacatacctc gctctgctaa tcctgttacc agtggctgct gccagtggcg
     4381 ataagtcgtg tcttaccggg ttggactcaa gacgatagtt accggataag gcgcagcggt
     4441 cgggctgaac ggggggttcg tgcacacagc ccagcttgga gcgaacgacc tacaccgaac
     4501 tgagatacct acagcgtgag cattgagaaa gcgccacgct tcccgaaggg agaaaggcgg
     4561 acaggtatcc ggtaagcggc agggtcggaa caggagagcg cacgagggag cttccagggg
     4621 gaaacgcctg gtatctttat agtcctgtcg ggtttcgcca cctctgactt gagcgtcgat
     4681 ttttgtgatg ctcgtcaggg gggcggagcc tatggaaaaa cgccagcaac gcggcctttt
     4741 tacggttcct ggccttttgc tggccttttg ctcacatgtt ctttcctgcg ttatcccctg
     4801 attctgtgga taaccgtatt accgcctttg agtgagctga taccgctcgc cgcagccgaa
     4861 cgaccgagcg cagcgagtca gtgagcgagg aagcggaaga gcgcccaata cgcaaaccgc
     4921 ctctccccgc gcgttggccg attcattaat gcagctggca cgacaggttt cccgactgga
     4981 aagcgggcag tgagcgcaac gcaattaatg tgagttagct cactcattag gcaccccagg
     5041 ctttacactt tatgcttccg gctcgtatgt tgtgtggaat tgtgagcgga taacaatttc
     5101 acacaggaaa cagctatgac catgattacg ccaagcttga taattcctgc agcccagctt
     5161 aattcttttc gagctcttta tgcttaagtt tacaatttaa tattcatact ttaagtattt
     5221 tttgtagtat cctagatatt gtgctttaaa tgctcacccc tcaaagcacc agtaatattt
     5281 tcatccactg aaataccatt aaattttcaa aaaaatacta tgcatataat gttatacata
     5341 taaacataaa acgccatgta aatcaaaaaa tatataaaaa tatgtataaa aataaatatg
     5401 cactaaatat aagctaatta tgcataaaaa ttaaagtgcc ctttattaac tagctagtcg
     5461 taattattta tatttctatg ttataaaaaa atcctcatat aataatataa ttaatatatg
     5521 taatgttttt tttattttat aattttaata taaaataata tgtaaattaa ttcaaaaaat
     5581 aaatataatt gttgtgaaac aaaaaacgta attttttcat ttgccttcaa aatttaaatt
     5641 tattttaata tttcctaaaa tatatatact ttgtgtataa atatataaaa atatatattt
     5701 gcttataaat aaataaaaaa ttttataaaa cataggggga tccatggttg gttcgctaaa
     5761 ctgcatcgtc gctgtgtccc agaacatggg catcggcaag aacggggacc tgccctggcc
     5821 accgctcagg aacgaattta gatatttcca gagaatgacc acaacctctt cagtagaagg
     5881 taaacagaat ctggtgatta tgggtaagaa gacctggttc tccattcctg agaagaatcg
     5941 acctttaaag ggtagaatta atttagttct cagcagagaa ctcaaggaac ctccacaagg
     6001 agctcatttt ctttccagaa gtctagatga tgccttaaaa cttactgaac aaccagaatt
     6061 agcaaataaa gtagacatgg tctggatagt tggtggcagt tctgtttata aggaagccat
     6121 gaatcaccca ggccatctta aactatttgt gacaaggatc atgcaagact ttgaaagtga
     6181 cacgtttttt ccagaaattg atttggagaa atataaactt ctgccagaat acccaggtgt
     6241 tctctctgat gtccaggagg agaaaggcat taagtacaaa tttgaagtat atgagaagaa
     6301 tgattaagga tcccgttttt cttacttata tatttatacc aattgattgt atttataact
     6361 gtaaaaatgt gtatgttgtg tgcatatttt tttttgtgca tgcacatgca tgtaaatagc
     6421 taaaattatg aacattttat tttttgttca gaaaaaaaaa actttacaca cataaaatgg
     6481 ctagtatgaa tagccatatt ttatataaat taaatcctat gaatttatga ccatattaaa
     6541 aatttagata tttatggaac ataatatgtt tgaaacaata agacaaaatt attattatta
     6601 ttattatttt tactgttata attatgttgt ctcttcaatg attcataaat agttggactt
     6661 gatttttaaa atgtttataa tatgattagc atagttaaat aaaaaaagtt gaaaaattaa
     6721 aaaaaaacat ataaacacaa atgatgtttt ttccttcaat ttcgatg
//
```

```
>Py1125_PyNOT1-G_TTPbd::GFP_Overexpression
LOCUS       pSL1125__final_         8361 bp    DNA        circular     21-APR-2021
SOURCE
  ORGANISM
COMMENT     Lindner Lab - used to create Py1125 = Py17XNL Overexpression of
            PyNOT1-G's TTP-binding domain fused to GFP, expressed from p230p
            locus
COMMENT     ApEinfo:methylated:1
FEATURES             Location/Qualifiers
     exon            7338..7901
                     /vntifkey="61"
                     /locus_tag="hDHFR"
                     /label="hDHFR"
                     /ApEinfo_label="hDHFR"
                     /ApEinfo_fwdcolor="pink"
                     /ApEinfo_revcolor="pink"
                     /ApEinfo_graphicformat="arrow_data {{0 1 2 0 0 -1} {} 0}
                     width 5 offset 0"
     misc_feature    2481..3077
                     /locus_tag="PyNOT1-G AA1-199"
                     /label="PyNOT1-G AA1-199"
                     /ApEinfo_label="PyNOT1-G AA1-199"
                     /ApEinfo_fwdcolor="cyan"
                     /ApEinfo_revcolor="green"
                     /ApEinfo_graphicformat="arrow_data {{0 1 2 0 0 -1} {} 0}
                     width 5 offset 0"
     misc_feature    1898..2471
                     /locus_tag="PbEF1a promoter"
                     /label="PbEF1a promoter"
                     /ApEinfo_label="PbEF1a promoter"
                     /ApEinfo_fwdcolor="cyan"
                     /ApEinfo_revcolor="green"
                     /ApEinfo_graphicformat="arrow_data {{0 1 2 0 0 -1} {} 0}
                     width 5 offset 0"
     misc_feature    3119..3797
                     /locus_tag="GFPmut2"
                     /label="GFPmut2"
                     /ApEinfo_label="GFPmut2"
                     /ApEinfo_fwdcolor="cyan"
                     /ApEinfo_revcolor="green"
                     /ApEinfo_graphicformat="arrow_data {{0 1 2 0 0 -1} {} 0}
                     width 5 offset 0"
     misc_feature    3804..4251
                     /locus_tag="PbDHFR/TS\3'UTR"
                     /label="PbDHFR/TS\3'UTR"
                     /ApEinfo_label="PbDHFR/TS\3'UTR"
                     /ApEinfo_fwdcolor="#ff0000"
                     /ApEinfo_revcolor="green"
                     /ApEinfo_graphicformat="arrow_data {{0 1 2 0 0 -1} {} 0}
                     width 5 offset 0"
     rep_origin      5648..6330
                     /locus_tag="ColE1 origin"
                     /label="ColE1 origin"
                     /ApEinfo_label="ColE1 origin"
                     /ApEinfo_fwdcolor="gray50"
                     /ApEinfo_revcolor="gray50"
                     /ApEinfo_graphicformat="arrow_data {{0 1 2 0 0 -1} {} 0}
                     width 5 offset 0"
     CDS             4891..5550
                     /locus_tag="AmpR"
                     /label="AmpR"
                     /ApEinfo_label="AmpR"
                     /ApEinfo_fwdcolor="yellow"
                     /ApEinfo_revcolor="yellow"
                     /ApEinfo_graphicformat="arrow_data {{0 1 2 0 0 -1} {} 0}
                     width 5 offset 0"
     misc_feature    join(6734..7043,7048..7332)
                     /locus_tag="PbEF1a 5'UTR"
                     /label="PbEF1a 5'UTR"
                     /ApEinfo_label="PbEF1a 5'UTR"
                     /ApEinfo_fwdcolor="#804040"
                     /ApEinfo_revcolor="#804040"
                     /ApEinfo_graphicformat="arrow_data {{0 1 2 0 0 -1} {} 0}
                     width 5 offset 0"
     misc_feature    7909..8356
                     /locus_tag="PbDHFR-TS 3'UTR"
                     /label="PbDHFR-TS 3'UTR"
                     /ApEinfo_label="PbDHFR-TS 3'UTR"
                     /ApEinfo_fwdcolor="cyan"
                     /ApEinfo_revcolor="green"
                     /ApEinfo_graphicformat="arrow_data {{0 1 2 0 0 -1} {} 0}
                     width 5 offset 0"
     misc_feature    7..918
                     /locus_tag="py p230p Homology Arm 1"
                     /label="py p230p Homology Arm 1"
                     /ApEinfo_label="py p230p Homology Arm 1"
                     /ApEinfo_fwdcolor="cyan"
                     /ApEinfo_revcolor="green"
                     /ApEinfo_graphicformat="arrow_data {{0 0.5 0 1 2 0 0 -1 0
                     -0.5} {} 0} width 5 offset 0"
     misc_feature    925..1873
                     /locus_tag="py p230p Homology Arm 2"
                     /label="py p230p Homology Arm 2"
                     /ApEinfo_label="py p230p Homology Arm 2"
                     /ApEinfo_fwdcolor="cyan"
                     /ApEinfo_revcolor="green"
                     /ApEinfo_graphicformat="arrow_data {{0 0.5 0 1 2 0 0 -1 0
                     -0.5} {} 0} width 5 offset 0"
ORIGIN
        1 ggtaccAGCT TTGTGTTTTA TTTGGATGTG CAACAATAGA TTTATGATTA ACCAATGGTA
       61 AATCATTATA ATCATATATT TCATCTTCAT TTACATATTT TACTAAAACA GTTTTTACAA
      121 TTTGTTCTGT TTCTTTATCT ACACAATGAC ATTCAAATTG ATATGATTTT GTTAATGTAT
      181 AATTAAAACC AGCTAATGCA TATTCTGGAT TTTCAAATGA ATAGTTAATA ATATTTTTAG
      241 TATTAATTTT ATAGGAATCA TGTAGATAGC CTTTTTCATA TACTTGAGTA AAACATTTTT
      301 CTGGCATTTT TTTTGTATTT TCGGGGCATC TAAAACCAAA ACTTGTATTT GATTTGGAAA
      361 TTATTATACA TAGATCTCCT TCACTATAAG CATTTAAATA ATTATATAAT TCACCTTTTC
      421 TAAAATCACA AAATGAAACT GGTCTATCTT TTAAATCTGT TCGAACATGT AATAAAATAA
      481 TTGCTCTTCT ATAATCAGTC CCACCATTTG TTACACCACA TTCTATGGTT AGTGGATATG
      541 ATCCATGATT TATAAATGAG GTCATAAAAA ATTTAAAACC TTTAAATTGT TCATTTATTT
      601 CTGGTATATC TATATCTTTT GTATTAAAAA ATAAATCTGT AAATTTATTT ATAGGAAATA
      661 CATTACTTAA ATATTTTATG GTATTAATAG GTTCTTTACC ATATACGATT CTATTATCTG
      721 TTATTTGCTC ACAATTATAC ATGAATAAAC TAAATTCATT TATTGTTTTT TCACATATTT
      781 GTATTTCACT TAATTTGCTT AAATTATAAT CAAAATTACA CATCTGATAA GGTATATCGT
      841 TAAAATATTT AATCGTTTTA CTATCATCAT CAACTATTTT TAAAATGTGA TCAGATGTTG
      901 CCATTCCATC ACCAAATGgg gcccTATGGA ACTACATCTA TATAAGAGAT TTTTTTATTT
      961 TTATATATTG GTTTTGAATC TAGATATTCA TAATTTTTAA TATATATTGT TCCTATATTT
     1021 TTGGTTTTAT AATCTTCACA TGTACAATAA AAATCAAATT CTTTTTTTGT AATTTTTGAA
     1081 ATCATTAAAA ATGTTGATGA ATTATTTATA AAATAATCTG CATCTATAAC TTTTATTTCA
     1141 TTAAATATAT TTTCGATATT TTCAGTTATT TGTTCATTCA TGTTACTTTT TATATATACC
     1201 TTTCTAAAAC AATTTTTTGG ATTAATTTTT CCAGTTGGAC AATTTAAGCT AACTACTATA
     1261 TTTTCTTTTA ATTCGATTTC ACAAAATTCA TTTTGTTTTT CTAAATCATA AACATTGGAA
     1321 TAGTAAAATT TTTTTTTAGA ATTACCTGAA AAATCACATC CATATAATTT ATTTTTTGTT
     1381 GTATTTTGAG AAACCCTTAT CACTGTAATA CCTTTTTGAC CTTTCATATT ATTAATTATT
     1441 GTTTTAGTAT TATCACAATA TATTTTCAAT ATTAATACTT TTTCTACAAT TAATGGAGTA
     1501 ACAAAAGTGA AATTTCCTTT CTCTTTGAAC ATTTTATTTT TATCTAATAT GATAACACCT
     1561 GGTAATATAT CATCTAATTT TTCTTTTTTG TCTAGATACA TTACTTCATC ATTTTCTGGT
     1621 ATGTATTCTA TATTTTTATA TTTTTCTAAT TCTTCTGGTT CAACATTTGC ATTTTCTAAA
     1681 TAAAAATTAT TTTTTTGACT ACTACGTTGT GATGTTTTCT TATTATTTAT TATGATATTA
     1741 CTATCAAAAA TGTCATCATC ACTTTTTACA ATTTTGGGAC ATTTAACTTG AACCATATTC
     1801 AGATTCTGTT CTAGATGTAT ATAACATGAA TGAACTGTAT CTACTTCATT AACGGGTTCA
     1861 AGAGAATTCG GTAgccgcgg tggcggccgc TctcgagAGC TTAATTCTTT TCGAGCTCTT
     1921 TATGCTTAAG TTTACAATTT AATATTCATA CTTTAAGTAT TTTTTGTAGT ATCCTAGATA
     1981 TTGTGCTTTA AATGCTCACC CCTCAAAGCA CCAGTAATAT TTTCATCCAC TGAAATACCA
     2041 TTAAATTTTC AAAAAAATAC TATGCATATA ATGTTATACA TATAAACATA AAACGCCATG
     2101 TAAATCAAAA AATATATAAA AATATGTATA AAAATAAATA TGCACTAAAT ATAAGCTAAT
     2161 TATGCATAAA AATTAAAGTG CCCTTTATTA ACTAGctagT CGTAATTATT TATATTTCTA
     2221 TGTTATAAAA AAATCCTCAT ATAATAATAT AATTAATATA TGTAATGTTT TTTTTATTTT
     2281 ATAATTTTAA TATAAAATAA TATGTAAATT AATTCAAAAA ATAAATATAA TTGTTGTGAA
     2341 ACAAAAAACG TAATTTTTTC ATTTGCCTTC AAAATTTAAA TTTATTTTAA TATTTCCTAA
     2401 AATATATATA CTTTGTGTAT AAATATATAA AAATATATAT TTGCTTATAA ATAAATAAAA
     2461 ATTTTATAAA Aatgactagg ATGAATAACA ATTTTAACAT TAATCTTCAG ATCGAGGATG
     2521 GAATCACCAA TAAATATGAA GCCGAAGTAA ATGGCTATTT TGCTAAATTG TACACAGGGG
     2581 AAATAACAGT CAACACAATG ATAGATATCA TGAAAAATCT ATCATGCTCC CCTAAAGGAT
     2641 CGAAAAATAA CGATATTTAT AAATCGATGC TATTAATCTT ATTTAATGAG TGCAAATTCT
     2701 TTCCTAAATA TCCAGTAGAA GAATTAGATA TAACTGCACA ATTGTTTGGA AAATTAATTA
     2761 AGCACAATTT ATTAATATCA TATGGTAATA CTTTATCTGT TGTATTAAAA TGTATTTTAG
     2821 AAGCTTTAAA AAAAGGATCT GATTCTAAAG TTTTTAATTT TGGGATAACA GCATTAGAGC
     2881 AATTTGAAGA CTCTTTAATA TGTTACCCTG CCTTTTTATC TTCCTTAATT CCATTACCTA
     2941 CTTTAAGGCA ATATAATCCG CAATATATAA TCCATTGCAA TGAATTATTA AACACACTTC
     3001 CAGAACAGTT TAGAACTCTT CCATATATAG ATGCTTCTAC AATACTTAAG ATAAAACACA
     3061 TCTCTGAAAT TAGTTCTcct agtagtaaag gagaagaact tttcactgga gttgtcccAA
     3121 TTCTTGTTGA ATTAGATGGT GATGTTAATG GGCACAAATT TTCTGTCAGT GGAGAGGGTG
     3181 AAGGTGATGC AACATACGGA AAACTTACCC TTAAATTTAT TTGCACTACT GGAAAACTAC
     3241 CTGTTCCATG GCCAACACTT GTCACTACTT TCGCGTATGG TCTTCAATGC TTTGCGAGAT
     3301 ACCCAGATCA TATGAAACAG CATGACTTTT TCAAGAGTGC CATGCCCGAA GGTTATGTAC
     3361 AGGAAAGAAC TATATTTTTC AAAGATGACG GGAACTACAA GACACGTGCT GAAGTCAAGT
     3421 TTGAAGGTGA TACCCTTGTT AATAGAATCG AGTTAAAAGG TATTGATTTT AAAGAAGATG
     3481 GAAACATTCT TGGACACAAA TTGGAATACA ACTATAACTC ACACAATGTA TACATCATGG
     3541 CAGACAAACA AAAGAATGGA ATCAAAGTTA ACTTCAAAAT TAGACACAAC ATTGAAGATG
     3601 GAAGCGTTCA ACTAGCAGAC CATTATCAAC AAAATACTCC AATTGGCGAT GGCCCTGTCC
     3661 TTTTACCAGA CAACCATTAC CTGTCCACAC AATCTGCCCT TTCGAAAGAT CCCAACGAAA
     3721 AGAGAGACCA CATGGTCCTT CTTGAGTTTG TAACAGCTGC TGGGATTACA CATGGCATGG
     3781 ATGAACTATA CAAATAAgga tccGTTTTTC TTACTTATAT ATTTATACCA ATTGATTGTA
     3841 TTTATAACTG TAAAAATGTG TATGTTGTGT GCATATTTTT TTTTGTGCAT GCACATGCAT
     3901 GTAAATAGCT AAAATTATGA ACATTTTATT TTTTGTTCAG AAAAAAAAAA CTTTACACAC
     3961 ATAAAATGGC TAGTATGAAT AGCCATATTT TATATAAATT AAATCCTATG AATTTATGAC
     4021 CATATTAAAA ATTTAGATAT TTATGGAACA TAATATGTTT GAAACAATAA GACAAAATTA
     4081 TTATTATTAT TATTATTTTT ACTGTTATAA TTATGTTGTC TCTTCAATGA TTCATAAATA
     4141 GTTGGACTTG ATTTTTAAAA TGTTTATAAT ATGATTAGCA TAGTTAAATA AAAAAAGTTG
     4201 AAAAATTAAA AAAAAACATA TAAACACAAA TGATGTTTTT TCCTTCAATT Tcggcgcctg
     4261 atgcggtatt ttctccttac gcatctgtgc ggtatttcac accgcatatg gtgcactctc
     4321 agtacaatct gctctgatgc cgcatagtta agccagcccc gacacccgcc aacacccgct
     4381 gacgcgccct gacgggcttg tctgctcccg gcatccgctt acagacaagc tgtgaccgtc
     4441 tccgggagct gcatgtgtca gaggttttca ccgtcatcac cgaaacgcgc gagacgaaag
     4501 ggcctcgtga tacgcctatt tttataggtt aatgtcatga taataatggt ttcttagacg
     4561 tcaggtggca cttttcgggg aaatgtgcgc ggaaccccta tttgtttatt tttctaaata
     4621 cattcaaata tgtatccgct catgagacaa taaccctgat aaatgcttca ataatattga
     4681 aaaaggaaga gtatgagtat tcaacatttc cgtgtcgccc ttattccctt ttttgcggca
     4741 ttttgccttc ctgtttttgc tcacccagaa acgctggtga aagtaaaaga tgctgaagat
     4801 cagttgggtg cacgagtggg ttacatcgaa ctggatctca acagcggtaa gatccttgag
     4861 agttttcgcc ccgaagaacg ttttccaatg atgagcactt ttaaagttct gctatgtggc
     4921 gcggtattat cccgtattga cgccgggcaa gagcaactcg gtcgccgcat acactattct
     4981 cagaatgact tggttgagta ctcaccagtc acagaaaagc atcttacgga tggcatgaca
     5041 gtaagagaat tatgcagtgc tgccataacc atgagtgata acactgcggc caacttactt
     5101 ctgacaacga tcggaggacc gaaggagcta accgcttttt tgcacaacat gggggatcat
     5161 gtaactcgcc ttgatcgttg ggaaccggag ctgaatgaag ccataccaaa cgacgagcgt
     5221 gacaccacga tgcctgtagc aatggcaaca acgttgcgca aactattaac tggcgaacta
     5281 cttactctag cttcccggca acaattaata gactggatgg aggcggataa agttgcagga
     5341 ccacttctgc gctcggccct tccggctggc tggtttattg ctgataaatc tggagccggt
     5401 gagcgtgggt ctcgcggtat cattgcagca ctggggccag atggtaagcc ctcccgtatc
     5461 gtagttatct acacgacggg gagtcaggca actatggatg aacgaaatag acagatcgct
     5521 gagataggtg cctcactgat taagcattgg taactgtcag accaagttta ctcatatata
     5581 ctttagattg atttaaaact tcatttttaa tttaaaagga tctaggtgaa gatccttttt
     5641 gataatctca tgaccaaaat cccttaacgt gagttttcgt tccactgagc gtcagacccc
     5701 gtagaaaaga tcaaaggatc ttcttgagat cctttttttc tgcgcgtaat ctgctgcttg
     5761 caaacaaaaa aaccaccgct accagcggtg gtttgtttgc cggatcaaga gctaccaact
     5821 ctttttccga aggtaactgg cttcagcaga gcgcagatac caaatactgt ccttctagtg
     5881 tagccgtagt taggccacca cttcaagaac tctgtagcac cgcctacata cctcgctctg
     5941 ctaatcctgt taccagtggc tgctgccagt ggcgataagt cgtgtcttac cgggttggac
     6001 tcaagacgat agttaccgga taaggcgcag cggtcgggct gaacgggggg ttcgtgcaca
     6061 cagcccagct tggagcgaac gacctacacc gaactgagat acctacagcg tgagcattga
     6121 gaaagcgcca cgcttcccga agggagaaag gcggacaggt atccggtaag cggcagggtc
     6181 ggaacaggag agcgcacgag ggagcttcca gggggaaacg cctggtatct ttatagtcct
     6241 gtcgggtttc gccacctctg acttgagcgt cgatttttgt gatgctcgtc aggggggcgg
     6301 agcctatgga aaaacgccag caacgcggcc tttttacggt tcctggcctt ttgctggcct
     6361 tttgctcaca tgttctttcc tgcgttatcc cctgattctg tggataaccg tattaccgcc
     6421 tttgagtgag ctgataccgc tcgccgcagc cgaacgaccg agcgcagcga gtcagtgagc
     6481 gaggaagcgg aagagcgccc aatacgcaaa ccgcctctcc ccgcgcgttg gccgattcat
     6541 taatgcagct ggcacgacag gtttcccgac tggaaagcgg gcagtgagcg caacgcaatt
     6601 aatgtgagtt agctcactca ttaggcaccc caggctttac actttatgct tccggctcgt
     6661 atgttgtgtg gaattgtgag cggataacaa tttcacacag gaaacagcta tgaccatgat
     6721 tacgccaagc ttgataattc ctgcagccca gcttaattct tttcgagctc tttatgctta
     6781 agtttacaat ttaatattca tactttaagt attttttgta gtatcctaga tattgtgctt
     6841 taaatgctca cccctcaaag caccagtaat attttcatcc actgaaatac cattaaattt
     6901 tcaaaaaaat actatgcata taatgttata catataaaca taaaacgcca tgtaaatcaa
     6961 aaaatatata aaaatatgta taaaaataaa tatgcactaa atataagcta attatgcata
     7021 aaaattaaag tgccctttat taactagcta gtcgtaatta tttatatttc tatgttataa
     7081 aaaaatcctc atataataat ataattaata tatgtaatgt tttttttatt ttataatttt
     7141 aatataaaat aatatgtaaa ttaattcaaa aaataaatat aattgttgtg aaacaaaaaa
     7201 cgtaattttt tcatttgcct tcaaaattta aatttatttt aatatttcct aaaatatata
     7261 tactttgtgt ataaatatat aaaaatatat atttgcttat aaataaataa aaaattttat
     7321 aaaacatagg gggatccatg gttggttcgc taaactgcat cgtcgctgtg tcccagaaca
     7381 tgggcatcgg caagaacggg gacctgccct ggccaccgct caggaacgaa tttagatatt
     7441 tccagagaat gaccacaacc tcttcagtag aaggtaaaca gaatctggtg attatgggta
     7501 agaagacctg gttctccatt cctgagaaga atcgaccttt aaagggtaga attaatttag
     7561 ttctcagcag agaactcaag gaacctccac aaggagctca ttttctttcc agaagtctag
     7621 atgatgcctt aaaacttact gaacaaccag aattagcaaa taaagtagac atggtctgga
     7681 tagttggtgg cagttctgtt tataaggaag ccatgaatca cccaggccat cttaaactat
     7741 ttgtgacaag gatcatgcaa gactttgaaa gtgacacgtt ttttccagaa attgatttgg
     7801 agaaatataa acttctgcca gaatacccag gtgttctctc tgatgtccag gaggagaaag
     7861 gcattaagta caaatttgaa gtatatgaga agaatgatta aggatcccgt ttttcttact
     7921 tatatattta taccaattga ttgtatttat aactgtaaaa atgtgtatgt tgtgtgcata
     7981 tttttttttg tgcatgcaca tgcatgtaaa tagctaaaat tatgaacatt ttattttttg
     8041 ttcagaaaaa aaaaacttta cacacataaa atggctagta tgaatagcca tattttatat
     8101 aaattaaatc ctatgaattt atgaccatat taaaaattta gatatttatg gaacataata
     8161 tgtttgaaac aataagacaa aattattatt attattatta tttttactgt tataattatg
     8221 ttgtctcttc aatgattcat aaatagttgg acttgatttt taaaatgttt ataatatgat
     8281 tagcatagtt aaataaaaaa agttgaaaaa ttaaaaaaaa acatataaac acaaatgatg
     8341 ttttttcctt caatttcgat g
//
>Py1299_PyNOT1-G_DTTPbd
LOCUS       pSL1299                 9313 bp    DNA        circular     21-APR-2021
SOURCE
  ORGANISM
COMMENT     Lindner Lab - used to create Py1299 = Py17XNL PyNOT1-G replaces
            TTP-binding domain with GFPmut2
COMMENT
COMMENT     ApEinfo:methylated:1
FEATURES             Location/Qualifiers
     exon            5165..5728
                     /vntifkey="61"
                     /locus_tag="hDHFR"
                     /label="hDHFR"
                     /ApEinfo_label="hDHFR"
                     /ApEinfo_fwdcolor="pink"
                     /ApEinfo_revcolor="pink"
                     /ApEinfo_graphicformat="arrow_data {{0 1 2 0 0 -1} {} 0}
                     width 5 offset 0"
     CDS             7791..8501
                     /vntifkey="4"
                     /locus_tag="GFPmut2"
                     /label="GFPmut2"
                     /ApEinfo_label="GFPmut2"
                     /ApEinfo_fwdcolor="pink"
                     /ApEinfo_revcolor="pink"
                     /ApEinfo_graphicformat="arrow_data {{0 1 2 0 0 -1} {} 0}
                     width 5 offset 0"
     misc_feature    1631..2078
                     /locus_tag="PbDHFR/TS\3'UTR"
                     /label="PbDHFR/TS\3'UTR"
                     /ApEinfo_label="PbDHFR/TS\3'UTR"
                     /ApEinfo_fwdcolor="#ff0000"
                     /ApEinfo_revcolor="green"
                     /ApEinfo_graphicformat="arrow_data {{0 1 2 0 0 -1} {} 0}
                     width 5 offset 0"
     rep_origin      3475..4157
                     /locus_tag="ColE1 origin"
                     /label="ColE1 origin"
                     /ApEinfo_label="ColE1 origin"
                     /ApEinfo_fwdcolor="gray50"
                     /ApEinfo_revcolor="gray50"
                     /ApEinfo_graphicformat="arrow_data {{0 1 2 0 0 -1} {} 0}
                     width 5 offset 0"
     CDS             2718..3377
                     /locus_tag="AmpR"
                     /label="AmpR"
                     /ApEinfo_label="AmpR"
                     /ApEinfo_fwdcolor="yellow"
                     /ApEinfo_revcolor="yellow"
                     /ApEinfo_graphicformat="arrow_data {{0 1 2 0 0 -1} {} 0}
                     width 5 offset 0"
     misc_feature    join(4561..4870,4875..5159)
                     /locus_tag="PbEF1a 5'UTR"
                     /label="PbEF1a 5'UTR"
                     /ApEinfo_label="PbEF1a 5'UTR"
                     /ApEinfo_fwdcolor="#804040"
                     /ApEinfo_revcolor="#804040"
                     /ApEinfo_graphicformat="arrow_data {{0 1 2 0 0 -1} {} 0}
                     width 5 offset 0"
     misc_feature    5736..6183
                     /locus_tag="PbDHFR-TS 3'UTR"
                     /label="PbDHFR-TS 3'UTR"
                     /ApEinfo_label="PbDHFR-TS 3'UTR"
                     /ApEinfo_fwdcolor="cyan"
                     /ApEinfo_revcolor="green"
                     /ApEinfo_graphicformat="arrow_data {{0 1 2 0 0 -1} {} 0}
                     width 5 offset 0"
     misc_feature    8523..9300
                     /locus_tag="NOT1G 3' Homology Arm (Coding Seq for
                     AA200-462)"
                     /label="NOT1G 3' Homology Arm (Coding Seq for AA200-462)"
                     /ApEinfo_label="NOT1G 3' Homology Arm (Coding Seq for
                     AA200-462)"
                     /ApEinfo_fwdcolor="cyan"
                     /ApEinfo_revcolor="green"
                     /ApEinfo_graphicformat="arrow_data {{0 1 2 0 0 -1} {} 0}
                     width 5 offset 0"
     misc_feature    15..742
                     /locus_tag="PyNOT1-G 5' Homology Arm"
                     /label="PyNOT1-G 5' Homology Arm"
                     /ApEinfo_label="PyNOT1-G 5' Homology Arm"
                     /ApEinfo_fwdcolor="cyan"
                     /ApEinfo_revcolor="green"
                     /ApEinfo_graphicformat="arrow_data {{0 0.5 0 1 2 0 0 -1 0
                     -0.5} {} 0} width 5 offset 0"
     misc_feature    6218..7784
                     /locus_tag="PyNOT1-G Promoter and 5'UTR"
                     /label="PyNOT1-G Promoter and 5'UTR"
                     /ApEinfo_label="PyNOT1-G Promoter and 5'UTR"
                     /ApEinfo_fwdcolor="cyan"
                     /ApEinfo_revcolor="green"
                     /ApEinfo_graphicformat="arrow_data {{0 0.5 0 1 2 0 0 -1 0
                     -0.5} {} 0} width 5 offset 0"
ORIGIN
        1 cctaggcgag atctCCCCCC TATAATAAAA AACCCCATAC ACTTATTTAG ATTTCTACAA
       61 TATATCTTAT CATAACATTT ACTTTTTCTC ATTAACTTTG TTTATCAACT AACATTATTA
      121 TATTTATAAA AATATATTAT CATTATTATT CTTAGTTCTA ATGTATATGC TTTTTAATTT
      181 AATAATATGA GAATTATAAA AAAAACAAAT TCTAGGGGGG AAATTAAAGT GGATTCGCAT
      241 AATCCTTAAA CGAAATATTC TCATAACATA AAACCATGGT AAAAGCCATT GATTATTCTT
      301 ATAAAAATAT TAAGAAATAT ATTAACTTAA TCATTATGTC TGTACATTTA TATATATATA
      361 ACTGCATATA TATGATAAGC ATTTTTTCTA CATAAAAATT GTGGTTTCTC GTTATTGTAT
      421 TTGAGAATAA TATACCCCTT TTATTCCTTT TCATAAAATA TCTTATTACT AAAGTTCAGA
      481 AAAATTATAT ATTTTTTAAT TTTAAAATAA AAGCATAACA AATTATATGT ATAGGCTAAC
      541 GCAAAAAAAA TAATCTTTAA CATATGTATT AAAAAAATAA TAAGACTTAT ATTTATTTTA
      601 AGTGATGAGT ATATATAATC ATAATTATAA TCAATTTTAA CTGTCCCTAT TTAAACAAAT
      661 AAAATCACCG TAGTTGAGTA TAGGATAGAA CATTTGTTCA TTCTCTTACT GCTCACAAAA
      721 AATATAAATA TAGAACAAAT ACacgcgtgg tggcggtctg aacgacatct tcgaggctca
      781 gaaaatcgaa tggcacgaag gcggtggcct caatgatata tttgaagcgc agaagatcga
      841 gtggcatgag ggtggcggtc tgaatgacat attcgaagcg cagaagatcg agtggcacga
      901 gtaacctagt agtaaaggag aagaactttt cactggagtt gtcccAATTC TTGTTGAATT
      961 AGATGGTGAT GTTAATGGGC ACAAATTTTC TGTCAGTGGA GAGGGTGAAG GTGATGCAAC
     1021 ATACGGAAAA CTTACCCTTA AATTTATTTG CACTACTGGA AAACTACCTG TTCCATGGCC
     1081 AACACTTGTC ACTACTTTCG CGTATGGTCT TCAATGCTTT GCGAGATACC CAGATCATAT
     1141 GAAACAGCAT GACTTTTTCA AGAGTGCCAT GCCCGAAGGT TATGTACAGG AAAGAACTAT
     1201 ATTTTTCAAA GATGACGGGA ACTACAAGAC ACGTGCTGAA GTCAAGTTTG AAGGTGATAC
     1261 CCTTGTTAAT AGAATCGAGT TAAAAGGTAT TGATTTTAAA GAAGATGGAA ACATTCTTGG
     1321 ACACAAATTG GAATACAACT ATAACTCACA CAATGTATAC ATCATGGCAG ACAAACAAAA
     1381 GAATGGAATC AAAGTTAACT TCAAAATTAG ACACAACATT GAAGATGGAA GCGTTCAACT
     1441 AGCAGACCAT TATCAACAAA ATACTCCAAT TGGCGATGGC CCTGTCCTTT TACCAGACAA
     1501 CCATTACCTG TCCACACAAT CTGCCCTTTC GAAAGATCCC AACGAAAAGA GAGACCACAT
     1561 GGTCCTTCTT GAGTTTGTAA CAGCTGCTGG GATTACACAT GGCATGGATG AACTATACAA
     1621 ATAAggatcc GTTTTTCTTA CTTATATATT TATACCAATT GATTGTATTT ATAACTGTAA
     1681 AAATGTGTAT GTTGTGTGCA TATTTTTTTT TGTGCATGCA CATGCATGTA AATAGCTAAA
     1741 ATTATGAACA TTTTATTTTT TGTTCAGAAA AAAAAAACTT TACACACATA AAATGGCTAG
     1801 TATGAATAGC CATATTTTAT ATAAATTAAA TCCTATGAAT TTATGACCAT ATTAAAAATT
     1861 TAGATATTTA TGGAACATAA TATGTTTGAA ACAATAAGAC AAAATTATTA TTATTATTAT
     1921 TATTTTTACT GTTATAATTA TGTTGTCTCT TCAATGATTC ATAAATAGTT GGACTTGATT
     1981 TTTAAAATGT TTATAATATG ATTAGCATAG TTAAATAAAA AAAGTTGAAA AATTAAAAAA
     2041 AAACATATAA ACACAAATGA TGTTTTTTCC TTCAATTTcg gcgcctgatg cggtattttc
     2101 tccttacgca tctgtgcggt atttcacacc gcatatggtg cactctcagt acaatctgct
     2161 ctgatgccgc atagttaagc cagccccgac acccgccaac acccgctgac gcgccctgac
     2221 gggcttgtct gctcccggca tccgcttaca gacaagctgt gaccgtctcc gggagctgca
     2281 tgtgtcagag gttttcaccg tcatcaccga aacgcgcgag acgaaagggc ctcgtgatac
     2341 gcctattttt ataggttaat gtcatgataa taatggtttc ttagacgtca ggtggcactt
     2401 ttcggggaaa tgtgcgcgga acccctattt gtttattttt ctaaatacat tcaaatatgt
     2461 atccgctcat gagacaataa ccctgataaa tgcttcaata atattgaaaa aggaagagta
     2521 tgagtattca acatttccgt gtcgccctta ttcccttttt tgcggcattt tgccttcctg
     2581 tttttgctca cccagaaacg ctggtgaaag taaaagatgc tgaagatcag ttgggtgcac
     2641 gagtgggtta catcgaactg gatctcaaca gcggtaagat ccttgagagt tttcgccccg
     2701 aagaacgttt tccaatgatg agcactttta aagttctgct atgtggcgcg gtattatccc
     2761 gtattgacgc cgggcaagag caactcggtc gccgcataca ctattctcag aatgacttgg
     2821 ttgagtactc accagtcaca gaaaagcatc ttacggatgg catgacagta agagaattat
     2881 gcagtgctgc cataaccatg agtgataaca ctgcggccaa cttacttctg acaacgatcg
     2941 gaggaccgaa ggagctaacc gcttttttgc acaacatggg ggatcatgta actcgccttg
     3001 atcgttggga accggagctg aatgaagcca taccaaacga cgagcgtgac accacgatgc
     3061 ctgtagcaat ggcaacaacg ttgcgcaaac tattaactgg cgaactactt actctagctt
     3121 cccggcaaca attaatagac tggatggagg cggataaagt tgcaggacca cttctgcgct
     3181 cggcccttcc ggctggctgg tttattgctg ataaatctgg agccggtgag cgtgggtctc
     3241 gcggtatcat tgcagcactg gggccagatg gtaagccctc ccgtatcgta gttatctaca
     3301 cgacggggag tcaggcaact atggatgaac gaaatagaca gatcgctgag ataggtgcct
     3361 cactgattaa gcattggtaa ctgtcagacc aagtttactc atatatactt tagattgatt
     3421 taaaacttca tttttaattt aaaaggatct aggtgaagat cctttttgat aatctcatga
     3481 ccaaaatccc ttaacgtgag ttttcgttcc actgagcgtc agaccccgta gaaaagatca
     3541 aaggatcttc ttgagatcct ttttttctgc gcgtaatctg ctgcttgcaa acaaaaaaac
     3601 caccgctacc agcggtggtt tgtttgccgg atcaagagct accaactctt tttccgaagg
     3661 taactggctt cagcagagcg cagataccaa atactgtcct tctagtgtag ccgtagttag
     3721 gccaccactt caagaactct gtagcaccgc ctacatacct cgctctgcta atcctgttac
     3781 cagtggctgc tgccagtggc gataagtcgt gtcttaccgg gttggactca agacgatagt
     3841 taccggataa ggcgcagcgg tcgggctgaa cggggggttc gtgcacacag cccagcttgg
     3901 agcgaacgac ctacaccgaa ctgagatacc tacagcgtga gcattgagaa agcgccacgc
     3961 ttcccgaagg gagaaaggcg gacaggtatc cggtaagcgg cagggtcgga acaggagagc
     4021 gcacgaggga gcttccaggg ggaaacgcct ggtatcttta tagtcctgtc gggtttcgcc
     4081 acctctgact tgagcgtcga tttttgtgat gctcgtcagg ggggcggagc ctatggaaaa
     4141 acgccagcaa cgcggccttt ttacggttcc tggccttttg ctggcctttt gctcacatgt
     4201 tctttcctgc gttatcccct gattctgtgg ataaccgtat taccgccttt gagtgagctg
     4261 ataccgctcg ccgcagccga acgaccgagc gcagcgagtc agtgagcgag gaagcggaag
     4321 agcgcccaat acgcaaaccg cctctccccg cgcgttggcc gattcattaa tgcagctggc
     4381 acgacaggtt tcccgactgg aaagcgggca gtgagcgcaa cgcaattaat gtgagttagc
     4441 tcactcatta ggcaccccag gctttacact ttatgcttcc ggctcgtatg ttgtgtggaa
     4501 ttgtgagcgg ataacaattt cacacaggaa acagctatga ccatgattac gccaagcttg
     4561 ataattcctg cagcccagct taattctttt cgagctcttt atgcttaagt ttacaattta
     4621 atattcatac tttaagtatt ttttgtagta tcctagatat tgtgctttaa atgctcaccc
     4681 ctcaaagcac cagtaatatt ttcatccact gaaataccat taaattttca aaaaaatact
     4741 atgcatataa tgttatacat ataaacataa aacgccatgt aaatcaaaaa atatataaaa
     4801 atatgtataa aaataaatat gcactaaata taagctaatt atgcataaaa attaaagtgc
     4861 cctttattaa ctagctagtc gtaattattt atatttctat gttataaaaa aatcctcata
     4921 taataatata attaatatat gtaatgtttt ttttatttta taattttaat ataaaataat
     4981 atgtaaatta attcaaaaaa taaatataat tgttgtgaaa caaaaaacgt aattttttca
     5041 tttgccttca aaatttaaat ttattttaat atttcctaaa atatatatac tttgtgtata
     5101 aatatataaa aatatatatt tgcttataaa taaataaaaa attttataaa acataggggg
     5161 atccatggtt ggttcgctaa actgcatcgt cgctgtgtcc cagaacatgg gcatcggcaa
     5221 gaacggggac ctgccctggc caccgctcag gaacgaattt agatatttcc agagaatgac
     5281 cacaacctct tcagtagaag gtaaacagaa tctggtgatt atgggtaaga agacctggtt
     5341 ctccattcct gagaagaatc gacctttaaa gggtagaatt aatttagttc tcagcagaga
     5401 actcaaggaa cctccacaag gagctcattt tctttccaga agtctagatg atgccttaaa
     5461 acttactgaa caaccagaat tagcaaataa agtagacatg gtctggatag ttggtggcag
     5521 ttctgtttat aaggaagcca tgaatcaccc aggccatctt aaactatttg tgacaaggat
     5581 catgcaagac tttgaaagtg acacgttttt tccagaaatt gatttggaga aatataaact
     5641 tctgccagaa tacccaggtg ttctctctga tgtccaggag gagaaaggca ttaagtacaa
     5701 atttgaagta tatgagaaga atgattaagg atcccgtttt tcttacttat atatttatac
     5761 caattgattg tatttataac tgtaaaaatg tgtatgttgt gtgcatattt ttttttgtgc
     5821 atgcacatgc atgtaaatag ctaaaattat gaacatttta ttttttgttc agaaaaaaaa
     5881 aactttacac acataaaatg gctagtatga atagccatat tttatataaa ttaaatccta
     5941 tgaatttatg accatattaa aaatttagat atttatggaa cataatatgt ttgaaacaat
     6001 aagacaaaat tattattatt attattattt ttactgttat aattatgttg tctcttcaat
     6061 gattcataaa tagttggact tgatttttaa aatgtttata atatgattag catagttaaa
     6121 taaaaaaagt tgaaaaatta aaaaaaaaca tataaacaca aatgatgttt tttccttcaa
     6181 tttcgatggg taccCCGCGG TGgcggccgc TctcgagGAG TATAGGATAG AACATTTGTT
     6241 CATTCTCTTA CTGCTCACAA AAAATATAAA TATAGAACAA ATACATATAT AAAAATATAT
     6301 CTACACACAT ATATATACAT GTATGAACAT AAAACATATG TACACAATAT AAATGAACTT
     6361 GAGTGGAAAA TTAATGTCAT AAGAATATTG TATATGTATT TTGGCAAATA ATATTAATTA
     6421 TTTAAATTGG AGTATAAAAA TCATCCCAAA ATAAACTAAA AATACACACA TTAATAAGAT
     6481 TAGTTAAAAA AATATATATT GTTAAATAAA TACCTAGAAT TAAAAAGGGG TCATATATAT
     6541 CCCTAATATA TACAATATAA AATATAAAAA CGTATGAAGC GAGTTACGAA ATTGTACTAC
     6601 TTTTTTATCC CCTCGTTTCC TGTTAACATT AAATAAACAT GACAAAAAAA AGAAAATAAT
     6661 CATAAAAAAT ATTGCAGAAT TTACACTTTT AATCTTTTTA TATATATTAT TTATTTTGCA
     6721 AATAAAATAA TAAAAAACGC TTTTACCAAA CCTAATATAA CATGAAAATT GTTATGAGAA
     6781 ATAATGTATA AAAAAGGAAA ATACAATTCA AAAATCGTAA ACCTTAAAAA AAGGTATATA
     6841 AATATTCACC ATATTAATAA GCAATTCATA AAATGTAAAA GAACAGTGTA TGCGCAATTA
     6901 TGAACAAATT ATTACCTAGT TTATTTATCT ACAAACGTAT GCCCACAATT TCAACTATAC
     6961 AATGTGTTAT ATGTATAATA AATATTTGTA TACAATATAT ACTTATTAAT ATAATACAAT
     7021 ATATATATAT ATATACATAA TTAATATATT TTTAGTGCAT ATAAATGTAC TTATTTAATT
     7081 CGTAAAAATA TTGCTAATTA AACCCCAATA TTGTATACAT ATACTAAGGA CATAAATTCC
     7141 TCTACGGTTT TATTCACATG CATTAAAAAA ATATAACTTA AGTATATTTT TTCTGTATAT
     7201 CCCCAAAAAA AAAAATGTGG TCAAAAAAAA TATTATATAT ATATAATTAA TTTTATAGGA
     7261 AAAAAACATT CCCAATAAAA TTAAAAAAAA TTCTCAATTT TTTTAATATT ATTTTCCTTT
     7321 TTGGTGATAT TCATATATAG TTAATGAATA TAACATAACA CCAATTTATA ATCCTGCGTT
     7381 TACTTTATAT GTAAAGTTTT CATAAAATCA TAATAACTAC GCAGTTATAA TAAGTTAAAA
     7441 AGGGAAAATG AAAAGAAAAA AAATATATTT TTCTATATGT GTGTGCATAT ATATTGTATA
     7501 TATATATTTA TAATCATAAA CAAAATAAGG CAAAAAAAAA TACCAAAAAA AAATAATAAA
     7561 AAATAAAATA AATAAATAAC CTCCAAAAAA ATAGAAAACA ATCCAAACAA GCCAAACATA
     7621 CCAAACATAC CAAACTTAGC AATCACAGCT GAAATTAAAA AAATAACAAA CACACCAAAT
     7681 CGTGTAAACG CAAAAAATAC CACATAACAA AACACAGTCA AATACCCATA TATATTGTTC
     7741 ATACCATATT GAGGTAATTA CAATAAAACA TGTGCTAAAA AATGactagt agtaaaggag
     7801 aagaactttt cactggagtt gtcccaattc ttgttgaatt agatggtgat gttaatgggc
     7861 acaaattttc tgtcagtgga gagggtgaag gtgatgcaac atacggaaaa cttaccctta
     7921 aatttatttg cactactgga aaactacctg ttccatggcc aacacttgtc actactttcg
     7981 cgtatggtct tcaatgcttt gcgagatacc cagatcatat gaaacagcat gactttttca
     8041 agagtgccat gcccgaaggt tatgtacagg aaagaactat atttttcaaa gatgacggga
     8101 actacaagac acgtgctgaa gtcaagtttg aaggtgatac ccttgttaat agaatcgagt
     8161 taaaaggtat tgattttaaa gaagatggaa acattcttgg acacaaattg gaatacaact
     8221 ataactcaca caatgtatac atcatggcag acaaacaaaa gaatggaatc aaagttaact
     8281 tcaaaattag acacaacatt gaagatggaa gcgttcaact agcagaccat tatcaacaaa
     8341 atactccaat tggcgatggc cctgtccttt taccagacaa ccattacctg tccacacaat
     8401 ctgccctttc gaaagatccc aacgaaaaga gagaccacat ggtccttctt gagtttgtaa
     8461 cagctgctgg gattacacat ggcatggatg aactatacaa agccggcgga agcggcggaa
     8521 gcTACAATAA TATTTCGAAC ACAAATATAT CCCATATACC TAACAATGAA ATTAAAAAAA
     8581 TATCTGTAAC AAATGTAAAT AATTTAAAGA ATATCAACAA TTTAGTTCCA CATTTATCTA
     8641 CCATAGATGT TCCAAATCAA ACTACTACTA ATTTTATTAC CCATAATAAT CTTAGTGATA
     8701 ATAATATCGA AAATAGTGTC AATAATAATA TAATTTCTGA ACCCCACAAT GATAACAATA
     8761 ATGTTAACAC TATTAATAAC TCAAATCGAA CAGCTAGCTA CATAAGTAAT AATAATCAAA
     8821 CTCAATTTAC CAATCTTATG AATAATATTT TGAACCAAAA CAATACTACT ACTACTACTA
     8881 ATAACAATAA CAACCTTAAT AATGATAATA TGCTATTATA TATTAACAAT ATTCTACCAA
     8941 ACCTCTCAAA TATAACAGAT AAAAATACTA TACCATCAAA TATAAATAAT TTACAAGCCA
     9001 TTAATAGAAA TAATCCAATA TATAAAAATA AAAACGCCAA TATTAATTTA AATGCTATAT
     9061 CAGCCATTAA TAATATTAAC AATCTTAATA GTGCATATAA TATTAACTTA TTAGACAATA
     9121 ATCCCTCTTC TAATATATTA TCACCAAACT CAAATAGAGT CTATCAAAAT AAAGGTTTTA
     9181 TTAATACAAA CGAACCAACA CATATTAATA GGAACATAAA TCTAGTAACA AATAATATGA
     9241 CACACAATAA AAATTTGAAT TCTACAAACC TCCTAAATAA TACTAATCAA ATTAAAATAG
     9301 ATATAAATAT GCC
//
```
