## Supplementary material for "The *Plasmodium* NOT1-G Paralogue Acts as an Essential Nexus for Sexual Stage Maturation and Parasite Transmission": Supp File 2 - Complete Flow Cytometry Panels

### Hart *et al.* Supplemental File 2 - Biological Replicate 1

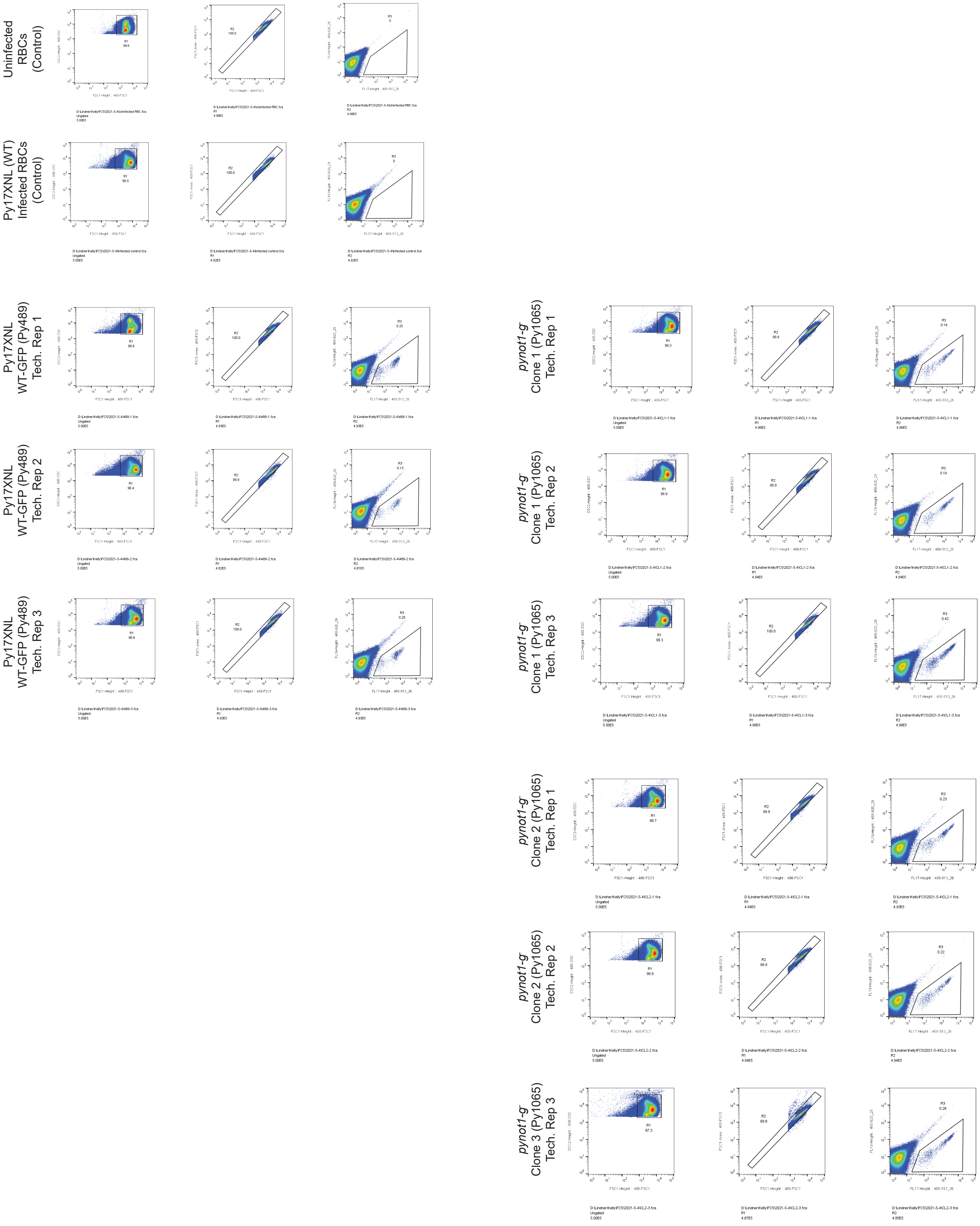

### Hart *et al.* Supplemental File 2 - Biological Replicate 2

Uninfected RBCs (Control)

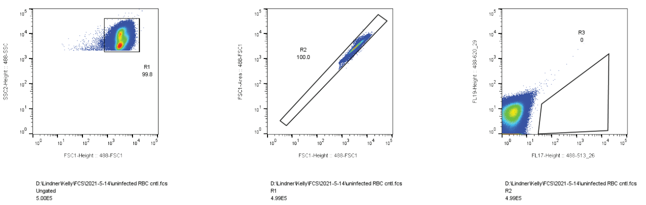

Py17XNL (WT) Infected RBCs (Control)

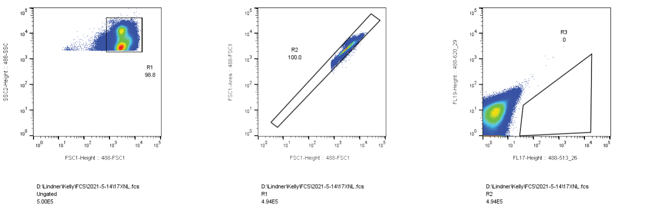

Py17XNL WT-GFP (Py489) Tech. Rep 1

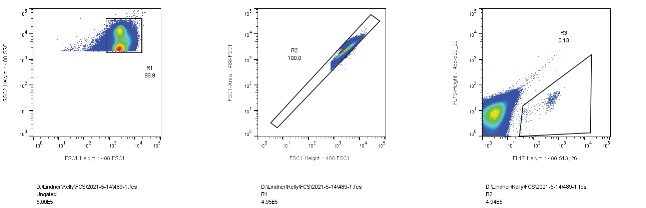

Py17XNL WT-GFP (Py489) Tech. Rep 2

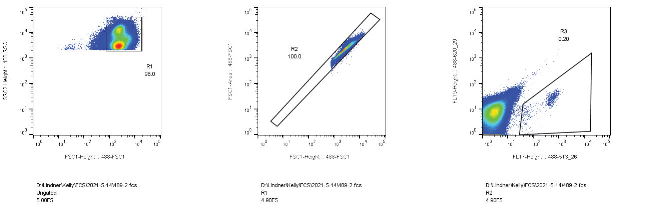

Py17XNL WT-GFP (Py489) Tech. Rep 3

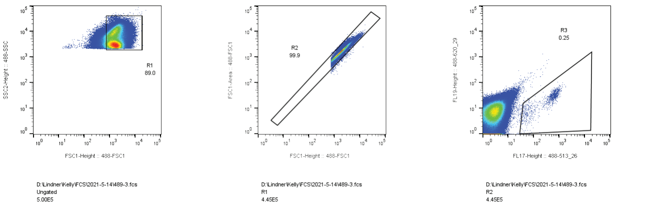

*pynot1-g* Clone 1 (Py1065) Tech. Rep 1

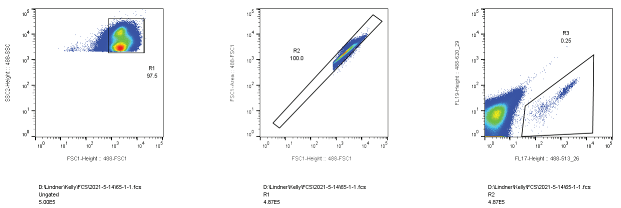

*pynot1-g* Clone 1 (Py1065) Tech. Rep 2

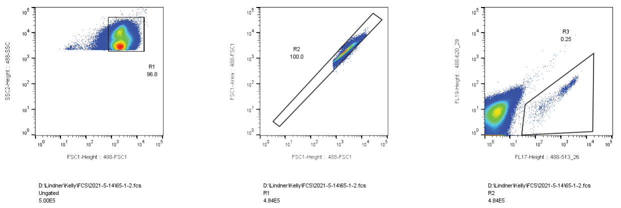

*pynot1-g* Clone 1 (Py1065) Tech. Rep 3

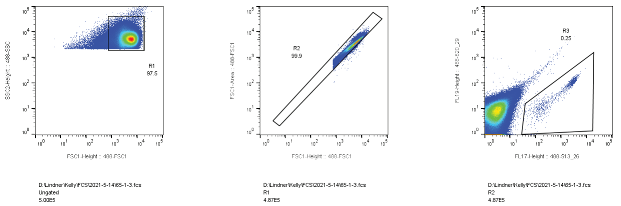

*pynot1-g* Clone 2 (Py1065) Tech. Rep 1

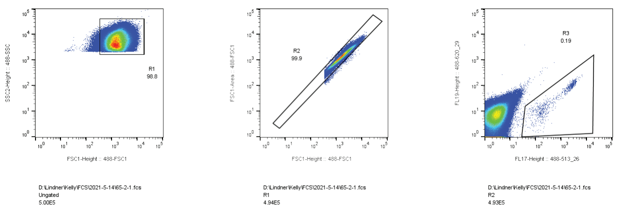

*pynot1-g* Clone 2 (Py1065) Tech. Rep 2

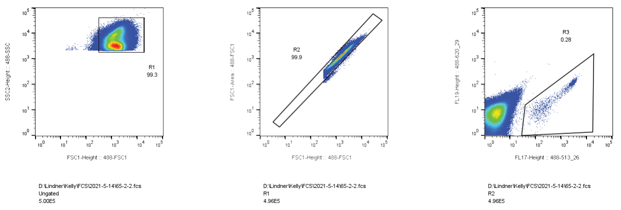

*pynot1-g* Clone 3 (Py1065) Tech. Rep 3

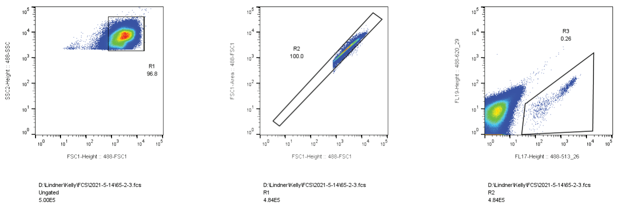
