## Supplementary material for "The *Plasmodium* NOT1-G Paralogue Acts as an Essential Nexus for Sexual Stage Maturation and Parasite Transmission": Supp File 3 - RNA-seq bioinformatics scripts and MakeFile: heatmap.r

```
#!/usr/bin/env Rscript
#
# Draws a heatmap from the output that contains a normalized matrix
#

# Set the plot dimensions.
WIDTH = 12
HEIGHT = 14

# Command line argument.
args = commandArgs(trailingOnly=TRUE)  

# Optional argument for FDR cutoff.
if (length(args) > 0) {
  LIMIT = as.integer(args[1])
} else {
  # Default FDR cutoff
  LIMIT = 5
}

# Input values should be in percent!
LIMIT = LIMIT/100

# Set the margins
MARGINS = c(17, 12)

# Relative heights of the rows in the plot.
LHEI = c(1, 5)

# Load the library.
suppressPackageStartupMessages(library(gplots))

# Read normalized counts from the standard input.
data = read.csv("stdin", header=T, as.is=TRUE)

# Subset data for values under a treshold.
data = subset(data, data$FDR <= LIMIT)

# The heatmap row names will be taken from the first column.
row_names = data[, 1]

# The normalized data starts past the rightmost of these columns.
idx = which(colnames(data) == "falsePos") + 1

# The normalized counts are on the right size.
counts = data[, idx : ncol(data)]

# Load the data from the second column on.
values = as.matrix(counts)

# Adds a little noise to each element to avoid the
# clustering function failure on zero variance rows.
values = jitter(values, factor = 1, amount = 0.00001)

# Normalize each row to a z-score
zscores = NULL
for (i in 1 : nrow(values)) {
    row = values[i,]
    zrow = (row - mean(row)) / sd(row)
    zscores = rbind(zscores, zrow)
}

# Set the row names on the zscores.
row.names(zscores) = row_names

# Turn the data into a matrix for heatmap2.
zscores = as.matrix(zscores)

# Open the drawing device.
pdf('|cat', width = WIDTH, height = HEIGHT)

# Set the color palette.
col = greenred

# Draw the heatmap.
heatmap.2(zscores, col=col, density.info="none", Colv=NULL,
    dendrogram="row", trace="none", margins=MARGINS, lhei=LHEI)

# Turn off the device.
dev.off()
```
