## Supplementary material for "The *Plasmodium* NOT1-G Paralogue Acts as an Essential Nexus for Sexual Stage Maturation and Parasite Transmission": Supp File 3 - RNA-seq bioinformatics scripts and MakeFile: make_gene_tin.py

```
import argparse, csv, sys
from collections import defaultdict

"""
Computing gene level TINs from the transcript TINS.
Uses either average(transcript_tins) or max(transcript_tins) to compute
the gene-level TIN values.
"""


def make_id_dict(fname):
    """
    Reads a file with geneids and transcript_ids
    and stores the transcript ids for each gene in a dict.
    """
    stream = csv.reader(open(fname), delimiter="\t")
    gene_store, transcript_store = defaultdict(list), dict()

    for row in stream:
        gid, tid = row[0], row[1]

        gene_store[gid].append(tid)
        transcript_store[tid] = gid
    return gene_store, transcript_store


def make_gene_TIN(tinfile, mapfile, average, maximum):
    gids, tids = make_id_dict(mapfile)

    transcript_tins = defaultdict(list)

    # Read the transcript tin file and compute gene TIN
    stream = csv.reader(open(tinfile), delimiter="\t")

    # Skip header
    header = next(stream)
    sample = header[3]

    # Collect the transcript tins for each gene.
    for row in stream:
        tid, eff_len, bg, tin = row[:4]
        gid = tids[tid] if tid in tids else None

        if gid:
            transcript_tins[gid].append(float(tin))

    # Print output header

    header = "\t".join(["gene", sample, "remark"])
    print(header)

    # Compute gene level tins.
    for gene, vals in transcript_tins.items():
        gtin = max(vals) if maximum else round(sum(vals) / len(vals), 2)

        remark = ""

        # Create a remark column
        if (len(vals) > 1) and maximum:
            remark = f"Maximum of {len(vals)} transcript TINS"

        if (len(vals) > 1) and average:
            remark = f"Average of {len(vals)} transcript TINS"

        out = "\t".join([gene, str(gtin), remark])
        print(out)


if __name__ == "__main__":
    parser = argparse.ArgumentParser(description='''Makes a gene_level TIN from the transcript TIN values.''')

    parser.add_argument('--tin_file', dest='tin_file', type=str, required=True,
                        help='Tab delimited file with TIN values each transcript.')

    parser.add_argument('--idmap', dest='idmap', type=str, required=True,
                        help='Tab delimited file geneid in the first column and transcript_id in the second column.')

    parser.add_argument('--max', dest='max', required=False, action='store_true',
                        help="""If Specified makes the gene level TIN by taking the maximum of transcript TINs.""")

    parser.add_argument('--avg', dest='avg', required=False, action='store_true',
                        help="""If Specified makes the gene level TIN by taking the avergae of transcript TINs.
                        This is done bu default""")

    args = parser.parse_args()

    tin_file = args.tin_file
    map_file = args.idmap
    maximum = args.max
    avg = args.avg

    if maximum and avg:
        print("""Either --max or --avg should be specified. Not both.
                 --avg is the default.
            """)
        sys.exit()

    if not (maximum and avg):
        avg = True

    if maximum:
        avg = False

    make_gene_TIN(tin_file, map_file, average=avg, maximum=maximum)
```
