## Supplementary material for "The *Plasmodium* NOT1-G Paralogue Acts as an Essential Nexus for Sexual Stage Maturation and Parasite Transmission": Supp File 3 - RNA-seq bioinformatics scripts and MakeFile: Makefile

```
SAMPLES=sampleinfo.txt
DATA=data
FASTQC=$(DATA)/fastqc_raw

REFS=refs
GENOME=$(REFS)/PlasmoDB-50_Pyoeliiyoelii17X_Genome.fasta
TRANSCRIPTS=PlasmoDB-50_Pyoeliiyoelii17X_transcripts.fa
KALLISTO_IDX=$(TRANSCRIPTS).idx

GFF=$(REFS)/PlasmoDB-50_Pyoeliiyoelii17X.gff 
GTF=$(REFS)/PlasmoDB-50_Pyoeliiyoelii17X.gtf

HISAT=hisat2
GCOUNTS=$(HISAT)/featurecounts
KCOUNTS=kallisto
RUNIN=$(HISAT)/runin_overlaps

SCRIPTS=scripts
IDMAP=$(REFS)/idmap.txt

rename_files:
	#
	# Include sample info in the filename for better readability.
	#
	cat $(SAMPLES) | grep -v "sample" | parallel --col-sep "\t" mv $(DATA)/{1}*R1*fastq.gz $(DATA)/"{2}.fastq.gz"

samples:
	#
	# Get sample names
	#
	cat $(SAMPLES) | parallel --col-sep "\t" echo {2} >samples.txt

fastqc:
	#
	# Create fastqc reports
	#
	mkdir -p $(FASTQC)
	cat samples.txt | parallel --verbose --progress -j 6 fastqc --nogroup -o $(FASTQC) $(DATA)/{}.fastq.gz

hisat2:
	mkdir -p $(HISAT)
	cat samples.txt | parallel --verbose --progress -j 12 'hisat2 -q -x $(GENOME) --rg-id {} --rg {} --rna-strandness R -U $(DATA)/{}.fastq.gz -S $(HISAT)/{}.sam --max-intronlen 5000 2>$(HISAT)/{}_summary.txt'

sam2bam:
	#       
	#convert sam to bam     
	#
	cat samples.txt |parallel -j 8 --verbose --progress 'samtools view -b $(HISAT)/{}.sam | samtools sort - -o $(HISAT)/{.}.bam -O bam -T $(HISAT)/{}_tmp'
	ls $(HISAT)/*bam | parallel -j 8 --verbose --progress samtools index {}

coverage:
	ls $(HISAT)/*.bam | parallel -j 6 --progress --verbose 'bedtools genomecov -ibam  {}  -split -bg  > {.}.bedgraph'
	ls $(HISAT)/*.bedgraph |parallel -j 8 --progress --verbose bedSort {} {.}.s.bedgraph
	ls $(HISAT)/*.s.bedgraph |sed 's/.s.bedgraph//' | parallel -j 3 --progress --verbose 'bedGraphToBigWig {}.s.bedgraph  $(GENOME).fai {}.bw'

gtf:
	gffread -T -o $(GTF)  $(GFF)

featurecounts:
	#
	# Count reads at gene level.
	#
	mkdir -p $(GCOUNTS)
	featureCounts -T 8 -t exon -s 2 -g gene_id -a $(GTF) -o $(GCOUNTS)/gcounts.txt $(HISAT)/*_asexual.bam $(HISAT)/*_sexual.bam
	#
	# format file.
	cat $(GCOUNTS)/gcounts.txt | grep -v "#" | sed -e '1s/.bam//g' -e '1s/hisat2\///g' >$(GCOUNTS)/gene_counts.txt 
 
transcriptome:
	#
	# Get all transcript sequences from annotation 	
	#
	gffread -W -w $(REFS)/$(TRANSCRIPTS) -g $(GENOME) $(GTF)

kallisto_index:
	kallisto index -i $(REFS)/$(TRANSCRIPT_IDX) $(REFS)/$(TRANSCRIPTS)

kallisto:
	#
	# Count transcript abundance in sense strand. Read1 is reverse, ie, rf-stranded.
	#
	mkdir -p $(KCOUNTS)
	cat samples.txt | parallel --verbose --progress -j 6 kallisto quant -t 4 -i $(REFS)/$(TRANSCRIPT_IDX) -o $(KCOUNTS)/{} --pseudobam --rf-stranded --single -l 150 -s 20 -b 10 $(DATA)/{}.fastq.gz

sort_rename_pseudobam:
	# sort and rename pseudobam
	cat samples.txt | parallel --verbose --progress -j 6  samtools sort  $(KCOUNTS)/{}/pseudoalignments.bam -o $(KCOUNTS)/{}/{}_pseudoalignments.bam
	# index pseudobam
	cat samples.txt | parallel --verbose --progress -j 6 samtools index $(KCOUNTS)/{}/{}_pseudoalignments.bam
	# Remove unsorted  file.
	cat samples.txt | parallel --verbose --progress -j 6 rm -f $(KCOUNTS)/{}/pseudoalignments.bam

depth:
	#
	# Get the depth at each position of transcript from the pseudobam file.
	#
	cat samples.txt | parallel --verbose --progress -j 10 'samtools depth -aa $(KCOUNTS)/{}/{}_pseudoalignments.bam >$(KCOUNTS)/{}/{}_depth.txt'

transcript_integriety:
	#
	# Get coverage uniformity across genome.
	#
	cat samples.txt | parallel --verbose --progress -j 10 'python $(SCRIPTS)/transcript_tin.py --cov $(KCOUNTS)/{}/{}_depth.txt >$(KCOUNTS)/{}/{}_tin.txt'
	#
	# Combine the tin values for all samples into a single table.
	#
	find $(KCOUNTS) -name "*_tin.txt" |sort  >tin_files.txt
	python $(SCRIPTS)/merge.py --file_list tin_files.txt --outfile $(KCOUNTS)/tin_combined.csv --merge_how intersection --merge_col tid,eff-length,background 

get_TPMS:
	cat samples.txt |combine_tpms $(KCOUNTS) |csvcut -t -C X1,X2,X3 | tr "," "\t" >$(KCOUNTS)/tpms.txt

get_counts:
	cat samples.txt |combine_counts $(KCOUNTS) |csvcut -t -C X1,X2,X3 | tr "," "\t" >$(KCOUNTS)/kallisto_counts.txt

gene_tins:$(IDMAP)
	#
	# Compute gene level tins from transcript_tins
	#
	cat samples.txt| parallel --verbose --progress -j 6 'python $(SCRIPTS)/make_gene_tin.py --tin_file $(KCOUNTS)/{}/{}_tin.txt  --idmap $(IDMAP) --max >$(KCOUNTS)/{}/{}_gene_tin.txt'
	#
	# Combine gene tin values for all samples.
	find $(KCOUNTS) -name "*_gene_tin.txt"  >tin_files.txt  
	python $(SCRIPTS)/merge.py --file_list tin_files.txt --outfile $(KCOUNTS)/gene_tin_combined.csv --merge_how intersection --merge_col gene

runin_overlaps:
	#
	# Check runins
	mkdir -p $(RUNIN)
	cat samples.txt |parallel --verbose --progress -j 4 'python $(SCRIPTS)/overlap.py $(HISAT)/{}.bam --ann $(GTF) --strand reverse --count_cutoff 20 --n 0 >$(RUNIN)/{.}_runin_20_40.txt'
```
