## Supplementary material for "The *Plasmodium* NOT1-G Paralogue Acts as an Essential Nexus for Sexual Stage Maturation and Parasite Transmission": Supp File 3 - RNA-seq bioinformatics scripts and MakeFile: merge.py

```
import pandas as pd
from functools import reduce
import glob, sys, argparse


def main():
    parser = argparse.ArgumentParser(description='''Merges tables''')

    parser.add_argument('--file_list', dest='file_list', type=str, required=True,
                        help='File listing all the tables to merge.')
    parser.add_argument('--outfile', dest='outfile', type=str, required=True,
                        help='Output file name')
    parser.add_argument('--merge_how', dest='merge_how', type=str, required=True,
                        help='''Specify how the tables should be merged. It should be either union, intersection or merge.\
                                if merge is specified a left join on the first table is performed.''' )
    parser.add_argument('--merge_col', dest='merge_col', type=str, required=True,
                        help=''' Specify the ID column to be used for merged.''')

    args = parser.parse_args()
    flist = args.file_list
    outfile = args.outfile
    merge_how = args.merge_how # merge, union or intersection
    m_column = args.merge_col #'giraffe_id'
    
    merge_column = m_column.strip().split(",")
    header = m_column.strip().split(",")
   
    merge_type = ""

    if merge_how == "merge":
        merge_type = 'left'  # merge on the 1st table.
    elif merge_how == "intersection":
        merge_type = "inner"
    else:
        merge_type = "outer"
    
    frames = list()

    fh = open(flist)
    for f in fh:
       header.append(f.strip().replace("_tin.txt", ""))
       #header.append('remark') #uncomment for gene tins
       df = pd.read_csv(f.strip(),sep="\t")
       frames.append(df)

    result = reduce(lambda left, right: pd.merge(left, right, on=merge_column,
                                                 how=merge_type), frames)
    
    result.columns = header
    result.to_csv(outfile, index=False)

    return


if __name__ == "__main__":
    main()
 
```
