## Supplementary material for "The *Plasmodium* NOT1-G Paralogue Acts as an Essential Nexus for Sexual Stage Maturation and Parasite Transmission": Supp File 3 - RNA-seq bioinformatics scripts and MakeFile: overlap.py

```
import csv, sys, os, math, random
import argparse, subprocess, string

"""
This script produces raw read-count counts, expected-tins and observed-tins at the gene level.
Inputs are bam files and gff3 file.

Dependencies:
    - featureCounts (conda install -c bioconda subread)
    - bedtools      (conda install -c bioconda bedtools)
    - samtools      (conda install -c bioconda samtools)
    
How it works?

Input(s) : bam file and annotation file
1. Intergenic and intronic annotation files (if background is specified) are created from the input ann file.
2. A bed file merging overlapping features are created (used to get depth per gene while calculating tin)
3. For each gene and intergenic region, counts are calculated in the sense and antisense strand.
4. Depending on gene strand, sense or antisense counts and tins are obtained and compared to that of the gene to
    determine the overlap.

"""


def get_filename(x):
    return os.path.splitext(os.path.basename(x))[0]


def add_string(elms, string):
    def attach(x):
        return x + string

    elms = list(map(attach, elms))
    return elms


def combine_counts(sense_counts, antisense_counts):
    """
    Sum up sense and antisense counts.
    Inputs are dictionary of list of dictionaries.
    Returns a dictionary of list of dictionaries.
    """
    total = dict()
    for key, vals in sense_counts.items():
        # print("***",key, vals)
        if key == "samples":
            total[key] = vals
            continue
        counts = []

        for idx, c in enumerate(vals):
            antisense_vals = antisense_counts[key]

            sample_sense_count = c
            sample_antisense_count = antisense_vals[idx]

            sample_count = sample_sense_count + sample_antisense_count

            counts.append(sample_count)

        total[key] = counts

    return total


def store_counts(fname):
    """
    Store featureCounts output in a dictionary.
    """
    store = dict()
    stream = csv.reader(open(fname, 'rt'), delimiter="\t")

    # skip header rows.
    next(stream)

    header = next(stream)
    samples = header[6:]
    samples = map(get_filename, samples)
    samples = add_string(samples, "_count")
    store.setdefault('samples', []).extend(samples)

    for row in stream:
        # Skip first 5 columns as counts start at 6th column
        gid, glen, cols = row[0], row[5], row[6:]
        for count in cols:
            store.setdefault(gid, []).append(int(count))

    return store


def get_attr_tag(feat):
    tag = {'gene': 'gene_id', 'transcript': 'transcript_id'}
    return tag.get(feat, "ID")


def get_counts(bam, ann, strand="", feat="exon", groupby="gene", paired=False, flag="", log="runlog.txt", ):
    """
    Use featurecounts to get the raw counts.
    """

    countsfile = f"counts_{flag}.txt" if len(bam.split(' ')) > 1 else get_filename(bam) + "_counts.txt"
    countsfile = countsfile.replace("_counts.txt", f'_{flag}_counts.txt') if flag else countsfile
    countsfile = os.path.join(TMP, countsfile)

    # Set paired end or single end.
    pflag = '-p' if paired else ""

    # Get the attr type to use for grouping.
    group_attr = get_attr_tag(groupby)

    # Set strandedness.
    if strand == "both":
        sflag = ""
    else:
        sflag = '-s 2' if strand == "reverse" else '-s 1'

    # Make the command.
    cmd = f'featureCounts --primary -M {pflag} {sflag} -t {feat} -O -g {group_attr} -a {ann} -o {countsfile} {bam} 2>>{log}'
    # print(cmd)

    # Run command and get counts.
    out = subprocess.run(cmd, shell=True, stdout=subprocess.PIPE, stderr=subprocess.PIPE, check=True,
                         universal_newlines=True)

    # Exit on error.
    exit_on_error(out.stderr) if out.stderr and out.returncode != 0 else None

    # Store counts in a dictionary.
    counts = store_counts(countsfile)

    return counts


def get_intron_coords(store):
    """
    Collect all intron coordinates.
    Input store is a dictionary of list where each element in the list is (start,end) of exons/cds
    Returns a dictionary of list where each element in the list is (start,end) of intron.
    """
    introns = dict()
    for key, vals in store.items():

        if len(vals) == 1:  # single exon genes
            continue
        # Collect intron starts and ends.
        istarts = [end + 1 for (start, end) in vals[:-1]]
        iends = [start - 1 for (start, end) in vals[1:]]
        coords = list(zip(istarts, iends))
        introns.setdefault(key, []).extend(coords)
    return introns


def parse(item):
    return item['chrom'], str(item['start']), str(item['end']), item['strand'], item['gid'], item['gtype']


def make_intron_gtf(genes, introns):
    """
    Create a GTF file for introns.
    """
    # intron gtf file.
    intron_ann = os.path.join(TMP, "introns.gtf")
    fi = open(intron_ann, "w")

    for gene, coords in introns.items():
        # Get gene coords
        gene_vals = genes[gene]
        chrom, start, end, strand, gid, gtype = parse(gene_vals)
        # Write gene line.
        gene_attr = f'ID {gid};gene_id {gid}'
        parent_ln = make_ann_row(chrom, gtype, start, end, strand, gene_attr)
        fi.write(parent_ln + "\n")

        for idx, coord in enumerate(coords):
            # Write intron line.
            istart, iend = str(coord[0]), str(coord[1])
            intron_id = f"{gid}-I" + str(idx + 1)
            intron_attr = f'ID {intron_id};gene_id={gid}; transcript_id={gid}'
            intron_ln = make_ann_row(chrom, "intron", istart, iend, strand, intron_attr)
            fi.write(intron_ln + "\n")

    return intron_ann


def get_background_noise(bam, strand, intron_ann, intron_len, groupby, paired):
    """
    Calculate background noise from intron specific gff file.
    Returns a dictionary with background noise.
    """
    noise_file = get_filename(bam) + "_noise.txt"
    noise_file = os.path.join(TMP, noise_file)
    nout = open(noise_file, "w")

    intron_counts = get_counts(bam=bam, ann=intron_ann, strand=strand,
                               feat="intron", groupby=groupby, paired=paired, flag="intron")

    for key, vals in intron_counts.items():
        if key == "samples":
            continue
        for val in vals:
            c, l = val, intron_len.get(key, 0)
            noise = 0 if c == 0 or l == 0 else round(int(c) / int(l), 4)
            nout.write(f'{key}\t{str(noise)}')
            nout.write("\n")

    return noise_file


def extract_attr(attr_str, attr_tag):
    attrs = attr_str.split(";")
    attrs = [a.strip() for a in attrs]  # remove all spaces
    try:
        feat_id = [val for val in attrs if val.startswith(attr_tag)][0]
    except:
        print(f'{attr_tag} not found for the groupby attribute in annotation file')
        sys.exit()
    feat_id = feat_id.replace(attr_tag + "=", "")  # for gff
    feat_id = feat_id.replace(attr_tag + " ", "")  # for gtf
    feat_id = feat_id.replace('"', "")  # remove all quotes.
    return feat_id


def make_genedict(chrom, start, end, strand, gid, gtype):
    g = dict()
    g['chrom'] = chrom
    g['start'] = start
    g['end'] = end
    g['strand'] = strand
    g['gid'] = gid  # feature id
    g['gtype'] = gtype
    return g


def parse_row(row):
    # chrom, start, end, strand, feat
    return row[0], int(row[3]), int(row[4]), row[6], row[2], row[8]


def merge(elms):
    overlapped = list()
    elms = sorted(elms)
    elms.append((1000000000, 1000000000))

    start = end = 0

    for (x1, y1), (x2, y2) in zip(elms, elms[1:]):

        start = x1 if start == 0 else start
        end = y1 if end == 0 else end

        # Case1 : left overlap
        if x2 <= end and y2 > end:
            end = y2

        # Case2 : complete overlap
        if x2 <= end and y2 <= end:
            start = start
            end = end

        # Case3 : no overlap
        if x2 > end:
            # print(start, end)
            overlapped.append((start, end))
            start = end = 0
    return overlapped


def merge_features(feats):
    """
    Make a union of features for each gene by merging overlapping features.
    """
    merged = dict()
    for gene, elms in feats.items():
        combined = merge(elms)
        merged.setdefault(gene, combined)
    return merged


def collect_features(ann, feat_type="exon"):
    """
    Collect the specified features grouped by gene.
    """

    feat_store, genes = dict(), dict()

    # Get groupby attr.
    groupby_attr = ["gene", "pseudogene"]

    # Exons are collected if feat_type is either of transcript, mRNA or intron.
    feat_type = "exon" if feat_type in ["transcript", "mRNA"] else feat_type

    # Get attribute tag for the.
    groupby_tag = "gene_id"

    # Make a list of features to be considered from annotation file.
    features = groupby_attr + [feat_type]

    # Parse file and collect features.
    stream = csv.reader(open(ann), delimiter="\t")

    for row in stream:
        # Continue if header.
        if row[0].startswith("#"):
            continue

        chrom, start, end, strand, feat, attrs = parse_row(row)

        if feat not in features:
            continue

        # Make sure start < end
        if start > end:
            start, end = end, start

        # Store all feature starts and ends for each grouby_attr.
        if feat in groupby_attr:
            gid = extract_attr(attrs, groupby_tag)

            # Store gene coordinates and geneid.
            genes[gid] = make_genedict(chrom=chrom, start=start, end=end, strand=strand, gid=gid, gtype=feat)
            feat_store.setdefault(gid, [])

        if feat == feat_type:
            # Get groupby tag. Doing this again in-case the annotation file is not properly sorted.
            parent = extract_attr(attrs, groupby_tag)

            if parent and parent not in feat_store:
                feat_store.setdefault(parent, [])

            feat_store[parent].append((start, end))

    # Exit if feat_type or group_by did not match with 3rd column in gff file.
    empty_feat_store = [k for k, a in feat_store.items() if a == []]

    if not genes or len(empty_feat_store) == len(genes):
        print(f'{feat_type} do not match with 3rd column in annotation file or \'gene\' feature row '
              f'not found in annotation file. Exiting.')
        sys.exit()
    return genes, feat_store


def check_short_gene(gene1, gene2):
    """
    Checks if gene2 is completely inside gene1.
    """
    chrom1, start1, end1 = gene1['chrom'], gene1['start'], gene1['end']
    chrom2, start2, end2 = gene2['chrom'], gene2['start'], gene2['end']

    if chrom1 == chrom2 and start2 >= start1 and end2 <= end1:
        return True
    return False


def get_neighbors(genes, short):
    """
    Add left and right gene information to the dictionary.
    If a gene is completely inside another, left and right gene information is left empty.
    """

    left_gene, right_gene, = "", ""
    left_gene_strand, right_gene_strand = "", ""

    # Filter short genes
    filtered = {k: v for k, v in genes.items() if k not in short}

    vals = list(filtered.values())
    # Sort genes based on chrom and start
    vals.sort(key=lambda x: (x['chrom'], x['start']))

    for idx, gene in enumerate(vals):

        gid, chrom = gene['gid'], gene['chrom']

        # Not the very first or the very last genes.
        if idx != 0 and idx != len(vals) - 1:
            prev_gene = vals[idx - 1]
            next_gene = vals[idx + 1]
            prev_chrom, next_chrom = prev_gene['chrom'], next_gene['chrom']

            if chrom != next_chrom:
                right_gene, right_gene_strand = "", ""
                left_gene, left_gene_strand = prev_gene['gid'], prev_gene['strand']

            elif chrom != prev_chrom:
                left_gene, left_gene_strand = "", ""
                right_gene, right_gene_strand = next_gene['gid'], next_gene['strand']

            else:
                left_gene = prev_gene['gid']
                left_gene_strand = prev_gene['strand']
                right_gene = next_gene['gid']
                right_gene_strand = next_gene['strand']

        # first gene
        if idx == 0:
            next_gene = vals[idx + 1]
            left_gene, left_gene_strand = "", ""
            right_gene, right_gene_strand = next_gene['gid'], next_gene['strand']

        # last gene
        if idx == len(vals) - 1:
            right_gene, right_gene_strand = "", ""
            prev_gene = vals[idx - 1]
            left_gene, left_gene_strand = prev_gene['gid'], prev_gene['strand']

        gene['left_gene'] = left_gene
        gene['left_gene_strand'] = left_gene_strand
        gene['right_gene'] = right_gene
        gene['right_gene_strand'] = right_gene_strand
        genes[gid] = gene

    # Take care of short genes
    for k, v in genes.items():
        if k in short:
            v['left_gene'], v['right_gene'] = "", ""
            v['left_gene_strand'], v['right_gene_strand'] = "", ""
        genes[k] = v

    return genes


def get_ref_names(string):
    """
    Returns the reference sequences names and length from the sam header string.
    """
    if string.startswith("@SQ"):
        elms = string.split("\t")
        elms = [e.strip() for e in elms]
        name = [e for e in elms if e.startswith("SN:")][0]
        slen = [e for e in elms if e.startswith("LN:")][0]
        name = name.replace("SN:", "")
        slen = slen.replace("LN:", "")

        return name, int(slen)


def parse_bam_header(bam):
    """
    Parses bam header to get reference names and lengths.
    """
    # Get one bam file if there are more than one.
    vals = bam.split(' ')
    aln = vals[0] if len(vals) > 1 else bam

    # Extract header from bam file.
    cmd = f'samtools view -H {aln}'
    res = subprocess.run(cmd, shell=True, stdout=subprocess.PIPE, check=True, universal_newlines=True)
    # Parse header to extract sequence name and length.
    res_list = res.stdout.split("\n")
    refs = map(get_ref_names, res_list)
    refs = filter(None, refs)
    return refs


def bam_to_genome(bam):
    """
    Generate a genome file from bam header.
    The genome file has the format <chrom><tab>size>
    """
    refs = parse_bam_header(bam)
    outfile = os.path.join(TMP, "chrom.txt")
    fh = open(outfile, "w")
    for r in refs:
        out = "\t".join([r[0], str(r[1])])
        fh.write(out + "\n")

    return outfile


def make_ann_row(chrom, feat, start, end, strand, attr):
    return "\t".join([chrom, ".", feat, str(start), str(end), ".", strand, ".", attr])


def make_gtf_feature_rows(chrom, start, end, strand, gene_id, idx=0, feat="exon"):
    """
    Creates gtf feature rows from chr,start,end,strand,gene_id
    """
    # Make ids for gene and feature.
    num = str(idx + 1)
    trans_id = gene_id + "_t" + num
    feat_id = gene_id + "_" + feat + num

    # Make 9th column
    attr = f'ID \"{feat_id}\";gene_id \"{gene_id}\";transcript_id \"{trans_id}\"'

    # Create feature rows.
    feat_ln = make_ann_row(chrom, feat, start, end, strand, attr)
    return feat_ln


def make_bed(coords, outfile="intergenic.bed"):
    """
    Create a bed file from a dictionary with name as key and coords as vals
    """
    outfile = os.path.join(TMP, outfile)
    fout = open(outfile, "w")

    for name, vals in coords.items():
        chrom, start, end = vals[0], vals[1], vals[2]
        out = "\t".join([chrom, str(start), str(end), name, ".", "+"])
        write_to_file([out], fout)
    return outfile


def make_intergenic_files(coords):
    ig_ann = make_intergenic_gtf(coords)
    ig_bed = make_bed(coords)
    return ig_bed, ig_ann


def write_to_file(rows, fh):
    for row in rows:
        fh.write(row + "\n")


def make_intergenic_gtf(coords, outfile="intergenic.gtf"):
    """
    Create a minimal gtf file for intergenic regions.
    """

    outfile = os.path.join(TMP, outfile)
    fout = open(outfile, "w")

    idx = 0
    for key, vals in coords.items():
        gid = key
        chrom, start, end = vals[0], vals[1], vals[2]
        feat_ln = make_gtf_feature_rows(chrom, start, end, "+", gid, idx, "intergenic")
        write_to_file([feat_ln], fout)
    return outfile


def name_intergenic_regions(intergenic, genes, short):
    """"
    Name the intergenic intervals with the geneids of the neighboring genes in the format gene_id1:geneid2
    if (igstart, igend) are in between gene1-start and gene2-start a, then it will get the name geen1:gene2.
    Returns a dictionary with intergenic-name as key and its coordinates as values.
    """
    ignames = dict()

    gcoords, igcoords, gnames = [], [], []

    # Filter short genes
    filtered = {k: v for k, v in genes.items() if k not in short}

    # Get gene (chrom, start,end, gid) in a sorted list
    for gene, vals in filtered.items():
        chrom, start, end, gid = vals['chrom'], vals['start'], vals['end'], vals['gid']
        gcoords.append((chrom, int(start), int(end), gid))
        gnames.append(gid)

    # Get the intergenic (chrom,start,end in a list)
    for item in intergenic:
        chrom, start, end = item.split("\t")
        start, end = int(start) + 1, int(end)
        igcoords.append((chrom, start, end))

    # Sort gene list and intergenic list by chrom and start
    gcoords.sort(key=lambda x: (x[0], x[1]))
    igcoords.sort(key=lambda x: (x[0], x[1]))

    # Go through the intergenic list and check if its coordinates are between the neighboring gene starts.
    for i in igcoords:
        chrom, start, end = i

        for idx, g in enumerate(gcoords):

            if idx == len(gcoords) - 1:
                # Last one
                gchrom1, gstart1, gend1, gid1 = gcoords[idx - 1]
                gchrom2, gstart2, gend2, gid2 = g
                if chrom == gchrom1 and chrom == gchrom2 and start > gstart1 and end < gstart2:
                    name = ":".join([gid1, gid2])
                    ignames[name] = i
                break

            gchrom1, gstart1, gend1, gid1 = g
            gchrom2, gstart2, gend2, gid2 = gcoords[idx + 1]
            if chrom == gchrom1 and chrom == gchrom2 and start > gstart1 and end < gstart2:
                name = ":".join([gid1, gid2])
                ignames[name] = i
                break
    return ignames


def make_intergenic(ann, genome):
    """
    Create intergenic regions using bedtools.
    """
    try:
        cmd = f"cat {ann}| awk '$3==\"gene\" {{print}}' |bedtools sort -faidx {genome} -i - | bedtools complement -i - -g {genome}"
        res = subprocess.run(cmd, shell=True, stdout=subprocess.PIPE, check=True, universal_newlines=True)
    except:
        print("Error creating the intergenic intervals. Check if gene feature is present in annotation file.")
        sys.exit()

    res_list = res.stdout.split("\n")
    res_list = res_list[:-1]

    return res_list


def get_short_genes(genes):
    """
    Filter short genes that are completely inside another.
    """
    short = []

    # get short genes
    for gene1, val1 in genes.items():

        chrom1, start1, end1, gid1 = val1['chrom'], val1['start'], val1['end'], val1['gid']

        for gene2, val2 in genes.items():
            chrom2, start2, end2, gid2 = val2['chrom'], val2['start'], val2['end'], val2['gid']

            if chrom1 != chrom2:
                continue

            if start1 >= start2 and end1 <= end2 and gid1 != gid2:
                short.append(gid1)

    return short


def parse_gene(gene):
    return gene['chrom'], gene['start'], gene['end'], gene['strand'], gene['gtype'], gene['gid']


def make_feature_bed(genes, merged):
    """
    Makes a bed file of features.
    """
    bedfile = os.path.join(TMP, "features.bed")
    fh = open(bedfile, "w")

    for key, vals in genes.items():
        chrom, start, end, strand, gtype, gid = parse_gene(vals)

        # If there are no features for a gene, continue
        if not merged[gid]:
            continue

        coords = merged[gid]

        for c in coords:
            start, end = str(c[0] - 1), str(c[1])
            out = "\t".join([chrom, start, end, gid, ".", strand])
            fh.write(out + "\n")

    return bedfile


def get_depth(bam, bed, strand='reverse', flag=""):
    """
    Use bedtools to get the depth at every position.
    """

    def get_strand_flag(strand):
        if strand == "both":
            lib_strand = ""
        else:
            lib_strand = '-S' if strand == "reverse" else '-s'
        return lib_strand

    # Set strandedness.
    lib_strand = get_strand_flag(strand)

    # Output file.
    suffix = "depth.txt"
    outfile = get_filename(bam)
    outfile = outfile.replace("_primary", "") + f"_{flag}_{suffix}"
    outfile = os.path.join(TMP, outfile)

    # Command to get the depth.
    cmd2 = f'bedtools coverage -d -a {bed} -b {bam} {lib_strand} | cut -f 4,8 > {outfile}'
    # print(cmd2)

    d = subprocess.run(cmd2, shell=True, stdout=subprocess.PIPE, stderr=subprocess.PIPE, check=True,
                       universal_newlines=True)
    # Exit on error.
    exit_on_error(d.stderr) if d.stderr and d.returncode != 0 else None

    # os.system(cmd2)

    return outfile


def get_obs_tin(bam, bed, strand, bgfile="", size=50, flag=""):
    # Get depth for each position in each gene.
    depth_file = get_depth(bam, bed, strand, flag=flag)

    # Get tin.
    tin_store = get_tin(depth_file, bgfile, size)

    # Collect sample name.
    sample = get_filename(bam).replace("_primary", "")
    tin_store['samples'] = sample + "_obs_tin"
    return tin_store


def get_tin(depth_file, bgfile="", size=50):
    """
    Calculate tin from the depth file.
    Return effective_length, background_noise and tin_score.
    """

    covs, background, store = dict(), dict(), dict()

    curr, uid, path = "", "", ""

    if bgfile:
        background = store_background(bgfile)

    stream = csv.reader(open(depth_file), delimiter="\t")

    for row in stream:

        uid, depth = row[0], float(row[1])

        # The very first time
        if not curr:
            curr = uid

        if uid != curr:
            fid, tlen, bg, tin = process_tin(coverages=covs, background=background, uid=curr, size=size)
            store[fid] = (tlen, bg, tin)
            curr = uid
            covs = dict()

        covs.setdefault(uid, []).append(depth)

    # Print the last element
    fid, tlen, bg, tin = process_tin(coverages=covs, background=background, uid=uid, size=size)
    store[fid] = (tlen, bg, tin)

    return store


def process_tin(coverages, background, uid, size):
    """
    Calculates tin for the coverages and store in a dictionary
    where values are a tuple with(effective_len, background,tin-score)
    """

    vals = coverages[uid]

    # Get background.
    bg = float(background.get(uid, 0.0))

    # Subtract background.
    vals = subtract_background(vals, bg)

    # Make coverages float.
    vals = list(map(float, vals))

    # Calculate tin score.
    tin, tlen = tin_score(cvg=vals, size=size)

    tin = round(tin, 1)

    return uid, tlen, bg, tin


def shannon_entropy(vals):
    """
    calculate shannon's H = -sum(P*log(P)). arg is a list of float numbers. Note we used
    natural log here.
    """

    val_sum = sum(vals)
    entropy = 0.0

    for i in vals:
        entropy += (i / val_sum) * math.log(i / val_sum)

    return 0 if entropy == 0.0 else -entropy


def tin_score(cvg, size=50):
    """
    calcualte TIN score
    cvg : coverage at each base position as a list
    size : no. of bases to be omitted from the beginning and end of the transcript so as to get the effective length.
    returns transcript tin score and its effective length
    """

    # Get effective size
    # For short genes: if effective length <=0, then gene length is teh effective length

    cvg = cvg[size:-size] if (size != 0 and len(cvg) - size * 2 > 0) else cvg

    eff_len = len(cvg)

    # Change to float
    cvg = map(float, cvg)

    # Remove zeros
    cvg = list(filter(None, cvg))

    if not cvg:
        tin = 0.0
        return tin, eff_len

    # Calculate shannon's entropy
    ent = shannon_entropy(cvg)

    # Calculate uniformity
    uni = math.exp(ent)

    # Calculate tin on the effective length
    tin = 100 * (uni) / eff_len

    return tin, eff_len


def get_cov(count, read_len, size):
    return count * read_len / size


def get_expected_tin(cov):
    return (1 - math.exp(-cov)) * 100


def get_primary_aln(bam):
    """
    Extract primary alignments from bam.
    """

    pbam = get_filename(bam) + "_primary.bam"
    pbam = os.path.join(TMP, pbam)
    cmd1 = f'samtools view -h -b -F 4 -F 2304 {bam} >{pbam}'
    cmd2 = f'samtools index {pbam}'

    d1 = subprocess.run(cmd1, shell=True, stdout=subprocess.PIPE, stderr=subprocess.PIPE, check=True,
                        universal_newlines=True)
    d2 = subprocess.run(cmd2, shell=True, stdout=subprocess.PIPE, stderr=subprocess.PIPE, check=True,
                        universal_newlines=True)

    # Exit on error.
    exit_on_error(d1.stderr) if d1.stderr and d1.returncode != 0 else None
    exit_on_error(d2.stderr) if d2.stderr and d2.returncode != 0 else None
    return pbam


def store_background(fname):
    store = dict()

    stream = csv.reader(open(fname), delimiter="\t")
    for row in stream:
        tid, bg = row[:]
        store[tid] = bg
    return store


def subtract_background(elms, bg):
    def z(x):
        return 0 if x < 0 else x

    # Subtract background
    elms = [val - bg for val in elms]

    #  Make coverage to 0 if value < 0
    elms = list(map(z, elms))
    return elms


def get_sample_vals(info, sample_index):
    """
    Extract sample specific count from a master dictionary of lists.
    """
    sample_vals = dict()
    for key, vals in info.items():
        if key == "samples":
            continue
        sample_vals[key] = vals[sample_index]

    return sample_vals


def collect_measures(counts, tin, sense_counts={}, antisense_counts={}, sense_tin={},
                     antisense_tin={}, strand="both"):
    """
    Collect exp,tin, obs_tin, tin_sense, tin_antisense in a single dictionary.
    """
    gtins = dict()

    for gene, vals in counts.items():
        d = dict()
        if strand == "both":
            d['count'] = vals
            d['tin'] = tin[gene][2]

        else:
            d['count'] = vals
            d['sense_count'] = sense_counts[gene]
            d['antisense_count'] = antisense_counts[gene]
            d['tin'] = tin[gene][2]
            d['sense_tin'] = sense_tin[gene][2]
            d['antisense_tin'] = antisense_tin[gene][2]

        gtins[gene] = d

    return gtins


def get_neighbor_values(lgene, rgene, ileft, iright, gmeasures, igmeasures):
    """
    Collect the tin and count values of left gene, left  intergenic region, right gene and right intergenic region
    in unstranded mode.
    """

    ileft_tin = igmeasures[ileft]['tin'] if ileft in igmeasures else None
    iright_tin = igmeasures[iright]['tin'] if iright in igmeasures else None
    ileft_count = igmeasures[ileft]['count'] if ileft in igmeasures else None
    iright_count = igmeasures[iright]['count'] if iright in igmeasures else None

    gleft_tin = gmeasures[lgene]['tin']
    gright_tin = gmeasures[rgene]['tin']
    gleft_count = gmeasures[lgene]['count']
    gright_count = gmeasures[rgene]['count']

    ig_vals = (ileft_tin, iright_tin, ileft_count, iright_count)
    gene_vals = (gleft_tin, gright_tin, gleft_count, gright_count)

    return gene_vals, ig_vals


def get_effective_length(merged, size=50):
    """
    Get length as the sum of feature lengths.
    """

    flen = dict()

    def x(a):
        return (a[1] - a[0] + 1)

    for key, vals in merged.items():
        # size*2 since we subtract size bases from both ends while calculating tin.
        tot_len = sum(map(x, vals))
        eff_len = tot_len - size * 2
        eff_len = tot_len if eff_len <= 0 else eff_len
        flen[key] = eff_len
    # Header
    flen['samples'] = "eff_length"

    return flen


def get_exp_tin(counts, paired, read_len, feat_len):
    """
    Calculate expected tin from counts.
    Returns a dictionary with expected tins.
    """

    def remove_string(x):
        return x.replace("_count", "")

    store = dict()

    for fid, vals in counts.items():
        # Process sample names.
        if fid == "samples":
            vals = list(map(remove_string, vals))
            vals = add_string(vals, "_exp_tin")
            store.setdefault(fid, []).extend(vals)
            continue

        # Process counts.
        for v in vals:
            count = v
            count = float(count) * 2 if paired else float(count)
            flen = feat_len[fid]
            cov = get_cov(count, read_len, flen)
            exp_tin = round(get_expected_tin(cov), 2)
            store.setdefault(fid, []).append(exp_tin)

    return store


def parse_counts(counts):
    res = dict()
    for key, vals in counts.items():
        if key == "samples":
            res[key] = vals
            continue
        for v in vals:
            res.setdefault(key, []).append(v['count'])
    return res


def check_runin(data, gmeasures, igmeasures, strand, tin_cutoff=40, count_cutoff=40):
    """
    Check runins (overlap) for genes with tin <= tin_cutoff and count >=count_cutoff.
    Tin  and count values of the feature is checked with the tin and count values in the feature strand
    of the neighboring genes on both sides.
    """

    ileft_tin, gleft_tin, iright_tin, gright_tin = 0, 0, 0, 0
    ileft_count, gleft_count, iright_count, gright_count = 0, 0, 0, 0

    runins, params = dict(), dict()

    for gene, vals in gmeasures.items():

        gene_tin = vals['tin']
        gene_count = vals['count']

        # Checking for overlap only for genes with tin in range 1 to 40 and (count!=0 and count >count_cutoff).
        # ie, enough reads/expression but not good tin.
        if gene_tin > tin_cutoff or gene_tin == 0 or gene_count < count_cutoff:
            continue

        gene_strand = data[gene]['strand']
        lgene_strand = data[gene]['left_gene_strand']
        rgene_strand = data[gene]['right_gene_strand']
        lgene = data[gene]['left_gene']
        rgene = data[gene]['right_gene']

        # The very first, the very last and short genes.
        if lgene == "" or rgene == "":
            continue

        # Intergenic dict keys : left_right
        ileft = ":".join([lgene, gene])
        iright = ":".join([gene, rgene])

        # Unstranded mode check.
        if strand == "both":
            gene_vals, ig_vals = get_neighbor_values(lgene, rgene, ileft, iright, gmeasures, igmeasures)
            gleft_tin, gright_tin, gleft_count, gright_count = gene_vals
            ileft_tin, iright_tin, ileft_count, iright_count = ig_vals

        # Stranded mode check.
        else:
            if gene_strand == "+" and strand == "reverse":
                ileft_tin = igmeasures[ileft]['antisense_tin'] if ileft in igmeasures else None
                iright_tin = igmeasures[iright]['antisense_tin'] if iright in igmeasures else None
                ileft_count = igmeasures[ileft]['antisense_count'] if ileft in igmeasures else None
                iright_count = igmeasures[iright]['antisense_count'] if iright in igmeasures else None

                gleft_tin = gmeasures[lgene]['antisense_tin'] if lgene_strand == "+"  else gmeasures[lgene]['sense_tin']
                gright_tin = gmeasures[rgene]['antisense_tin'] if rgene_strand == "+" else gmeasures[rgene]['sense_tin']
                gleft_count = gmeasures[lgene]['antisense_count'] if lgene_strand == "+"  else gmeasures[lgene]['sense_count']
                gright_count = gmeasures[rgene]['antisense_count'] if rgene_strand == "+" else gmeasures[rgene]['sense_count']

            if gene_strand == "-" and strand == "reverse":
                ileft_tin = igmeasures[ileft]['sense_tin'] if ileft in igmeasures else None
                iright_tin = igmeasures[iright]['sense_tin'] if iright in igmeasures else None
                ileft_count = igmeasures[ileft]['sense_count'] if ileft in igmeasures else None
                iright_count = igmeasures[iright]['sense_count'] if iright in igmeasures else None

                gleft_tin = gmeasures[lgene]['antisense_tin'] if lgene_strand == "-" else gmeasures[lgene]['sense_tin']
                gright_tin = gmeasures[rgene]['antisense_tin'] if rgene_strand == "-" else gmeasures[rgene]['sense_tin']
                gleft_count = gmeasures[lgene]['antisense_count'] if lgene_strand == "-" else gmeasures[lgene]['sense_count']
                gright_count = gmeasures[rgene]['antisense_count'] if rgene_strand == "-" else gmeasures[rgene]['sense_count']

            if gene_strand == "+" and strand == "same":
                ileft_tin = igmeasures[ileft]['sense_tin'] if ileft in igmeasures else None
                iright_tin = igmeasures[iright]['sense_tin'] if iright in igmeasures else None
                ileft_count = igmeasures[ileft]['sense_count'] if ileft in igmeasures else None
                iright_count = igmeasures[iright]['sense_count'] if iright in igmeasures else None

                gleft_tin = gmeasures[lgene]['sense_tin'] if lgene_strand == "+" else gmeasures[lgene]['antisense_tin']
                gright_tin = gmeasures[rgene]['sense_tin'] if rgene_strand == "+" else gmeasures[rgene]['antisense_tin']
                gleft_count = gmeasures[lgene]['sense_count'] if lgene_strand == "+" else gmeasures[lgene]['antisense_count']
                gright_count = gmeasures[rgene]['sense_count'] if rgene_strand == "+" else gmeasures[rgene]['antisense_count']

            if gene_strand == "-" and strand == "same":
                ileft_tin = igmeasures[ileft]['antisense_tin'] if ileft in igmeasures else None
                iright_tin = igmeasures[iright]['antisense_tin'] if iright in igmeasures else None
                ileft_count = igmeasures[ileft]['antisense_count'] if ileft in igmeasures else None
                iright_count = igmeasures[iright]['antisense_count'] if iright in igmeasures else None

                gleft_tin = gmeasures[lgene]['sense_tin'] if lgene_strand == "-" else gmeasures[lgene]['antisense_tin']
                gright_tin = gmeasures[rgene]['sense_tin'] if rgene_strand == "-" else gmeasures[rgene]['antisense_tin']
                gleft_count = gmeasures[lgene]['sense_count'] if lgene_strand == "-" else gmeasures[lgene]['antisense_count']
                gright_count = gmeasures[rgene]['sense_count'] if rgene_strand == "-" else gmeasures[rgene]['antisense_count']

        # If the intergenic region is None, set the values to be equivalent to gene,
        # so that a general condition can be applied.

        ileft_tin = gene_tin if ileft_tin is None else ileft_tin
        iright_tin = gene_tin if iright_tin is None else iright_tin
        ileft_count = gene_count if ileft_count is None else ileft_count
        iright_count = gene_count if iright_count is None else iright_count

        if left_tin_condition(gene_tin, gleft_tin, ileft_tin) and \
                left_count_condition(gene_count, gleft_count, ileft_count) and \
                right_tin_condition(gene_tin, gright_tin, iright_tin) and \
                right_count_condition(gene_count, gright_count, iright_count):
            runins[gene] = "both"
        elif left_tin_condition(gene_tin, gleft_tin, ileft_tin) and \
                left_count_condition(gene_count, gleft_count, ileft_count):
            runins[gene] = "left"
        elif right_tin_condition(gene_tin, gright_tin, iright_tin) and \
                right_count_condition(gene_count, gright_count, iright_count):
            runins[gene] = "right"

    return runins


def left_tin_condition(gene_tin, gleft_tin, igleft_tin):
    return gleft_tin >= gene_tin and igleft_tin >= gene_tin


def right_tin_condition(gene_tin, gright_tin, igright_tin):
    return gright_tin >= gene_tin and igright_tin >= gene_tin


def left_count_condition(gene_count, gleft_count, igleft_count):
    return gleft_count >= gene_count and igleft_count >= gene_count


def right_count_condition(gene_count, gright_count, igright_count):
    return gright_count >= gene_count and igright_count >= gene_count


def exit_on_error(err):
    print(err)
    sys.exit()


def check_strand(strand):
    return True if strand in ['both', 'same', 'reverse'] else False


def check_libtype(libtype):
    return True if libtype in ['single', 'paired'] else False


def check_bam(files):
    """
    Check if all files have the extension .bam
    """
    extns = [os.path.splitext(os.path.basename(f))[1] for f in files]
    return len(set(extns)) == 1


def check_gtf(ann):
    """
    Check if annotation is a GTF file.
    """
    ext = os.path.splitext(os.path.basename(ann))[1]
    # get 3 characters from extension.
    ext = ext[:4]
    if ext not in [".GTF", ".gtf"]:
        return False
    return True


def check_inputs(args):
    # Make sure the alignment file extension is bam.
    if not check_bam(args.bams):
        print("All alignment files must be have '.bam' extension.")
        sys.exit()

    # Make sure entered strand is valid.
    if not check_strand(args.strand):
        print("Invalid strand. Available options are both, same, reverse")
        sys.exit()

    # Make sure entered libtype is valid.
    if not check_libtype(args.lib_type):
        print("Invalid lib_type. Available options are single or paired")
        sys.exit()

    if not args.ann or not check_gtf(args.ann):
        print("GTF annotation file with gene feature row is required.")
        sys.exit()

    return


def get_random_string(length):
    # choose from all lowercase letter
    letters = string.ascii_lowercase
    random_str = ''.join(random.choice(letters) for i in range(length))
    return random_str


def run(args):
    bams = args.bams
    ann = args.ann
    feat = args.feat
    size = args.size
    background = args.bg
    libtype = args.lib_type
    read_len = args.read_len
    strand = args.strand
    tin_cutoff = args.tin_cutoff
    count_cutoff = args.count_cutoff
    #tmp = args.tmp

    groupby = "gene"

    bg_file, intron_len, intron_gtf = None, None, None

    # Check if inputs are valid.
    check_inputs(args)

    # Generate a random temporary folder for intermediate files.
    global TMP
    TMP = "tmp_" + get_random_string(3)
    os.system(f'mkdir -p {TMP}')

    # Create run-log.
    log = os.path.join(TMP, "runlog.txt")

    # Make a string of bam files.
    bam = " ".join(bams)

    # Paired or single.
    paired = True if libtype == "paired" else False

    # Extract features from annotation file.
    genes, features = collect_features(ann=ann, feat_type=feat)

    # Get genes that are completely inside another.
    short = get_short_genes(genes)

    # Add neighbors
    genes = get_neighbors(genes=genes, short=short)

    # Merge overlapping features by groupby attribute.
    merged = merge_features(features)

    # Make a bed file of merged features.
    bed = make_feature_bed(genes, merged)

    # Generate genome file.
    genomefile = bam_to_genome(bam)

    # Get intergenic regions
    interg_coords = make_intergenic(ann=ann, genome=genomefile)

    # Name intergenic regions
    intergenic = name_intergenic_regions(interg_coords, genes, short)

    # Make intergenic bed and gff files.
    ig_bed, ig_ann = make_intergenic_files(intergenic)

    # Make intergenic annotation file.
    ig_ann = make_intergenic_gtf(intergenic)

    # Get intron coordinates for background noise calculation.
    if background:
        introns = get_intron_coords(merged)
        intron_len = get_effective_length(introns, size=0)
        intron_gtf = make_intron_gtf(genes, introns)

    # Get gene counts in both sense and antisense.
    gcounts_sense = get_counts(bam=bam, ann=ann, strand="same", feat=feat,
                               groupby=groupby, paired=paired, flag="gene_sense", log=log)

    gcounts_antisense = get_counts(bam=bam, ann=ann, strand="reverse", feat=feat,
                                   groupby=groupby, paired=paired, flag="gene_antisense", log=log)

    # Get intergenic counts in both sense and antisense.
    igcounts_sense = get_counts(bam=bam, ann=ig_ann, strand="same", feat="intergenic",
                                groupby=groupby, paired=paired, flag="ig_sense", log=log)

    igcounts_antisense = get_counts(bam=bam, ann=ig_ann, strand="reverse", feat="intergenic",
                                    groupby=groupby, paired=paired, flag="ig_antisense", log=log)

    # Get the observed gene counts and intergenic according to the library strand.
    if strand == "both":
        # Gene counts in unstranded mode. Sum up both sense and antisense counts.
        gene_counts = combine_counts(gcounts_sense, gcounts_antisense)
        ig_counts = combine_counts(igcounts_sense, igcounts_antisense)
    else:
        gene_counts = gcounts_antisense if strand == "reverse" else gcounts_sense
        ig_counts = igcounts_antisense if strand == "reverse" else igcounts_sense

    # Get feature length
    feat_len = get_effective_length(merged, size)

    # Calculate expected tin.
    exp_tins = get_exp_tin(counts=gene_counts, paired=paired, read_len=read_len, feat_len=feat_len)

    # Sample specific TIN calculations begins.
    runins, obs_tins = dict(), dict()

    for idx, bam in enumerate(bams):

        # Parse out the gene counts and intergenic counts for the current bam.
        sample_gene_counts = get_sample_vals(info=gene_counts, sample_index=idx)
        sample_ig_counts = get_sample_vals(info=ig_counts, sample_index=idx)

        sample_gcounts_sense = get_sample_vals(info=gcounts_sense, sample_index=idx)
        sample_gcounts_antisense = get_sample_vals(info=gcounts_antisense, sample_index=idx)

        sample_ig_counts_sense = get_sample_vals(info=igcounts_sense, sample_index=idx)
        sample_ig_counts_antisense = get_sample_vals(info=igcounts_antisense, sample_index=idx)

        # Calculate background noise if "--bg" is specified.
        if background:
            bg_file = get_background_noise(bam=bam, strand=strand,
                                           intron_ann=intron_gtf, intron_len=intron_len, groupby=groupby, paired=paired)

        # Extract primary alignments
        pbam = get_primary_aln(bam)

        if strand == "both":
            # Get gene tin.
            gene_tin = get_obs_tin(bam=pbam, bed=bed, strand=strand, bgfile=bg_file, size=size, flag="gene")

            # Get intergenic tin.
            ig_tin = get_obs_tin(bam=pbam, bed=ig_bed, strand=strand, size=0, flag="ig")

            # Get the gene counts and tins calculated for the sample in a single place.
            gene_measures = collect_measures(counts=sample_gene_counts, tin=gene_tin,
                                             strand=strand)

            # Get intergenic counts and tins for the sample single place.
            intergenic_measures = collect_measures(counts=sample_ig_counts, tin=ig_tin, strand=strand)

        else:

            # Get gene sense and antisense tin.
            gtin_sense = get_obs_tin(bam=pbam, bed=bed, strand="same", bgfile=bg_file, size=size, flag="gene_sense")
            gtin_antisense = get_obs_tin(bam=pbam, bed=bed, strand="reverse", bgfile=bg_file, size=size,
                                         flag="gene_antisense")

            # Get intergenic sense and antisense tin. Intergenic bed has + as sense strand.
            igtin_sense = get_obs_tin(bam=pbam, bed=ig_bed, strand="same", size=0, flag="ig_sense")
            igtin_antisense = get_obs_tin(bam=pbam, bed=ig_bed, strand="reverse", size=0, flag="ig_antisense")

            # Get tin according to the library strand.
            gene_tin = gtin_antisense if strand == "reverse" else gtin_sense
            ig_tin = igtin_antisense if strand == "reverse" else igtin_sense

            # Get the counts and the different types of tins calculated for a gene in a single place.
            gene_measures = collect_measures(counts=sample_gene_counts, tin=gene_tin,
                                             sense_counts=sample_gcounts_sense,
                                             antisense_counts=sample_gcounts_antisense,
                                             sense_tin=gtin_sense, antisense_tin=gtin_antisense, strand=strand)

            # Get intergenic counts and tins in a single place.
            intergenic_measures = collect_measures(counts=sample_ig_counts, tin=ig_tin,
                                                   sense_counts=sample_ig_counts_sense,
                                                   antisense_counts=sample_ig_counts_antisense,
                                                   sense_tin=igtin_sense,
                                                   antisense_tin=igtin_antisense, strand=strand)

        # Check overlap
        overlaps = check_runin(data=genes, gmeasures=gene_measures, igmeasures=intergenic_measures,
                               strand=strand, tin_cutoff=tin_cutoff, count_cutoff=count_cutoff)

        # Collect obs_tins.
        for uid, vals in gene_tin.items():
            if uid == "samples":
                obs_tins.setdefault(uid, []).append(vals)
            else:
                obs_tins.setdefault(uid, []).append(vals[2])

        # Collect runins
        for gene in gene_counts:
            if gene == "samples":
                continue
            overlap = overlaps[gene] if gene in overlaps else ""
            runins.setdefault(gene, []).append(overlap)
        runins.setdefault('samples', []).append(get_filename(bam) + "_overlap")

    # Collect all results, gid,counts, exp_tin, obs_tin
    results = dict()

    for k, v in gene_counts.items():
        vals = list()
        vals.append(feat_len[k])
        vals.extend(v)
        vals.extend(exp_tins[k])
        vals.extend(obs_tins[k])
        vals.extend(runins[k])
        results[k] = vals

    # Print results
    for gene, vals in results.items():
        vals = map(str, vals)
        vals = "\t".join(vals)
        out = "\t".join([gene, vals])
        print(out)

    # Clean up temporary files.
    # cmd = f'rm -f {TMP}/*primary.bam*'
    cmd = f'rm -rf {TMP}'
    os.system(cmd)


if __name__ == "__main__":
    parser = argparse.ArgumentParser(description="""Calculates transcript overlaps from neighboring genes based on
                                                    gene reads counts and gene tins.""")

    parser.add_argument("bams", nargs='+')

    parser.add_argument('--ann', dest='ann', type=str, required=True,
                        help='GTF file. ')

    parser.add_argument('--feat', dest='feat', type=str, required=False, default="exon",
                        help="""Feature on which TIN needs to be calculated for a gene. It should match with the 3rd column of the annotation file. 
                        Default is exon.""")

    parser.add_argument('--strand', dest='strand', type=str, required=False, default="both",
                        help="""Strand on which tin should be calculated. Default is both (un-stranded).
                             Possible values are both, same, reverse""")

    parser.add_argument('--lib_type', dest='lib_type', type=str, required=False, default='single',
                        help="""Specify if the library is paired or single. 
                        Valid options are 'single' or 'paired. Default is single.'""")

    parser.add_argument('--read_len', dest='read_len', type=int, required=False, default=100,
                        help="""Read length. Default is 100 bp'""")

    parser.add_argument('--bg', dest='bg', required=False, action='store_true', default=False,
                        help="""When specified background noise will be subtracted from tin calculations.
                              """)

    parser.add_argument('--n', dest='size', type=int, required=False, default=50,
                        help="""No. of bases to be subtracted from each ends of the feature to calculate effective length.
                                Default is 50.
                                """)

    parser.add_argument('--tin_cutoff', dest='tin_cutoff', type=int, required=False, default=40,
                        help="""Tin cutoff to be used for checking overlaps. Genes with tin < tin-cutoff and
                         count >count_cutoff are checked for overlaps. Default is 40.
                        """)
    parser.add_argument('--count_cutoff', dest='count_cutoff', type=int, required=False, default=40,
                        help="""Count cutoff to be used for checking overlaps. ie, minimum reads required to consider a gene as being expressed. 
                        Genes with count > count-cutoff and tin <tin-cutoff are checked for overlaps. Default is 40.
                        """)

    # parser.add_argument('--tmp', dest='tmp', type=str, required=False,
    #                     help="""Folder to store intermediate files. By default a random folder will be generated.""")

    args = parser.parse_args()

    run(args)
```
