## Supplementary material for "The *Plasmodium* NOT1-G Paralogue Acts as an Essential Nexus for Sexual Stage Maturation and Parasite Transmission": Supp File 3 - RNA-seq bioinformatics scripts and MakeFile: README.txt

Scripts used in the analysis.

1. Makefile - Makefile with all commands used in the bioinformatics analysis.
2. transcript_tin.py - Custom python script used to calculaute transcript TIN values.
3. make_gene_tin.py - Custom python script used to compute gene-level TIN from transcript TINs.
4. overlap.py - Custom python script used to flag potential transcript overlaps.
5. merge.py - Custom python script used to merge sample tin files into a single file.
6. deseq2.r - Custom R script used for DESeq2 differential expression analysis and the generation of MA plots.
7. heatmap.r - Custom R script used to generate heatmaps.
8. volcanoplot.r - Custom R script used to generate volcano plots.
9. summary_plots.r - Custom R script used to generate pca plot and sample-to-sample distance heatmap.
