## Supplementary material for "The *Plasmodium* NOT1-G Paralogue Acts as an Essential Nexus for Sexual Stage Maturation and Parasite Transmission": Supp File 3 - RNA-seq bioinformatics scripts and MakeFile: summary_plots.r

```
#!/usr/bin/env Rscript
#
# This script uses function in DESeq2 package and  generates pca plot and sample-to-sample distance heatmap of all samples.
#
# Usage:
# cat counts.txt | Rscript summary_plots.r
#

# Set the plot dimensions.
WIDTH = 6
HEIGHT = 5

# Input stream.
infile = "stdin"

# Load the library while suppressing verbose messages.
suppressPackageStartupMessages(library(DESeq2))

reps1 = 6
reps2 = 6

# Set up the conditions based on the experimental setup.
asexual_WT= rep("WT_asexual", 3)
sexual_WT = rep("WT_sexual", 3)
cond1 = c(asexual_WT,sexual_WT)

asexual_KO =rep("KO_asexual", 3)
sexual_KO = rep("KO_sexual", 3)
cond2 = c(asexual_KO,sexual_KO)

# Read the data from the standard input.
counts = read.table(infile, header=TRUE, sep="\t", row.names=1 )

# Assume the last N + M columns are the count matrix.
idx = ncol(counts) - (reps1 + reps2)

# Cut out the valid columns.
#counts = counts[-c(1:idx)]
if (idx > 0) counts = counts[-c(1:idx)] else counts=counts

# Some tools generate the estimated counts as real numbers
# DESeq 2 allows only integers. We need to convert real numbers to rounded integers.
numeric_idx = sapply(counts, mode) == 'numeric'
counts[numeric_idx] = round(counts[numeric_idx], 0)

# Build the dataset
samples = names(counts)
condition = factor(c(cond1, cond2))
colData = data.frame(samples=samples, condition=condition)

# Create DESEq2 dataset.
dds = DESeqDataSetFromMatrix(countData=counts, colData=colData, design = ~condition)

# Variance Stabilizing Transformation.
vsd = vst(dds)

# Open the drawing device.
pdf('pca.pdf', width = WIDTH, height = HEIGHT)

plotPCA(vsd, intgroup=c("condition"))

dev.off()

library(pheatmap)
library("RColorBrewer")

sampleDists = dist(t(assay(vsd)))
sampleDistMatrix = as.matrix(sampleDists)

rownames(sampleDistMatrix) = vsd$condition
colnames(sampleDistMatrix) = NULL

colors = colorRampPalette( rev(brewer.pal(9, "Blues")) )(255)

# Open the drawing device.

pdf('heatmap.pdf', width = 8, height = HEIGHT)

pheatmap(sampleDistMatrix,
         clustering_distance_rows=sampleDists,
         clustering_distance_cols=sampleDists,
         col=colors)

dev.off()
```
