## Supplementary material for "The *Plasmodium* NOT1-G Paralogue Acts as an Essential Nexus for Sexual Stage Maturation and Parasite Transmission": Supp File 3 - RNA-seq bioinformatics scripts and MakeFile: transcript_tin.py

```
import csv, math, sys
import argparse
import os

"""
This script calculate Transcript Integrity Number using Shannon's Entropy formula.
Input is a tab separated file of the format 'transcript-name\tpos\tdepth'
The file should include all positions including positions with 0 coverage. (ie, entire length).
"""


def shannon_entropy(vals):
    """
    calculate shannon's H = -sum(P*log(P)). arg is a list of float numbers. Note we used
    natural log here.
    """

    val_sum = sum(vals)
    entropy = 0.0

    for i in vals:
        entropy += (i / val_sum) * math.log(i / val_sum)

    return 0 if entropy == 0.0 else -entropy


def tin_score(cvg, size=50):
    """
    calcualte TIN score
    cvg : coverage at each base position as a list
    size : no. of bases to be omitted from the beginning and end of the transcript so as to get the effective length.
    returns transcript tin score and its effective length
    """

    # Get effective size
    cvg = cvg[size:-size]

    eff_len = len(cvg)

    # Change to float
    # cvg = map(float, cvg)

    # Remove zeros
    cvg = list(filter(None, cvg))

    if not cvg:
        tin = 0.0
        return tin, eff_len

    # Calculate shannon's entropy
    ent = shannon_entropy(cvg)

    # Calculate uniformity
    uni = math.exp(ent)

    # Calculate tin on the effective length
    tin = 100 * (uni) / eff_len

    return tin, eff_len


def store_background(fname):
    store = dict()

    stream = csv.reader(open(fname), delimiter="\t")
    for row in stream:
        tid, bg = row[:]
        store[tid] = bg
    return store


def subtract_background(elms, bg):
    def z(x):
        return 0 if x < 0 else x

    # Subtract background
    elms = [val - bg for val in elms]

    #  Make coverage to 0 if value < 0
    elms = list(map(z, elms))

    return elms


def print_last_elm(coverages, background, uid, size):
    vals = coverages[uid]

    #  Get background
    bg = float(background[uid]) if background else 0

    # Subtract background
    vals = subtract_background(vals, bg)

    # Make coverages float
    vals = list(map(float, vals))

    # Calculate tin score
    tin, tlen = tin_score(cvg=vals, size=size)
    tin = round(tin, 1)
    print("\t".join([uid, str(tlen), str(bg), str(tin)]))
    return tin


def get_tin(covfile, bgfile="", size=50):
    covs, background, store = dict(), dict(), dict()

    curr, uid, path = "", "", ""

    header = ["tid", "eff-length", "background", "tin-score"]
    print("\t".join(header))

    if bgfile:
        background = store_background(bgfile)

    stream = csv.reader(open(covfile), delimiter="\t")

    for row in stream:
        uid, depth = row[0], float(row[2])

        # The very first time
        if not curr:
            curr = uid

        if uid != curr:
            vals = covs[curr]

            # Get background noise
            bg = float(background[curr]) if background else 0

            # Subtract background
            vals = subtract_background(vals, bg)

            # Calculate tin score
            tin, tlen = tin_score(cvg=vals, size=size)
            tin = round(tin, 1)

            print("\t".join([curr, str(tlen), str(bg), str(tin)]))

            store['name'] = curr
            store['tin'] = tin
            curr = uid
            covs = dict()

        covs.setdefault(uid, []).append(depth)

    # Print the last element
    tin = print_last_elm(coverages=covs, background=background, uid=uid, size=size)
    store['name'] = uid
    store['tin'] = tin

    return store


def main():
    parser = argparse.ArgumentParser(description='''Calculates TIN score from a depth file.''')

    parser.add_argument('--cov', dest='cov_file', type=str, required=True,
                        help='Tab delimited file with coverages at every base position for each transcript.')

    parser.add_argument('--bg', dest='bg_file', type=str, required=False,
                        help="""Tab delimited file with background noises for each transcript.
                             """)

    parser.add_argument('--n', dest='size', type=int, required=False, default=50,
                        help="""No. of bases to be subtracted from the ends of transcript to calculate effective length.
                                Default is 50.
                                """)

    args = parser.parse_args()

    cov_file = args.cov_file
    bg_file = args.bg_file
    size = args.size

    get_tin(covfile=cov_file, bgfile=bg_file, size=size)


if __name__ == "__main__":
    main()
```
