## Supplementary material for "The *Plasmodium* NOT1-G Paralogue Acts as an Essential Nexus for Sexual Stage Maturation and Parasite Transmission": Supp File 3 - RNA-seq bioinformatics scripts and MakeFile: volcanoplot.r

```
#!/usr/bin/env Rscript

# This program creates volcano plot from differential expression result.

# Usage:
#
# cat results.csv | ./volcanoplot.r 'Plot Title'  >volcanoplot.pdf

suppressPackageStartupMessages(library(EnhancedVolcano))

# Command line argument.
args = commandArgs(trailingOnly=TRUE)  

# Input stream.
fname = "stdin"

# The first argument to the script.
title = args[1]
print(title)
if (is.na(title)) title = "" else title=title

# Set the plot dimensions.
WIDTH = 10
HEIGHT = 5

# Read data
res = read.csv(fname, header=TRUE, sep=",", row.names=1 )

# create custom key-value pairs for 'high', 'low', 'mid' expression by fold-change
# this can be achieved with nested ifelse statements

# create custom key-value pairs for 'high', 'low', 'mid' expression by fold-change
# this can be achieved with nested ifelse statements

keyvals <- ifelse( res$log2FoldChange < -2.5, 'dark green', 
              ifelse(res$log2FoldChange > -2.5 & res$log2FoldChange < -1, 'light blue',
                ifelse(res$log2FoldChange > 1 & res$log2FoldChange < 2.5, 'pink',
                    ifelse(res$log2FoldChange > 2.5, 'dark red',
                       'black'))))


keyvals[is.na(keyvals)] <- 'black'

names(keyvals)[keyvals == 'dark green'] <- 'log(2) FC< -2.5'
names(keyvals)[keyvals == 'black'] <- 'log(2) FC between -1 and +1'
names(keyvals)[keyvals == 'light blue'] <- 'log(2) FC between -1 and -2.5'
names(keyvals)[keyvals == 'pink'] <- 'log(2) FC between +1 and +2.5'
names(keyvals)[keyvals == 'dark red'] <- 'log(2) FC > +2.5'

# Open the drawing device.
pdf('|cat', width = WIDTH, height = HEIGHT)

# Draw the plot.
EnhancedVolcano(res,
                lab = rownames(res),
                x = 'log2FoldChange',
                y = 'FDR',
                xlim = c(-12,12),
                pCutoff = 0.05,
                FCcutoff = 2.5,
                #cutoffLineType = "blank", 
                colCustom = keyvals, colAlpha = 0.75, 
                gridlines.major=FALSE,gridlines.minor = FALSE,
                labSize = 0, legendPosition = "right",
                legendLabSize = 10,legendIconSize = 2,
                title =  title, subtitle = "",
                ylab= bquote(~-Log[10]~"FDR")
                
)

dev.off()
```
